# Synaptic endocytic failure drives dopamine deficiency and protein transfer-mediated striatal dopamine-like neuron compensation

**DOI:** 10.1101/2025.05.14.653914

**Authors:** Zhaoyan Li, Youneng Lin, Ana Caulino-Rocha, Bandhan Mukherjee, Man Liu, Xiaoxing Cao, Zhen Wang, Chuying Lai, Hua Huang, Zheng Ding, Ira Milosevic, Mian Cao

**Affiliations:** Programme in Neuroscience and Behavioural Disorders, Duke-NUS Medical School, Singapore; Multidisciplinary Institute of Aging, Centre for Innovative Biomedicine and Biotechnology, University of Coimbra, Coimbra, Portugal; Centre for Human Genetics, Nuffield Department of Medicine, University of Oxford, Oxford, UK; Department of Physiology, National University of Singapore, Singapore; Electrophysiology Core Facility, Yong Loo Lin School of Medicine, National University of Singapore, Singapore 117544, Singapore

**Keywords:** Synaptojanin 1, presynaptic endocytic failure, dopamine deficiency, dopamine-like neurons, inter-neuronal protein transfer, striatal compensation, Parkinson’s disease

## Abstract

Presynaptic endocytic dysfunction is an early feature of Parkinson’s disease (PD), but how dopamine circuits adapt to chronic synaptic failure remains unclear. In dopamine neuron-specific Synaptojanin 1 knockout mice, loss of this PD-linked endocytic protein causes terminal dystrophy and striatal dopamine deficiency, yet motor coordination is preserved. Chronic endocytic failure induces striatal dopamine-like neurons (DALNs) with circuit-dependent dopaminergic protein accumulation. DALNs do not arise through transcriptional reprogramming; instead, they acquire dopaminergic proteins from midbrain dopamine neurons through selective long-range transfer that favors soluble cargoes and can convey disease-associated proteins, including α-synuclein and tau. Functionally, DALNs support local dopamine production, partially preserve motor performance and increase resistance to acute dopaminergic lesions. This adaptation is shared across PD-linked endocytic mutants and is developmentally constrained, but is not reproduced by acute denervation. Thus, synaptic endocytic stress triggers inter-neuronal protein transfer as an adaptive circuit mechanism supporting dopamine compensation and neuroprotection in early PD.

## Introduction

Parkinson’s disease (PD) is characterized by progressive loss of dopamine neurons (DANs) in the substantia nigra (SNc), leading to striatal dopamine (DA) deficiency and motor impairment. However, overt motor symptoms typically emerge only after substantial disruption of dopaminergic innervation. Increasing evidence suggests that synaptic dysfunction precedes neuronal loss and represents an early event in PD pathogenesis. This prolonged pre-symptomatic phase implies that dopaminergic circuits can maintain function despite ongoing synaptic stress, but the underlying adaptive mechanisms remain poorly understood^1,2^.

Genetic studies strongly implicate presynaptic membrane trafficking in PD. More than 20 PD risk genes and over 90 genomic risk loci have been identified, several of which encode proteins involved in endocytic trafficking and synaptic vesicle recycling^3^. A major synaptic vesicle recycling pathway is clathrin-mediated endocytosis, in which the lipid phosphatase Synaptojanin 1 (SJ1) has a central role^4^. Together with its interactor Endophilin, SJ1 promotes clathrin uncoating by dephosphorylating PI(4,5)P_2_ and facilitating release of clathrin adaptors from newly formed vesicles^5–7^. Disruption of this machinery causes clathrin-coated vesicle accumulation and severe synaptic defects across model organisms^5–8^.

The relevance of SJ1 to PD is supported by human genetics. A homozygous R258Q mutation in SJ1, which impairs the Sac phosphatase domain, causes early-onset Parkinsonism and established SJ1 as PARK20^9,10^. Additional homozygous and compound-heterozygous SJ1 mutations have been associated with Parkinsonism, seizures and neurodegeneration^11,12^. Disease severity broadly correlates with the extent of SJ1 loss of function: truncating mutations cause severe neurodegeneration, epilepsy and early mortality, whereas missense mutations often produce milder phenotypes including Parkinsonism and seizure susceptibility. Rare heterozygous variants, eQTL associations and reduced SJ1 expression in PD tissues further suggest that impaired SJ1 function may contribute to both familial and sporadic PD^13–15^.

Mouse models mirror this genotype–phenotype relationship. Constitutive SJ1 knockout mice die shortly after birth with severe epilepsy^16,17^, whereas aged SJ1-haploinsufficient mice develop α-synuclein accumulation, autophagic and lysosomal abnormalities, and dopaminergic terminal degeneration^18–20^. Our previously generated SJ1 R258Q knock-in mice exhibit early-onset neurological abnormalities, generalized synaptic endocytic defects and selective degeneration of nigrostriatal dopaminergic terminals^21^. SJ1 also interacts genetically with other PD-associated endocytic proteins. Auxilin (Aux) (PARK19) knockout mice develop similar dopaminergic terminal abnormalities, and combining auxilin loss with SJ1 R258Q markedly exacerbates synaptic endocytic defects, striatal dystrophy, and mortality^22^. Loss of the PD-associated PI4P phosphatase Sac2 similarly worsens SJ1-related phenotypes^23^. Together, these studies implicate defective presynaptic membrane recycling as a convergent mechanism of dopaminergic vulnerability.

However, constitutive whole-body mutants cannot address the cell-autonomous requirement for SJ1 in DANs or reveal how striatal circuits adapt to chronic dopaminergic synaptic stress. Here, we generated DAN-specific SJ1 conditional knockout mouse models to define primary DAN defects and secondary striatal adaptations. Complete SJ1 loss caused severe dopaminergic terminal dystrophy, impaired DA transmission and striatal DA deficiency, yet motor coordination remained largely preserved. Unexpectedly, chronic endocytic failure induced striatal dopamine-like neurons (DALNs) with circuit-dependent dopaminergic features and reduced GABAergic excitability. DALNs acquired these features through a previously unrecognized long-range, selective protein transfer pathway from midbrain DANs, supporting local DA compensation and circuit-level protection. This adaptation was shared across PD-linked endocytic mutant mice, but absent from acute DA lesion models and limited after adult SJ1 deletion. Together, these findings reveal selective inter-neuronal protein transfer as an adaptive circuit response to chronic presynaptic endocytic stress in early PD.

## Results

### DAN-specific loss of SJ1 causes striatal terminal dystrophy and DA deficiency

To define the cell-autonomous role of SJ1 in DANs, we generated SJ1 floxed mice by inserting loxP sites flanking exon 4 of *Synj1* gene using CRISPR/Cas9 and crossed them with DAT-IRES-Cre mice to obtain DAT-Cre;SJ1 flx/flx mice, hereafter referred to as SJ1 cKO^DAT^ (Fig. S1A). Immunostaining confirmed efficient SJ1 depletion in tyrosine hydroxylase (TH)-positive DANs in the SNc and ventral tegmental area (VTA), while SJ1 expression was preserved in TH-negative midbrain neurons and non-dopaminergic regions such as the hippocampus (Fig. 1A, B; Fig. S1B). SJ1 cKO^DAT^ mice survived into adulthood without seizures, consistent with the idea that the epilepsy observed in constitutive SJ1 mutants reflects SJ1 loss outside the dopaminergic system.

**Figure 1.**
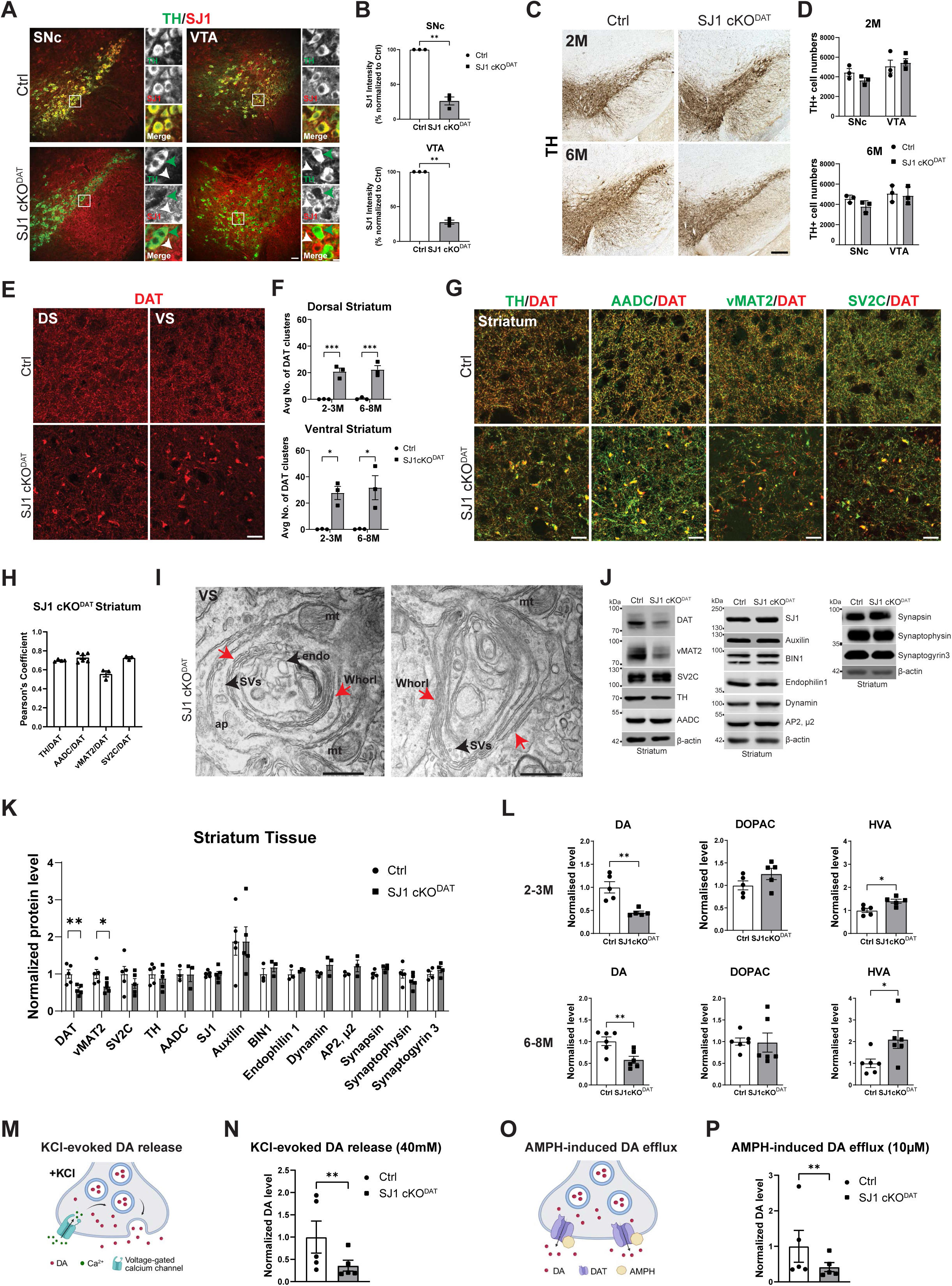
SJ1 conditional KO (SJ1 cKO^DAT^) in dopamine neurons (DANs) exhibit dystrophic dopaminergic axon terminals and dysregulated dopaminergic system in the striatum (A) Double staining of TH and SJ1 to show SJ1 specifically KO in midbrain dopamine neurons (DANs). Green arrows: TH-positive DANs; White arrows: TH-negative neurons. Scale bar: 20 µm. Inset: 10 µm. (B) Immunofluorescence analysis reveals significant reduction of SJ1 intensity in both SNc and VTA of SJ1 cKO^DAT^ compared to controls, confirming specific knockout in DANs. Mean fluorescence intensity of SJ1 was quantified and normalized to the corresponding control within each experimental batch, with control expression set to 100%. n=3 mice per genotype. (C), (D) Stereological analysis of TH-positive DANs in the SNc and VTA region of Ctrl and SJ1 cKO^DAT^ mice at 2-3 months and 6-8 months age group (C). There is no significant loss of DANs in both ages (D). The estimated cell numbers in one hemisphere are shown. n=3 mice per genotype. Scale bar: 200 µm. (E) DAT staining of 2 months old Ctrl and SJ1 cKO^DAT^ shows clustering phenotype in dorsal striatum (DS) and ventral striatum (VS). Scale bar: 20 µm. (F) Quantification for DAT clusters at 2-3 months and 6-8 months of age. The average number of clusters is counted in five randomly selected, 300 × 300-µm regions of interest (ROIs). n=3 mice per genotype. (G) Double staining of DAT with TH, AADC, vMAT2 and SV2C showed co-clustering in the dorsal striatum of SJ1 cKO^DAT^ mice. Scale bar: 20 µm. (H) Pearson’s correlation coefficient for striatum double staining shown in (G) reveals strong linear relationship between DAT with TH, AADC, vMAT2 and SV2C. (I) Representative EM micrographs of nerve terminals from the ventral striatum (VS) of SJ1 cKO^DAT^. Red arrows: Whorl-like multilayered membranes; Synaptic vesicles (SVs), endosomes (endo), autophagosome (AP) and mitochondria (mt) were also labelled in the images. Scale bar: 500 nm. (J), (K) Representative western blot and analysis for DA markers, endocytic and general synaptic markers from 2-3 months old Ctrl and SJ1 cKO^DAT^ half brain striatal homogenates. DAT and vMAT2 in SJ1 cKO^DAT^ exhibited significant reduction after normalization. n=3-5 mice per genotype. (L) HPLC analysis of total DA and its metabolites’ content in the striatum of SJ1 cKO^DAT^ mice showed an average of 50% reduction in striatal DA level, enhanced HVA level and non-significant change in DOPAC level for 2-3 months old and 6-8 months old. n number used for each genotype=5 for 2-3M; n=6 for 6-8M. (M) Illustration of KCl-evoked DA release in dopaminergic terminals. (N) Quantification of normalized DA concentration from KCl-evoked DA release in SJ1 cKO^DAT^ acute striatal slices (age range 2-9M) revealed significant reduction, suggesting impaired vesicular release. n=5 mice per genotype. (O) Illustration of AMPH-induced DA efflux in dopaminergic terminals. (P) Quantification of normalized DA concentration from AMPH-induced DA efflux in SJ1 cKO^DAT^ acute striatal slices (age range 2-9M) revealed significant reduction, suggesting reduced DAT-mediated DA efflux. n=5 mice per genotype. Data are represented as mean ± SEM. Statistical analysis: Welch’s unpaired t-test (B), multiple comparison with Holm-Šídák correction (D), paired t-test for each protein (K, L), ordinary one-way ANOVA with post-hoc Tukey’s test (F); *p<0.05, **p<0.01 and ***p<0.001.

Although TH-positive DAN somata were largely preserved in the SNc and VTA at 2–3 and 6–8 months of age (Fig. 1C, D), the striatum showed prominent DAT-positive dystrophic clusters in SJ1 cKO^DAT^ mice but not in controls (Fig. 1E, F). These clusters emerged by 3 weeks, increased in size and number, and peaked at 2–3 months (Fig. S1C). In contrast to SJ1^RQ^-KI mice, in which pathology is largely restricted to the dorsolateral striatum, SJ1 cKO^DAT^ mice showed widespread dystrophy across dorsal and ventral striatum, including the nucleus accumbens (NAc) core, lateral shell and olfactory tubercle (OT), with the strongest pathology in the NAc core (Fig. S1D–F). However, several VTA-derived projections, including those to the NAc medial shell, prefrontal cortex and basolateral amygdala, were relatively spared (Fig. S1G–I). Because these regions show low or undetectable DAT expression, DAT-enriched dopaminergic terminals may be particularly vulnerable to SJ1 loss. Thus, complete SJ1 deletion in DANs causes broad but projection-selective dopaminergic terminal dystrophy in a cell-autonomous and gene dosage-dependent manner.

The dystrophic clusters contained multiple dopaminergic terminal markers, including TH, aromatic L-amino acid decarboxylase (AADC), vesicular monoamine transporter 2 (vMAT2), and synaptic vesicle glycoprotein 2C (SV2C) (Fig. 1G, H; Fig. S2A), and partially colocalized with the presynaptic marker synaptotagmin-1 (Fig. S2B). They did not colocalize with D1R, D2R dopamine receptors or the medium spiny neuron marker Darpp32, confirming their dopaminergic terminal origin (Fig. S2C–F). No obvious astrocytic or microglial activation was detected (Fig. S3A–C). Although DAT-positive clusters partially colocalized with α-synuclein, pathological pSer129-α-synuclein aggregation was not detected in the striatum or midbrain (Fig. S3D–G).

Dystrophic clusters were negative for canonical clathrin-associated proteins, including clathrin light chain, endophilin 1 and amphiphysin 1 (Fig. S4A–E). Consistently, electron microscopy revealed multilayered, whorl-like membrane structures similar to those previously observed in SJ1^RQ^-KI and Aux-KO mice (Fig. 1I). These structures correlate with dopaminergic marker protein clusters and reflect abnormal plasma membrane invaginations at dopaminergic terminals. They contained synaptic vesicles, mitochondria, autophagosomes and endo/lysosomes (Fig. 1I; Fig. S5A, B), but did not show clathrin-coated vesicle accumulation. Thus, SJ1 loss in DAN terminals produces abnormal membrane remodeling rather than classical CCV buildup.

We next asked whether this terminal pathology compromises DA transmission. Western blot analysis showed reduced striatal DAT and vMAT2 levels, whereas TH, AADC and SV2C were not significantly altered (Fig. 1J, K). Levels of other clathrin-associated endocytic proteins and general synaptic markers were largely unchanged. HPLC analysis showed significantly reduced striatal DA total content at both 2–3 and 6–8 months, accompanied by increased DA metabolite HVA (Fig. 1L), consistent with impaired vesicular sequestration and enhanced degradation or oxidation of cytosolic DA. Finally, acute striatal slices from SJ1 cKO^DAT^ mice showed reduced DA release after both KCl-evoked depolarization and amphetamine-induced efflux (Fig. 1M–P). These findings establish SJ1 loss as a primary driver of dopaminergic terminal synaptopathy and DA deficiency, providing a model to examine how striatal circuits adapt to chronic presynaptic dopaminergic failure.

### SJ1 cKO^DAT^ mice show altered locomotion but preserved motor coordination

We next asked how widespread dopaminergic terminal pathology and striatal DA deficiency affect behavior. In the open field, SJ1 cKO^DAT^ mice showed reduced total distance traveled at both 2–4 and 6–8 months, indicating decreased basal locomotion or exploratory activity (Fig. 2A, B). Given the impaired amphetamine-induced DA release observed in acute slices, we then administered amphetamine and monitored locomotion for 60 min. Unexpectedly, SJ1 cKO^DAT^ mice displayed exaggerated amphetamine-induced hyperlocomotion at both ages (Fig. 2C–G), suggesting dysregulated downstream DA signaling despite impaired presynaptic DA efflux.

**Figure 2.**
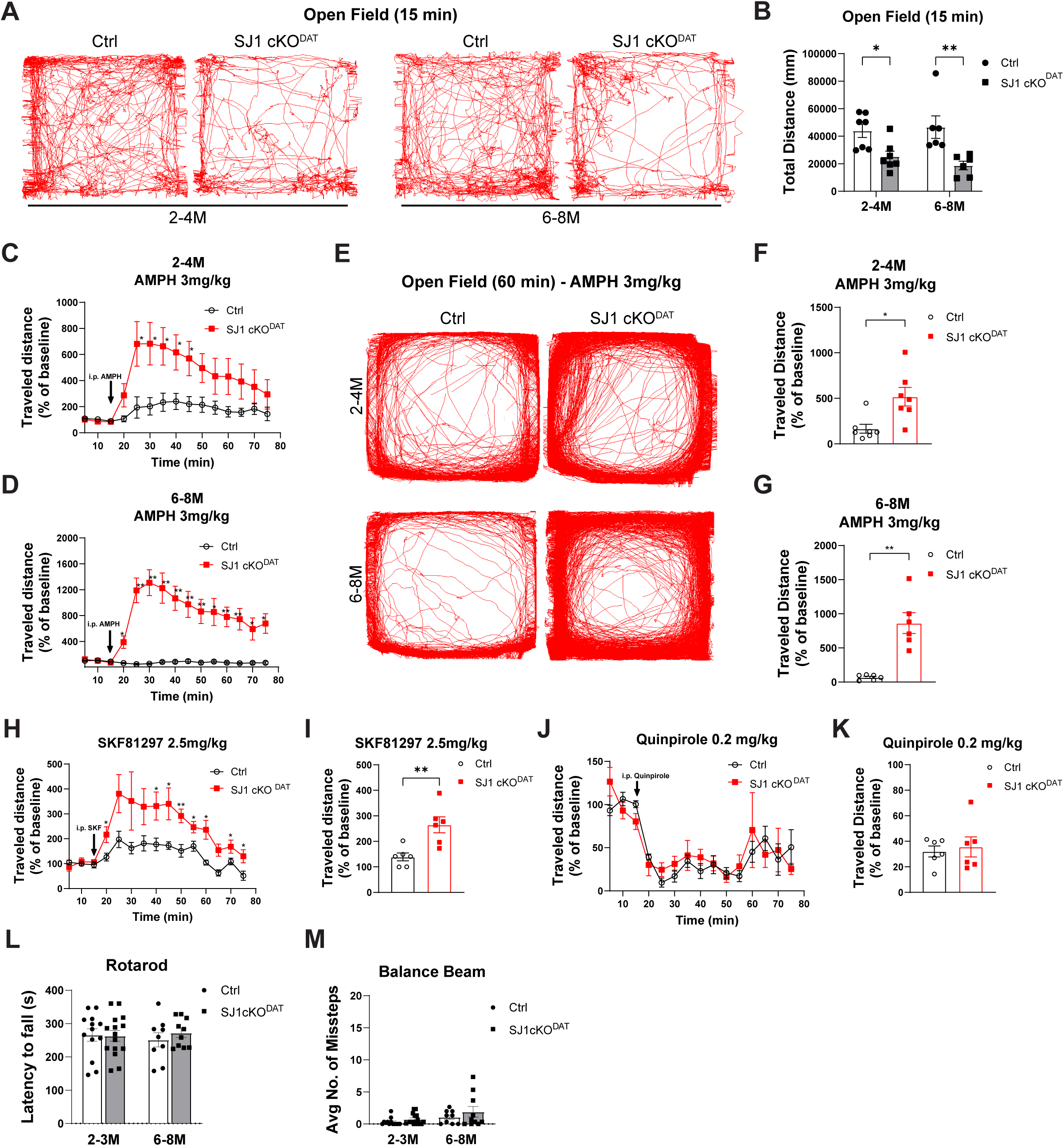
SJ1 cKO^DAT^ displayed altered locomotor activity but no coordination deficit (A) Representative open-field activity traces (15 min) from Ctrl and SJ1 cKO^DAT^ mice at both time points illustrate reduced locomotor activity observed in mutants. (B) Total distance traveled in open field test was recorded for 15 min. Significant p-value for total distance at both 2-4 and 6-8 months old SJ1 cKO^DAT^ mice indicates reduced spontaneous locomotion during baseline. n numbers used for each age group (2-4M: 7; 6-8M: 6). (C), (D) Quantification of average distance traveled, expressed as percent of baseline, over the time course. Locomotor activity (% of baseline) of SJ1 cKO^DAT^ mice increased over 60 min session treated with 3mg/kg amphetamine (AMPH) in 2-4 (C) and 6-8 (D) months old. n numbers used for each age group (2-4M: 7; 6-8M: 6). (E) Representative open-field activity traces (60 min) from Ctrl and SJ1 cKO^DAT^ mice at both time points illustrate AMPH-induced heightened hyperactivity observed in mutants. (F), (G) Average traveled distance in the open field. % of baseline across 60 min session in 2-4 (F) and 6-8 (G) months old treated with 3 mg/kg AMPH, showing significant increase in mutant’s locomotion. n numbers used for each age group (2-4M: 7; 6-8M: 6). (H), (I) Average total distance traveled over the time course (H) and average distance traveled in open field (I), both expressed as percent of baseline, showing significantly elevated locomotion in SJ1 cKO^DAT^ mice (age range 4-10M) in response to D1R agonist, SKF81297 at 2.5mg/kg. n=6 per genotype. (J), (K) Average total distance traveled over the time course (J) and average distance traveled in open field (K), both expressed as percent of baseline, showing comparable reduced locomotion in Ctrl and SJ1 cKO^DAT^ mice (age range 6-8M) in response to D2R agonist, quinpirole at 0.2mg/kg. n=6 per genotype. (L) Accelerated rotarod performance of two age groups. Latency to fall did not differ significantly between Ctrl and SJ1 cKO^DAT^ mice. n numbers used for Ctrl and SJ1 cKO^DAT^ mice (2–3M: 13 and 15; 6–8 M: 9 and 10). (M) Number of missteps during the balance beam test did not differ significantly between Ctrl and SJ1 cKO^DAT^ mice. n numbers used for Ctrl and SJ1 cKO^DAT^ mice (2–3M: 13 and 15; 6– 8M: 9 and 10). Data are represented as mean ± SEM. Statistical analysis: Welch’s unpaired t-test (F,G,I,K), ordinary two-way ANOVA with post-hoc Tukey’s test (B,L,M), repeated measures two-way ANOVA with post-hoc Tukey’s test (C,D,H,J); *p<0.05 and **p<0.01.

To test this possibility, we examined behavioral responses to dopamine receptor agonists. SJ1 cKO^DAT^ mice showed further increased hyperlocomotion after D1 receptor activation, whereas D2 receptor-induced hypolocomotion was comparable to controls (Fig. 2H–K), suggesting that D1R sensitivity is enhanced in the DA-deficient striatum. However, unlike SJ1^RQ^-KI mice, which show pronounced motor coordination deficits, SJ1 cKO^DAT^ mice performed normally on rotarod and balance beam tests (Fig. 2L, M). Thus, severe dopaminergic terminal dysfunction is accompanied by altered locomotor regulation but no detectable impairment on motor coordination, consistent with compensatory adaptations within the striatal circuit.

### Local adaptive induction of striatal dopamine-like neurons (DALNs) in SJ1 cKO^DAT^ mice

In addition to TH/DAT-positive dystrophic dopaminergic terminal clusters, we observed a striking emergence of TH-immunoreactive neurons within the striatum of SJ1 cKO^DAT^ mice (Fig. 3A). Striatal TH-positive interneurons (THINs) have been reported in rodents^24–26^, primates^27,28^ and humans^29–31^, but are typically undetectable by IHC in wild-type (WT) mice because of low TH protein expression^24,32,33^. In SJ1 cKO^DAT^ mice, however, these cells displayed robust TH protein accumulation; we therefore refer to them as striatal dopamine-like neurons (DALNs) to distinguish them from WT THINs.

**Figure 3.**
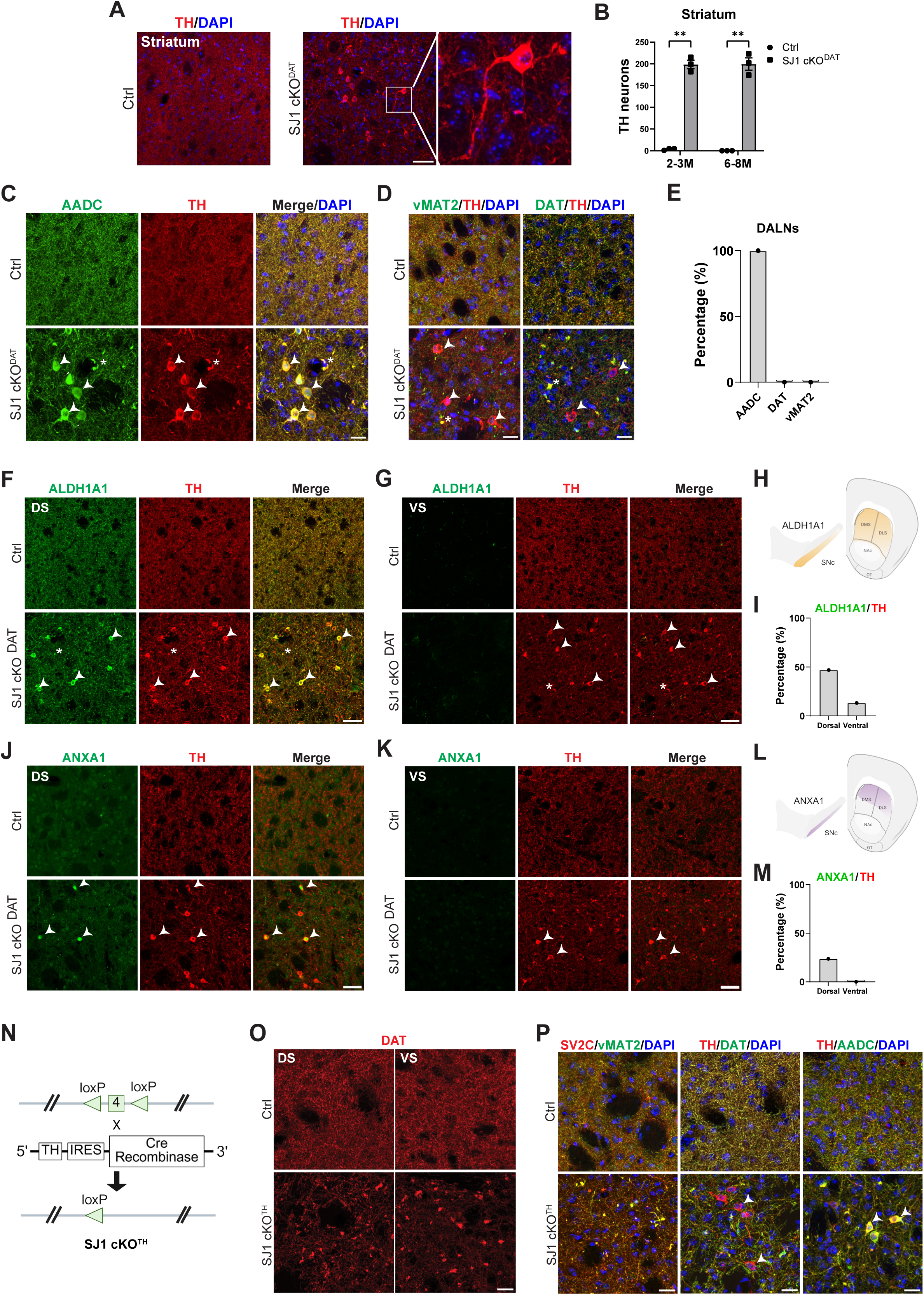
Robust appearance of DA-like neurons (DALNs) in the striatum of SJ1 cKO^DAT^ and SJ1 cKO^TH^ mouse models (A) TH immunostaining detected robust emergence of TH-positive neurons in the striatum of 2 months old SJ1 cKO^DAT^ mice but not in Ctrl. Scale bar: 50 µm. (B) TH-positive DALNs were selectively detected in 2-3 months and 6-8 months in SJ1 cKO^DAT^ striatum. Number of DALNs were quantified on single coronal striatum section of one hemisphere. n=3 mice per genotype. (C) Representative image of DALNs co-expressing TH and AADC in the striatum of 2 months old SJ1 cKO^DAT^ mice compared to Ctrl. White arrows: DALNs. Asterisk: clusters. Scale bar: 20 µm. (D) Double staining of vMAT2 and DAT with TH shows vMAT2 and DAT are clustered with TH at DA terminals but not expressed in DALNs. White arrows: DALNs. Asterisk: clusters. Scale bar: 20 µm. (E) Percentage analysis demonstrated that TH-positive DALNs were predominantly positive for AADC, whereas overlap with DAT and vMAT2 was negligible. (F), (G) Double staining of ALDH1A1 and TH shows ALDH1A1 is detected in both dystrophic DA terminals and DALNs in SJ1 cKO^DAT^ dorsal (F) but not ventral (G) striatum. White arrows: DALNs. Asterisk: clusters. Scale bar: 50 µm. (H) Simplified schematic illustration of ALDH1A1 labelling in SNc and striatum. (I) Percentage analysis revealed a higher proportion of DALNs co-expressing ALDH1A1 in the dorsal than in the ventral striatum. (J), (K) Double staining of ANXA1 and TH shows ANXA1 is detected in subpopulation of DALNs in SJ1 cKO^DAT^ dorsal (J) but not ventral (K) striatum. White arrows: DALNs. Scale bar: 50 µm. (L) Simplified schematic illustration of ANXA1 labelling in SNc and striatum. (M) Percentage analysis revealed a subset of DALNs co-expressing ANXA1 selectively in the dorsal but not ventral striatum. (N) Generation of SJ1 conditional KO in TH-IRES-Cre line by crossing SJ1 flx/flx mice with TH-IRES-Cre (TH-Cre;SJ1 flx/flx), hereafter referred to as SJ1 cKO^TH^. Appropriate littermate controls (SJ1 flx/flx or TH-Cre;SJ1 +/flx) were used in all experiments. (O) DAT staining of 2 months old Ctrl and SJ1 cKO^TH^ showed similar clustering phenotype in dorsal and ventral striatum. Scale bar: 20 µm. (P) Double staining of SV2C with vMAT2, TH with DAT and AADC showed the same dystrophic DA terminal pathology positive for all the markers in SJ1 cKO^TH^ striatum. Similarly, DALNs (white arrows) were robustly induced, co-expressing TH and AADC, but not DAT. Scale bar: 20 µm. Data are represented as mean ± SEM. Statistical analysis: multiple unpaired t-tests, with Holm-Šídák correction for multiple comparisons (B); **p<0.01.

DALNs appeared in both dorsal and ventral striatum (Fig. S5A), reaching approximately 200 cells per coronal section (Fig. 3B). Their emergence paralleled dopaminergic terminal dystrophy, becoming detectable at 3 weeks, peaking at 2–3 months, and persisting into adulthood and aging (Fig. S5B, C). Because SJ1 deletion is restricted to midbrain DANs, DALN induction represents a robust and sustained local striatal adaptation. To define DALN identity, we co-stained for major striatal neuronal populations, including MSNs and neuropeptide Y (NPY), parvalbumin (PV), calretinin (CR) and choline acetyltransferase (ChAT)-positive interneurons^34^. DALNs were negative for NPY, PV and ChAT, whereas a small subset partially co-expressed CR or Darpp32 (Fig. S5D–H), consistent with population heterogeneity.

We next examined whether DALNs acquire DA-like molecular features. All DALNs co-expressed TH and AADC, the two enzymes required to convert L-tyrosine to L-DOPA and then to DA (Fig. 3C). By contrast, the dopamine transporters vMAT2 and DAT, required for vesicular packaging and reuptake, colocalized with dystrophic terminals but were absent from DALN somata (Fig. 3D, E).

Given the heterogeneity of midbrain DANs^35,36^, we tested whether DALNs acquire subtype-specific features linked to distinct dopaminergic projections. Aldehyde dehydrogenase 1A1 (ALDH1A1), a marker of A9/SNc DANs that preferentially innervate dorsal striatum and are particularly vulnerable in PD^37,38^, was detected in SNc (Fig. S6A). Surprisingly, within the dorsal striatum, ALDH1A1 not only co-aggregated with DAT-positive dystrophic terminals but was also strongly expressed in dorsal DALNs, whereas it was largely absent in the ventral striatum (Fig. 3F–I; Fig. S6B). This spatially restricted pattern is consistent with A9 nigrostriatal innervation of the dorsal striatum and suggests circuit-dependent acquisition of A9-like features in dorsal DALNs.

We further examined annexin A1 (ANXA1), a marker of a highly vulnerable subtype within ALDH1A1-positive DANs^39,40^. ANXA1 labeled a subset of ALDH1A1/TH-positive DANs in the ventral SNc (Fig. S6C, D). Similarly, only a small subset of DALNs co-expressed ANXA1, selectively in the dorsolateral striatum, and these ANXA1-positive DALNs also expressed ALDH1A1 (Fig. 3J–M; Fig. S6E, F). In contrast, calbindin, a marker of A10/VTA DANs projecting mainly to ventral striatum, was detected in VTA TH-positive neurons but not in striatal DALNs (Fig. S6G–I).

Together, these findings indicate that DALNs exhibit circuit-dependent adaptation, with subpopulation of dorsal striatal DALNs acquiring A9/SNc DA-like molecular features.

### A second genetic SJ1 cKO model recapitulates dopaminergic terminal pathology and striatal DALN adaptation

To validate these findings, we generated an independent cell type-specific knockout model by crossing SJ1 flx/flx mice with the TH-IRES-Cre line, producing SJ1 cKO^TH^ mice (Fig. 3N). SJ1 was efficiently depleted in TH-positive DANs of the SNc and VTA (Fig. S7A). Similar to SJ1 cKO^DAT^ mice, SJ1 cKO^TH^ mice developed early-onset striatal dystrophic clusters positive for TH, AADC, DAT, vMAT2 and SV2C (Fig. 3O, P). These clusters were broadly distributed across the striatum, with ALDH1A1-positive clusters selectively enriched in the dorsal region (Fig. S7B), confirming involvement of both SNc and VTA projections.

Importantly, DALNs were also robustly induced in SJ1 cKO^TH^ striatum, paralleling terminal pathology. These DALNs co-expressed TH and AADC (Fig. 3P), and dorsal striatal subsets expressed ALDH1A1 (Fig. S7B, C), recapitulating the DA-like features. Thus, two independent DA-targeted SJ1 cKO models demonstrate that SJ1 loss in DANs induces cell-autonomous terminal pathology together with a secondary striatal DALN adaptation.

### Reporter labeling and electrophysiology distinguish DALNs from THINs in striatum

To genetically distinguish striatal DALNs from WT THINs, we crossed the Cre-dependent Ai9 tdTomato reporter line with both SJ1 cKO^TH^ and SJ1 cKO^DAT^ mice (Fig. 4A). In the midbrain, tdTomato expression was restricted to TH-positive DANs in both control and mutant mice (Fig. S7D, E).

**Figure 4.**
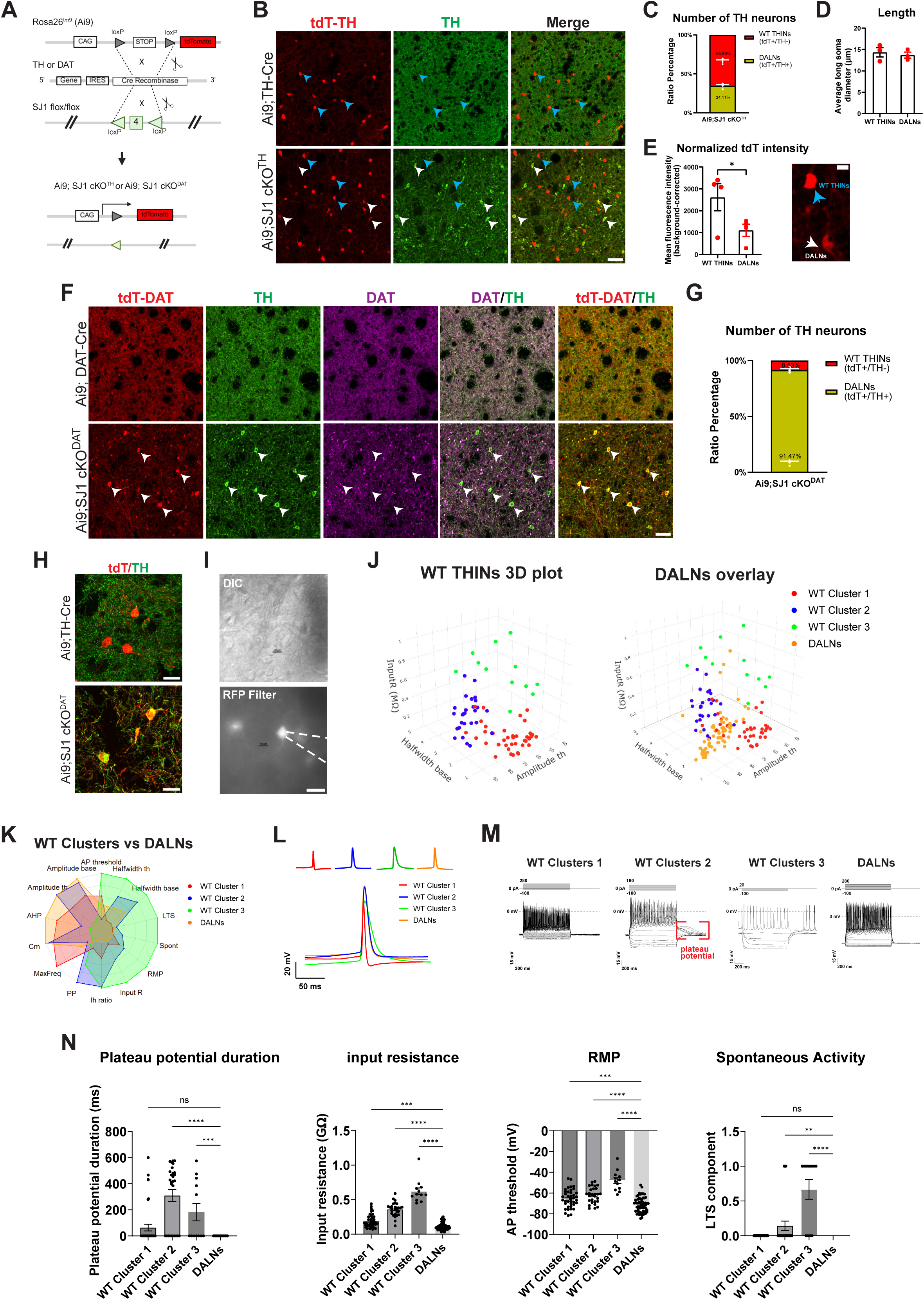
DALNs form a distinct electrophysiological subcluster with reduced intrinsic excitability compared to WT THINs (A) Simplified schematic showing the generation of Ai9-tdTomato labelled SJ1 cKO^TH^ and SJ1 cKO^DAT^ mice, referred to as Ai9;SJ1 cKO^TH^ and Ai9;SJ1 cKO^DAT^, respectively. Ai9-tdTomato reporter driven by CAG promoter expresses DA-specific tdTomato signal after Cre-mediated recombination by either TH-or DAT-IRES-Cre line. (B) Representative images of tdTomato expression and immunostaining of TH in the striatum of 2 months old Ai9;TH-Cre (Ctrl) and Ai9;SJ1 cKO^TH^. Blue arrows: tdTomato expressing WT THINs driven by TH-Cre recombination (negative for TH immunostaining). White arrows: DALNs expressing tdTomato signal (positive for TH immunostaining). Scale bar: 50 µm. (C) Percentage of TH-Cre driven tdTomato expressing neurons positive for TH immunostaining shown in (B). In average, 34.11% of total tdTomato expressing neurons were positive for TH immunostaining (DALNs). The number of neurons was quantified on a single stitched coronal striatum section for both tdTomato expressing neurons and tdTomato/TH-positive neurons. n=3 mice per genotype. (D) Quantification of the average long soma diameter of WT THINs and DALNs showed no significant differences in cell size. n=3 mice. (E) The average tdTomato fluorescence intensity of WT THINs (blue arrow) is significantly higher than DALNs (white arrow) when compared in the striatum of Ai9;SJ1 cKO^TH^. n=4 mice. Scale bar: 10 µm. (F) Representative images of tdTomato expression and immunostaining of TH and DAT in the striatum of Ai9;DAT-Cre (Ctrl) and Ai9;SJ1cKO^DAT^. tdTomato expressing striatal neurons were positive for TH but not DAT immunostaining (white arrows). Scale bar: 50 µm. (G) Percentage of tdTomato expressing neurons positive for TH immunostaining in DAT-Cre dependant background shown in (F). Majority of tdTomato expressing neurons were DALNs. The number of neurons was quantified on a single stitched coronal striatum section for both tdTomato expressing neurons and tdTomato/TH-positive neurons. n=3 mice per genotype. (H) Representative images of tdTomato expression and TH immunostaining in Ai9;TH-Cre and Ai9;SJ1cKO^DAT^ showing WT THINs and DALNs. Scale bar: 50 µm. (I) Representative images of a patch-clamped tdTomato-positive DALN in the dorsal striatum. Upper: differential interference contrast (DIC) image shows the patched neuron. Lower: epifluorescence image shows tdT (RFP filter) expression in the same cell. Scale bar: 20 µm. (J) A representative three-dimensional scatter plot showing clustering of WT THINs (left) into three electrophysiological subtypes. Overlay of DALNs (right) recorded from Ai9;SJ1 cKO^DAT^ mice onto the same three-dimensional feature space reveals that DALNs occupy a distinct region that differs from three other WT THINs clusters. (K) Radar plot summarizing normalized population mean values (0–1 scale) of 14 electrophysiological parameters for the three WT THINs clusters and DALNs, revealing distinct profiles across the four cell groups. (L) Representative traces of the first action potential elicited in WT THINs clusters and DALNs. DALNs showed action potential waveform properties that differ from WT cluster 1 and cluster 3, while closely resemble those of WT cluster 2. (M) Representative evoked firing traces of the three WT THIN clusters and DALNs in response to current injection. Notably, WT THIN cluster 2 exhibits a prolonged depolarizing plateau potential (red box), which is absent in DALNs. (N) Bar charts comparing four representative electrophysiological parameters among WT THINs clusters and DALNs, including plateau potential duration, input resistance, resting membrane potential (RMP), and spontaneous activity. Consistent with reduced intrinsic excitability inferred from passive membrane properties, DALNs exhibited altered membrane parameters compared to WT THINs. Data are presented as mean ± SEM. Statistical analysis: Kruskal–Wallis test followed by Dunn’s multiple comparison test (N); **p < 0.01, ***p < 0.001, ****p < 0.001.

We first examined TH-Cre-dependent labeling in the striatum. Consistent with previous reports, tdTomato-positive striatal neurons in Ai9;TH-Cre controls, representing genetically labeled WT THINs, were largely negative for TH immunostaining (Fig. 4B). In contrast, Ai9;SJ1 cKO^TH^ striatum contained two tdTomato-positive populations: approximately 34% were TH-immunoreactive DALNs, whereas the remaining 66% were TH-negative WT THINs (Fig. 4B, C). Although DALNs and neighboring WT THINs had comparable soma size (Fig. 4D), tdTomato fluorescence intensity was significantly lower in DALNs (Fig. 4E), suggesting distinct reporter or protein regulation in response to SJ1 loss.

We next examined DAT-Cre-dependent labeling. In Ai9;DAT-Cre controls, tdTomato labeled dopaminergic axons but not striatal neurons, consistent with the absence of DAT expression in WT THINs (Fig. 4F). Unexpectedly, Ai9;SJ1 cKO^DAT^ mice contained tdTomato-positive striatal neurons, nearly all of which were TH-immunoreactive DALNs (Fig. 4F, G). Although DAT protein was not detectable in DALN somata, this labeling pattern indicates that DALNs are molecularly distinct from WT THINs and are associated with DAN dysfunction.

To determine whether DALNs are also functionally distinct, we performed whole-cell patch-clamp recordings from tdTomato-positive DALNs in Ai9;SJ1 cKO^DAT^ striatal slices and WT THINs in Ai9;TH-Cre control slices (Fig. 4H, I). Consistent with previous studies^24,41^, WT THINs segregated into three subtypes by unsupervised clustering of 14 intrinsic membrane properties (Table S1; Fig. 4J; Fig. S8A, B). In contrast, DALNs were comparatively homogeneous and occupied a discrete region that did not overlap with WT THIN subtypes (Fig. 4J). Combined clustering also identified DALNs as a distinct population enriched in a single cluster, and radar plot analysis further confirmed broad differences in intrinsic membrane properties (Fig. 4K; Fig. S8A).

Analysis of the first evoked action potential showed that DALNs differed from WT cluster 1 and cluster 3 neurons but partially resembled cluster 2 neurons (Fig. 4L). However, step-current recordings revealed markedly reduced DALN excitability compared with WT THIN subtypes (Fig. 4M). Notably, whereas WT cluster 2 neurons displayed prolonged depolarizing plateau potentials, a feature known to be modulated by DA, none of the recorded DALNs exhibited detectable plateau potentials (0/63; Fig. 4M, N). DALNs also showed reduced input resistance, a more hyperpolarized resting membrane potential and absence of spontaneous firing (Fig. 4N). This coordinated reduction in excitability may decrease GABAergic inhibitory output and thereby contribute to striatal circuit compensation under DA deficient conditions.

Finally, recordings from labeled SNc and VTA DANs in Ai9;DAT-Cre control and SJ1 cKO^DAT^ mice revealed only mild changes in several action-potential-related parameters in SNc neurons, without broad alterations in intrinsic membrane properties (Fig. S8C–N). Thus, SJ1 loss primarily disrupts DA transmission at axon terminals, whereas DALNs emerge as a distinct striatal adaptive state with reduced intrinsic excitability.

### DALNs acquire dopaminergic marker proteins from midbrain DANs rather than through intrinsic transcription

A key mechanistic question was how striatal neurons acquire DA-like features. We first combined RNAscope with immunostaining to examine TH mRNA and protein expression. As positive controls, both TH protein and Th mRNA were detected in midbrain DANs, locus coeruleus noradrenergic neurons, and TH-positive neurons in the olfactory bulb, cortex and cerebellum (Fig. 5A; Fig. S9A). Surprisingly, although DALNs in both SJ1 cKO^DAT^ and SJ1 cKO^TH^ striatum accumulated high levels of TH protein, they lacked the corresponding *Th* transcripts (Fig. 5A, D, E; Fig. S9B). By contrast, a distinct striatal population was *Th* mRNA-positive but TH protein-negative, consistent with WT THINs and also observed in control DAT-Cre, TH-Cre and B6 striatum (Fig. 5A; Fig. S9B–E).

**Figure 5.**
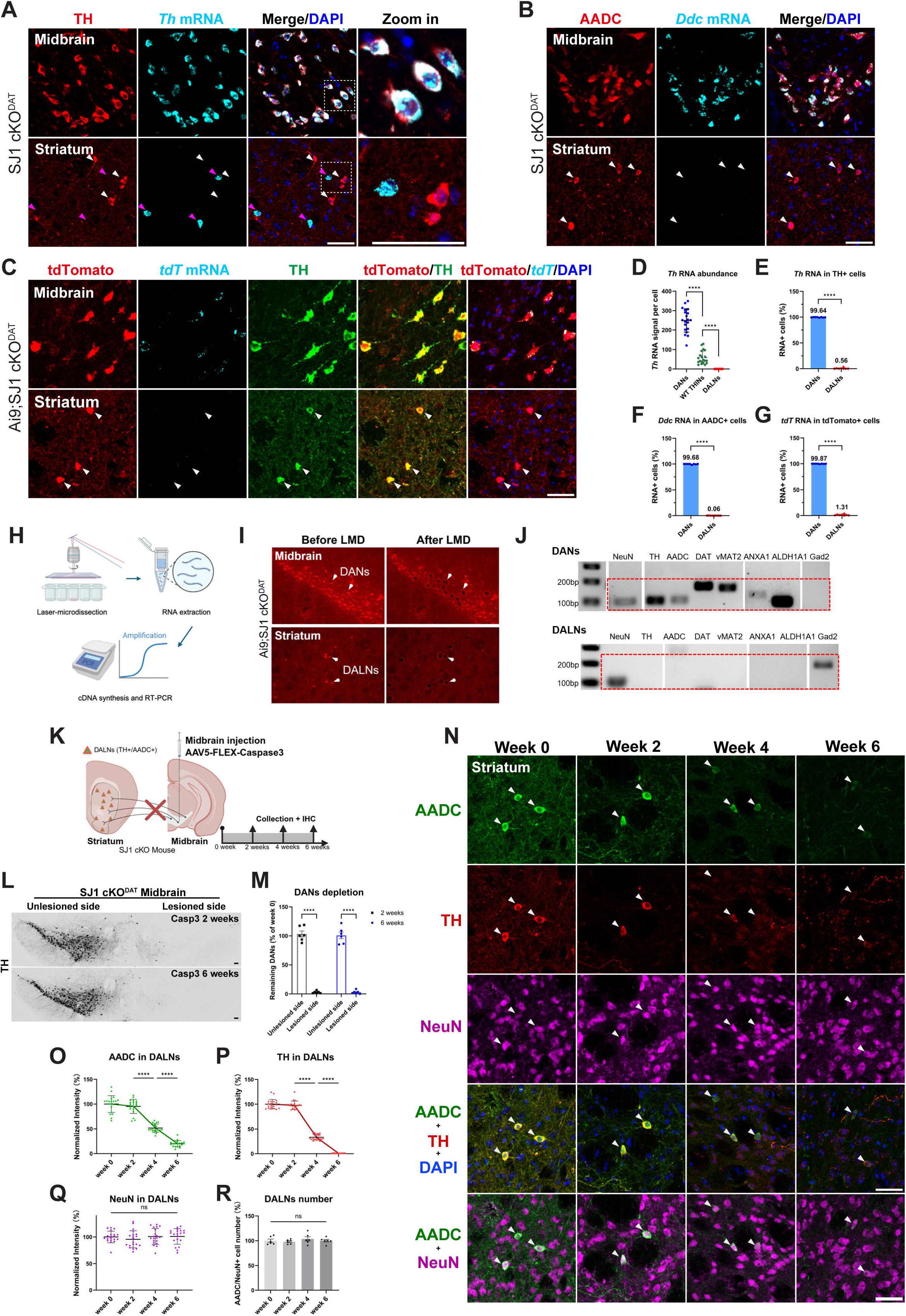
Dopaminergic marker proteins in DALNs are not intrinsically synthesized but derived from DANs (A) Representative RNAscope-IF images of TH protein and *Th* mRNA in the midbrain and striatum of SJ1 cKO^DAT^ mice. Midbrain DANs were positive for both TH protein and *Th* mRNA. In the striatum, DALNs were positive for TH protein but negative for *Th* mRNA (white arrows), WT THINs were negative for TH protein but positive for *Th* mRNA (pink arrows). Scale bar: 20 µm. (B) Representative RNAscope-IF images of AADC protein and *Ddc* mRNA in SJ1 cKO^DAT^ mice. Midbrain DANs were positive for both AADC protein and *Ddc* mRNA, whereas striatal AADC-positive DALNs lacked detectable *Ddc* mRNA (white arrows). Scale bar: 20 µm. (C) Representative RNAscope-IF images of TH, tdTomato protein and *tdTomato* mRNA (*tdT*) in Ai9;SJ1 cKO^DAT^ mice. Midbrain TH/tdTomato-positive DANs expressed the *tdTomato* mRNA, whereas striatal TH/tdTomato-positive DALNs lacked detectable *tdTomato* mRNA (white arrows). Scale bar: 20 µm. (D) Quantification of *Th* mRNA abundance in DANs, WT THINs and DALNs. *Th* mRNA abundance was quantified as the number of RNAscope puncta per cell [or normalized puncta intensity per cell]. The *Th* mRNA abundance was highest in DANs, lower in WT THINs and undetectable in DALNs. Each point represents one cell. Twenty cells from four mice were analysed, and cells from the same mouse were averaged for statistical analysis (n=4 mice). (E) –(G) Quantification of the percentage of protein-positive cells that were also positive for the corresponding mRNA in the midbrain and striatum. Midbrain DANs showed robust expression of the corresponding mRNAs, whereas striatal DALNs showed no detectable expression of *Th* (E), *Ddc* (F) or *tdTomato* (G) mRNA. Each point represents one tissue section. Eight sections from four mice were analysed per experiment, and sections from the same mouse were averaged for statistical analysis (n=4 mice). (H) Schematic illustration of laser capture microdissection followed by reverse transcription PCR (LCM-RT-PCR) analysis of midbrain DANs and striatal DALNs. (I) Representative red-channel fluorescence images of Ai9;SJ1 cKO^DAT^ midbrain and striatum before and after LCM, showing selective isolation of tdTomato-positive DANs and DALNs. (J) Representative agarose gel images of PCR products from laser-captured DANs and DALNs. DANs expressed *Rbfox3* (NeuN) and the dopaminergic markers *Th* (TH), *Ddc* (AADC), *Slc6a3* (DAT), *Slc18a2* (vMAT2), *Anxa1* (ANXA1) and *Aldh1a1* (ALDH1A1), but not the GABAergic neuronal marker *Gad2* (GAD2). In contrast, DALNs expressed NeuN and Gad2, but none of the examined dopaminergic markers, including *Th*, *Ddc*, *Slc6a3*, *Slc18a2*, *Anxa1* and *Aldh1a1*. (K) Schematic illustration of the experimental design. High-dose AAV5-FLEX-Caspase3 injection into SJ1 cKO^DAT^ mice. Brains were collected 2, 4 and 6 weeks after injection for immunohistochemical analysis. (L) Representative TH immunostaining of lesioned and unlesioned sides at 2 and 6 weeks after injection, showing complete loss of TH-positive DANs at the lesioned side by 2 weeks, with no further change observed at 6 weeks. (M) Quantification of remaining TH-positive DANs at lesioned and unlesioned sides at 2 and 6 weeks. Each point represents one tissue section. Eight sections from four mice were analysed per experiment, and sections from the same mouse were averaged for statistical analysis (n=4 mice). (N) Representative images of AADC, TH and NeuN immunostaining in the striatum at 0, 2, 4 and 6 weeks after midbrain DAN ablation. White arrows indicate DALNs. (O) –(Q) Quantification of AADC (O), TH (P) and NeuN (Q) fluorescence levels in DALNs. AADC and TH levels progressively decreased after DAN ablation, whereas NeuN levels remained unchanged. Each point represents one cell. Twenty cells from four mice were analysed per time point, and cells from the same mouse were averaged for statistical analysis (n = 4 mice). (R) Quantification of striatal DALNs, identified as TH/AADC/NeuN triple-positive cells, at 0, 2, 4 and 6 weeks after DAN ablation. DALN numbers remained unchanged. Each point represents one tissue section. Six sections from four mice were analysed per time point, and sections from the same mouse were averaged for statistical analysis (n=4 mice). Data in dot plots are presented as median with interquartile range. Bar charts are presented as mean ± SEM. Statistical analysis: repeated-measures one-way ANOVA with post hoc Tukey’s test (D), paired t-test (E-G), two-tailed paired t-tests with Holm–Šídák correction (M), ordinary one-way ANOVA with post hoc Tukey’s test (O-R); *p < 0.05, **p < 0.01, ***p < 0.001 and ****p < 0.0001.

RNAscope further confirmed the absence of *Ddc*, *Aldh1a1* and *Anxa1* mRNAs in DALNs, despite expression of the corresponding proteins in subsets of cells (Fig. 5B, F; Fig. S10A, B). DALNs also lacked DAT protein and the corresponding *Slc6a3* mRNA (Fig. S10C). In contrast, housekeeping transcripts such as *Polr2a* and *Ppib* were readily detected, whereas the bacterial negative control probe *DapB-C1* was not (Fig. S10D–F), supporting RNA integrity and the specificity of the negative dopaminergic transcript signals. Thus, DALNs contain dopaminergic proteins without undergoing intrinsic dopaminergic transcriptional reprogramming.

Reporter analyses supported this conclusion. In Ai9;SJ1 cKO^DAT^ mice, DALNs were tdTomato protein-positive but tdTomato mRNA-negative, indicating that tdTomato protein was not generated by DAT-Cre-mediated transcription within these cells (Fig. 5C, G). In Ai9;SJ1 cKO^TH^ mice, WT THINs and DALNs showed distinct reporter/transcript profiles: WT THINs were *Th* mRNA-positive and tdTomato protein/mRNA-positive, whereas DALNs were TH and tdTomato protein-positive but mRNA-negative (Fig. S11A, B). Ai9;TH-Cre controls confirmed that WT THINs normally expressed *Th* and tdTomato mRNAs and tdTomato protein but lacked detectable TH protein (Fig. S11C, D).

To independently validate the transcriptional profile of DALNs, we isolated tdTomato-positive midbrain DANs and striatal DALNs from Ai9;SJ1 cKO^DAT^ mice by laser-capture microdissection (Fig. 5H, I). RT-PCR confirmed the absence of *Th* and other dopaminergic transcripts in DALNs compared with DANs, while detecting the GABAergic transcript Gad2 (Fig. 5J), supporting their striatal interneuron origin.

These findings suggested that DALNs acquire dopaminergic marker proteins from an external source. To test whether midbrain DANs provide this source, we selectively ablated DANs in SJ1 cKO^DAT^ mice using AAV-mediated Cre-dependent caspase-3 expression (Fig. 5K). DANs were largely eliminated within 2 weeks, as confirmed by TH staining (Fig. 5L, M). Loss of dopaminergic innervation caused a progressive decline in TH and AADC protein levels in striatal DALNs from 2 to 6 weeks after ablation (Fig. 5N–P). Importantly, the total number of DALNs remained unchanged, as quantified by NeuN staining (Fig. 5Q, R), indicating that DALNs were retained but lost their DA-like protein state. Thus, DALNs maintain dopaminergic features through long-range crosstalk with midbrain DANs.

### Selective DAN–DALN connectivity mediates long-range protein transfer

The dependence of DALN dopaminergic features on intact DAN innervation led us to test whether midbrain DANs engage DALNs through a long-range transfer pathway. We first labeled DANs using Cre-dependent viral tracing by unilaterally injecting AAV5-FLEX-tdTomato into the ventral midbrain of SJ1 cKO^TH^, SJ1 cKO^DAT^ and the corresponding TH-Cre or DAT-Cre controls (Fig. 6A). Two weeks later, TH immunostaining confirmed robust tdTomato expression in midbrain DANs and their striatal projections (Fig. 6B, C). Strikingly, tdTomato-positive DALNs were detected in the striatum of both SJ1 cKO^TH^ and SJ1 cKO^DAT^ mice (Fig. 6C–E). Because the virus was restricted to the midbrain, this labeling suggests long-range transfer of DAN-derived material, most likely protein, to striatal DALNs.

**Figure 6.**
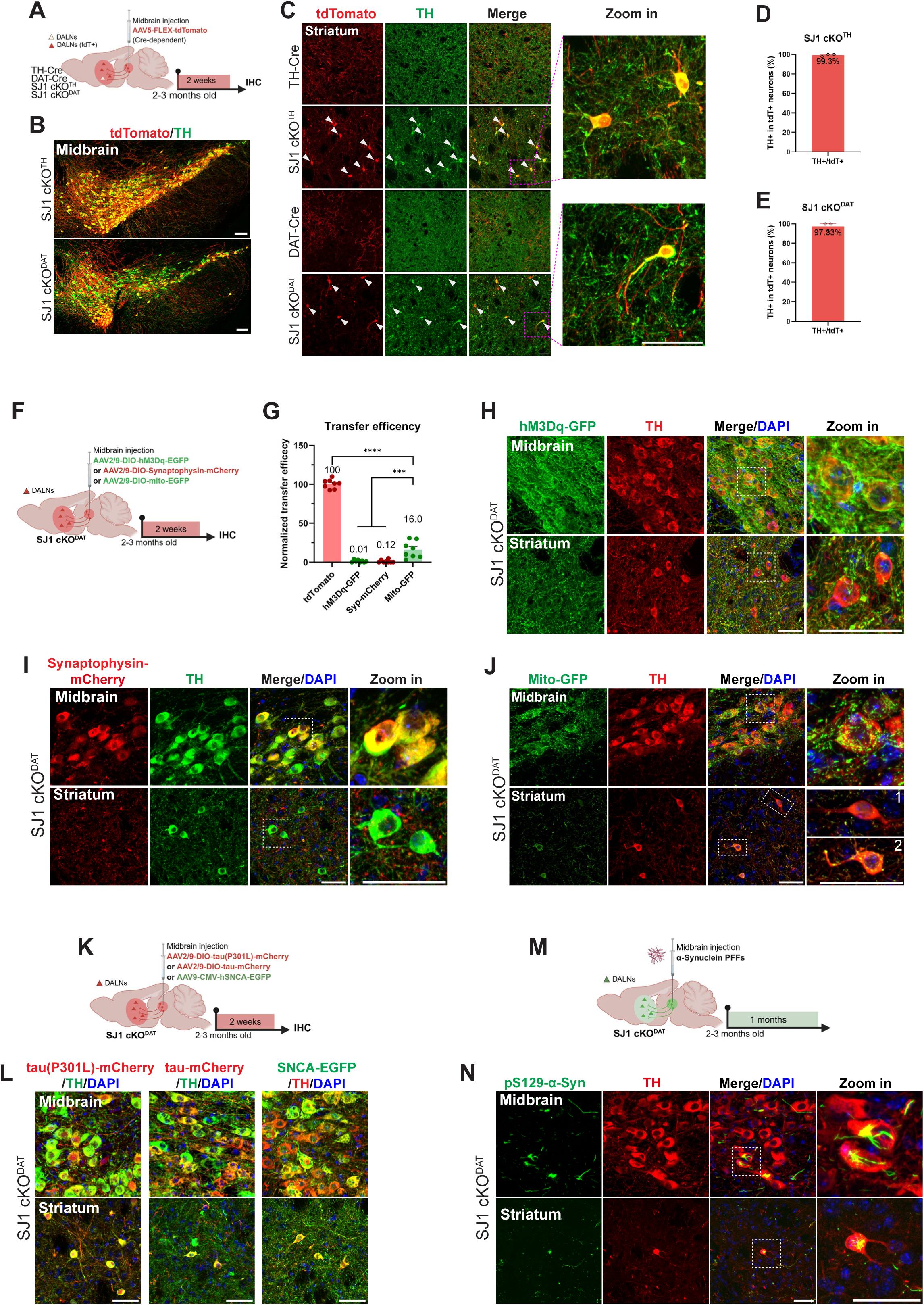
DANs form a selective, long-range cargo-transfer pathway to DALNs for physiological and pathological proteins (A) Schematic showing experiment procedure and timeline. AAV5-FLEX-tdTomato (Cre-dependent) was injected into the ventral midbrain of SJ1 cKO^TH^, SJ1 cKO^DAT^ mice and Cre mouse controls at 2-3 months old. The animal brains were collected 2 weeks post injection for IHC. (B) Representative images of TH immunostaining and tdTomato expression in the injected midbrain DANs of both SJ1 cKO^TH^ and SJ1 cKO^DAT^ mice. Scale bar: 100 µm. (C) Representative images of TH immunostaining and tdTomato expression in the striatum. While tdTomato in TH-Cre and DAT-Cre mice only labeled some DA fibers, it labeled both DA fibers and DALNs in SJ1 cKO^TH^ and SJ1 cKO^DAT^. White arrows: DALNs double positive for TH and tdTomato signals. Scale bar: 20 µm. (D), (E) Percentages of tdTomato-positive striatal neurons co-expressing TH in SJ1 cKO^TH^ (D) and SJ1 cKO^DAT^ (E) mice. Cells were quantified from one stitched coronal striatal section per mouse. n=3 mice per genotype. (F) Experimental design for assessing cargo transfer. AAV2/9-DIO-hM3Dq-EGFP, AAV2/9-DIO-synaptophysin-mCherry or AAV2/9-DIO-mito-GFP was injected into the ventral midbrain of SJ1 cKO^DAT^ mice, and brains were collected 2 weeks later for IHC. (G) Quantification of transfer efficiency into DALNs, normalized to freely diffusible tdTomato signal set to 100%. hM3D-GFP and synaptophysin-mCherry showed minimal transfer, whereas mito-GFP reached approximately 16% of the reference level. Eight sections from four mice were analysed per construct, with sections from the same mouse averaged for statistical analysis (n=4 mice). (H) –(J) Representative midbrain and striatal images following expression of hM3D-GFP (H), synaptophysin-mCherry (I) and mito-GFP (J). hM3D-GFP and synaptophysin-mCherry showed little or no signal in DALNs, whereas punctate mito-GFP signals were detected in a subset of DALNs. Enlarged views in (J) show DALNs with low (cell 1) or abundant (cell 2) mito-GFP signals. Scale bars: 20 µm. (K) Experimental design for assessing transfer of disease-associated proteins. AAV2/9-DIO-tau(P301L)-mCherry, AAV2/9-DIO-tau-mCherry or AAV9-CMV-hSNCA-EGFP was injected into the ventral midbrain of SJ1 cKO^DAT^ mice, and brains were collected 2 weeks later for IHC. (L) Representative merged TH, reporter and DAPI images showing tau(P301L)-mCherry, tau-mCherry and α-synuclein-EGFP in midbrain DANs and striatal DALNs. Scale bar: 20 µm. (M) Experimental design for α-synuclein PFF injection into the ventral midbrain of SJ1 cKO^DAT^ mice. Brains were collected 1 month later for IHC. (N) Representative TH and pS129-α-synuclein images showing phosphorylated α-synuclein aggregates in midbrain DANs and striatal DALNs. Scale bar: 20 µm. Data are presented as mean ± SEM. *p < 0.05, **p < 0.01, ***p < 0.001 and ****p < 0.0001. Statistical analysis: ordinary one-way ANOVA with post hoc Tukey’s test (G); ***p < 0.001 and ****p < 0.0001.

We next used a Cre-independent strategy by injecting AAV2-mTH-EGFP into the midbrain of B6 control and SJ1 cKO mice (Fig. S12A). EGFP labeled midbrain DANs and was detected selectively in TH-positive DALNs in the SJ1 cKO striatum (Fig. S12B–E). This result was further confirmed in Ai9;SJ1 cKO^TH^ mice, in which EGFP was present exclusively in tdTomato/TH double-positive DALNs and absent from tdTomato-positive/TH-negative WT THINs (Fig. S12K, L).

Finally, we injected pan-neuronal AAV-hSyn-mCherry into the midbrain of SJ1 cKO mice (Fig. S12F). mCherry was expressed in multiple neuronal populations near the injection site, including DANs (Fig. S12G). In the striatum, however, mCherry-positive neurons were found exclusively among TH-positive DALNs (Fig. S12H–J). By contrast, direct striatal injection of AAV-hSyn-mCherry labeled diverse local neurons, including DARPP32-positive MSNs (Fig. S12M, N), arguing against nonspecific viral leakage from dystrophic dopaminergic terminals.

These three independent labeling strategies reveal a selective and previously unrecognized long-range DAN–DALN connection that mediates inter-neuronal protein transfer following SJ1 loss in the dopaminergic system.

### DAN–DALN transfer is cargo-selective and conveys physiological and pathological proteins

The viral-tracing experiments above showed that soluble fluorescent proteins, including tdTomato, EGFP and mCherry, can be transferred from midbrain DANs to DALNs, consistent with the endogenous transfer of soluble DA enzymes such as TH and AADC. We therefore tested whether this pathway accommodates proteins with different subcellular localizations.

Transmembrane proteins, including Channelrhodopsin-2 (ChR2)-GFP and DREADD hM3Dq-GFP, were largely retained in dopaminergic axons and dystrophic terminals, with little detectable transfer to DALNs (Fig. 6G, H; Fig. S12O, P), similar to the restricted localization of endogenous DAT. Nuclear cargo showed even stronger restriction: histone H2B-GFP robustly labeled DAN nuclei in the midbrain but was not detected in DALNs (Fig. S12O, Q).

Organelle-associated cargoes showed distinct transfer patterns. Synaptophysin-mCherry, a synaptic vesicle-associated protein, was largely confined to dopaminergic terminals and showed little or no detectable transfer to DALNs (Fig. 6G, I), consistent with the absence of endogenous vMAT2 from DALNs. By contrast, mito-EGFP showed heterogeneous transfer: some DALNs contained little or no signal, whereas others displayed punctate labeling within somata or neurites, indicating that mitochondrial cargo can enter a subset of DALNs, although less efficiently than soluble proteins (Fig. 6G, J).

We next asked whether disease-associated proteins could access this pathway. α-synuclein-GFP expressed in midbrain DANs was specifically detected in DALNs (Fig. 6K, L). Likewise, both WT tau and pathogenic P301L tau, associated with frontotemporal dementia and parkinsonism, were transferred from midbrain DANs to striatal DALNs (Fig. 6K, L). In addition, injection of α-synuclein preformed fibrils into the midbrain induced pS129 α-synuclein in DANs and striatal fibers, with pS129-positive pathology also detected in a subset of DALNs (Fig. 6M, N), suggesting that fibrillar α-synuclein seeds can traverse this route and amplify aggregation in recipient cells.

Together, these findings define DAN–DALN transfer as a cargo-selective pathway that preferentially conveys soluble proteins while restricting membrane, nuclear and organelle cargoes. This pathway is not limited to physiological protein sharing but can also mediate the spread of disease-related proteins, linking early buffering to potential pathological propagation.

### DAN–DALN protein transfer supports local DA compensation and circuit-level protection

Because DALNs acquire TH and AADC from DANs, we asked whether they are capable of local DA synthesis. Immunostaining with an anti-dopamine antibody revealed that approximately 75% of DALNs were DA-positive, consistent with endogenous DA production (Fig. 7A–D). TH in both DALNs and bona fide midbrain DANs was detected in its phosphorylated Ser40 form, a modification associated with TH activation, supporting that transferred TH remains catalytically competent (Fig. 7E–G). L-DOPA treatment further increased intracellular DA levels in DALNs, similarly to DANs (Fig. 7H–J), indicating that transferred AADC is functional and can convert L-DOPA to DA.

**Figure 7.**
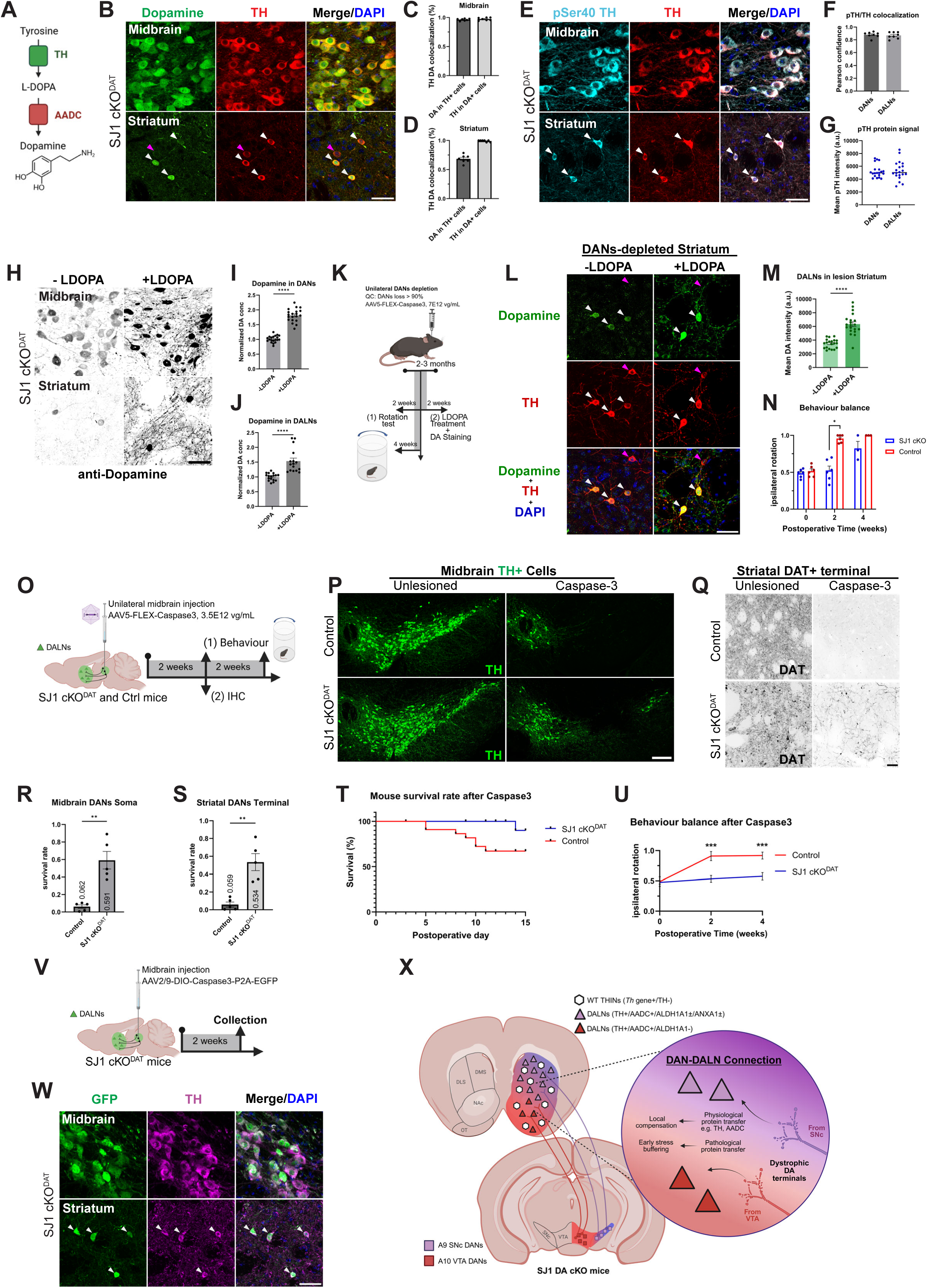
DAN-to-DALN protein transfer enables local DA synthesis and protects the dopaminergic system from acute lesions (A) Schematic illustration of dopamine synthesis mediated by TH and AADC. (B) Representative TH and dopamine staining in the midbrain and striatum of SJ1 cKO^DAT^ mice. Midbrain DANs were positive for both TH and dopamine. In the striatum, white arrows indicate dopamine-positive DALNs, whereas pink arrows indicate dopamine-negative DALNs. (C, D) Quantification of TH and dopamine co-expression in the midbrain (C) and striatum (D). TH and dopamine signals showed extensive overlap in DANs. In the striatum, nearly all dopamine-positive cells were TH-positive, whereas approximately 75% of TH-positive DALNs were dopamine-positive. Each point represents one tissue section. Eight sections from four mice were analysed per group, and sections from the same mouse were averaged for statistical analysis (n=4 mice). (E) Representative TH and phospho-Ser40 TH immunostaining in the midbrain and striatum. White arrows indicate DALNs positive for both TH and phospho-Ser40 TH. (F) Colocalization analysis of TH and phospho-Ser40 TH signals in the midbrain and striatum using Pearson’s correlation coefficient in striatum region. Each point represents one tissue section. Eight sections from four mice were analysed, and sections from the same mouse were averaged for statistical analysis (n=4 mice). (G) Quantification of phospho-Ser40 TH fluorescence intensity in DANs and DALNs. Each point represents one cell. No significant difference was detected between the two cell populations. Twenty cells from four mice were analysed per group, and cells from the same mouse were averaged for statistical analysis (n=4 mice). (H) Representative dopamine staining in the midbrain and striatum of SJ1 cKO^DAT^ mice before and after L-DOPA administration. (I), (J) Quantification of dopamine levels in midbrain DANs (I) and striatal DALNs (J) before and after L-DOPA administration. Dopamine levels increased in both cell populations after treatment. Each point represents one cell. Twenty cells from four mice were analysed per group, and cells from the same mouse were averaged for statistical analysis (n=4 mice). (K) Experimental design for high-dose dopaminergic lesion. AAV5-FLEX-Caspase3 (7 × 10^12^ vg/ml) was unilaterally injected into the ventral midbrain of SJ1 cKO^DAT^ mice. Rotational behaviour was assessed at 0, 2 and 4 weeks, and a separate cohort received L-DOPA before dopamine staining at 2 weeks. (L) Representative striatal TH and dopamine staining 2 weeks after high-dose AAV5-FLEX-Caspase3 injection, with or without L-DOPA treatment. (M) Quantification of dopamine levels in striatal DALNs showing increased dopamine signal after L-DOPA treatment. Each point represents one cell. Twenty cells from four mice were analysed per group, and cells from the same mouse were averaged for statistical analysis (n=4 mice). (N) Ipsilateral rotation after high-dose dopaminergic lesion. Control and SJ1 cKO^DAT^ mice showed balanced behaviour at baseline. Control mice developed marked ipsilateral rotation by 2 weeks, whereas SJ1 cKO^DAT^ mice maintained behavioural balance until 4 weeks. Each point represents one mouse. Six mice per genotype were analysed at baseline and 2 weeks, and three mice per genotype were analysed at 4 weeks. (O) Experimental design for moderate-dose dopaminergic lesion. AAV5-FLEX-Caspase3 (3.5 × 10^12^ vg/ml) was unilaterally injected into the ventral midbrain of control and SJ1 cKO^DAT^ mice. Rotational behaviour was assessed at 0, 2 and 4 weeks, and brains were collected at 2 weeks for immunohistochemical analysis. (P), (Q) Representative TH immunostaining of midbrain DAN soma (P) and DAT immunostaining of striatal DA terminals (Q) 2 weeks after lesion. Scale bars: 200 µm (P) and 20 µm (Q). (R), (S) Quantification of surviving midbrain TH-positive DAN soma (R) and striatal DAT-positive terminals (S). SJ1 cKO^DAT^ mice showed significantly greater survival than controls. Each point represents one mouse. Each point represents one mouse. (T) Postoperative survival curves showing higher survival in SJ1 cKO^DAT^ mice than in controls after dopaminergic lesion. Postoperative survival was analysed in 50 mice, including 28 SJ1 cKO^DAT^ mice and 22 controls. Survival curves were compared using the log-rank (Mantel–Cox) test. (U) Ipsilateral rotation after moderate-dose lesion. Both genotypes showed balanced behaviour at baseline. Control mice developed sustained ipsilateral rotation at 2 and 4 weeks, whereas SJ1 cKO^DAT^ mice maintained behavioural balance. Significant differences were detected between genotypes at both time points. Each point represents one mouse. Four mice per genotype were analysed. (V) Experimental design for tracing lesion-associated protein transfer. AAV2/9-DIO-Caspase3-P2A-EGFP was injected into the ventral midbrain of SJ1 cKO^DAT^ mice, and brains were collected 2 weeks later for immunohistochemical analysis. (W) Representative GFP immunostaining in the midbrain and striatum following AAV2/9-DIO-Caspase3-P2A-EGFP injection. GFP signal was detected in both midbrain DANs and striatal DALNs, indicating transfer of lesion-associated protein cargo. (X) Working model. In the SJ1 DA cKO striatum, dystrophic DA terminals establish a circuit-dependent, long-range protein transfer pathway with local DALNs. Transfer of physiological dopaminergic proteins, including TH and AADC, supports local dopamine compensation, whereas redistribution of pathological proteins may buffer cellular stress in DANs and reduce neuronal vulnerability. Data in dot plots are presented as median with interquartile range. Bar charts are presented as mean ± SEM. Statistical analysis: paired t-test (G), Welch’s unpaired t-test (I, J, M, R, S), mixed-effects model with genotype and time as factors, followed by Šídák’s multiple-comparisons test (N), repeated-measures two-way ANOVA with genotype and time as factors, followed by Šídák’s multiple-comparisons test (U); *p < 0.05, **p < 0.01, ***p < 0.001 and ****p < 0.0001.

To exclude the possibility that DA detected in DALNs simply reflects transport from midbrain DANs, we performed unilateral DAN ablation using high-dose AAV5-FLEX-caspase-3 in SJ1 cKO^DAT^ mice (Fig. 7K). Under this condition, in which the dense dopaminergic fiber background was largely removed, DALN DA signals were more readily detected and remained responsive to L-DOPA (Fig. 7L, M), further supporting local DA handling and synthesis. Behaviorally, DAT-Cre control mice with unilateral DAN lesion displayed robust unidirectional rotation, whereas SJ1 cKO^DAT^ mice maintained motor balance at 2 weeks after high-dose lesion. This compensation was not sustained at 4 weeks, consistent with gradual loss of DA-like features in DALNs after dopaminergic denervation (Fig. 7N).

We next tested whether this circuit adaptation influences neuronal vulnerability. Using low-dose AAV-FLEX-caspase-3 to induce milder ablation, we found that SJ1 cKO^DAT^ mice retained more TH-positive DANs in the midbrain and more DAT-positive dopaminergic fibers in the striatum on the lesioned side than controls (Fig. 7O–S), suggesting increased resistance to apoptotic stress. At the animal level, SJ1 cKO^DAT^ mice showed higher post-lesion survival and a more balanced rotational phenotype at 2 and 4 weeks after ablation (Fig. 7T, U), consistent with partial preservation of dopaminergic circuit function. A similar protective effect was observed after 6-OHDA lesion, indicating that resistance was not specific to one injury paradigm (Fig. S13A–G). Notably, fluorescently labeled caspase-3 was also detected in DALNs when expressed in midbrain DANs of SJ1 cKO^DAT^ mice (Fig. 7V, W), suggesting that injury-related cargo can also enter DALNs through this pathway.

These findings indicate that DAN–DALN protein transfer supports local DA compensation and may also redistribute and buffer pathological burden at the circuit level, thereby reducing vulnerability to dopaminergic and apoptotic insults.

### DALN adaptation is shared across PD-linked endocytic mutants but constrained by developmental timing and disease context

To determine whether DALN emergence represents a broader response in PD models, we examined additional mouse models carrying loss-of-function mutations in PD-linked clathrin uncoating genes (Fig. 8A). In previous studies, we reported milder dopaminergic terminal dystrophy and striatal DALN-like adaptation in SJ1^RQ^-KI and Aux-KO/SJ1^RQ^-KI double mutant mice^21,22^ (Fig. 8B). Re-analysis of these models showed that, beyond TH and AADC, a subset of dorsal striatal DALNs also expressed ALDH1A1 and ANXA1 (Fig. 8C, D). In SJ1^RQ^-KI and Ai9;SJ1^RQ^-KI mice, we further confirmed the absence of *Th, Ddc* and *tdT* mRNA in DALNs and observed selective transfer of AAV-expressed mCherry protein from DANs to DALNs (Fig. 8E-H. S14A), supporting the same non-transcriptional transfer mechanism.

**Figure 8.**
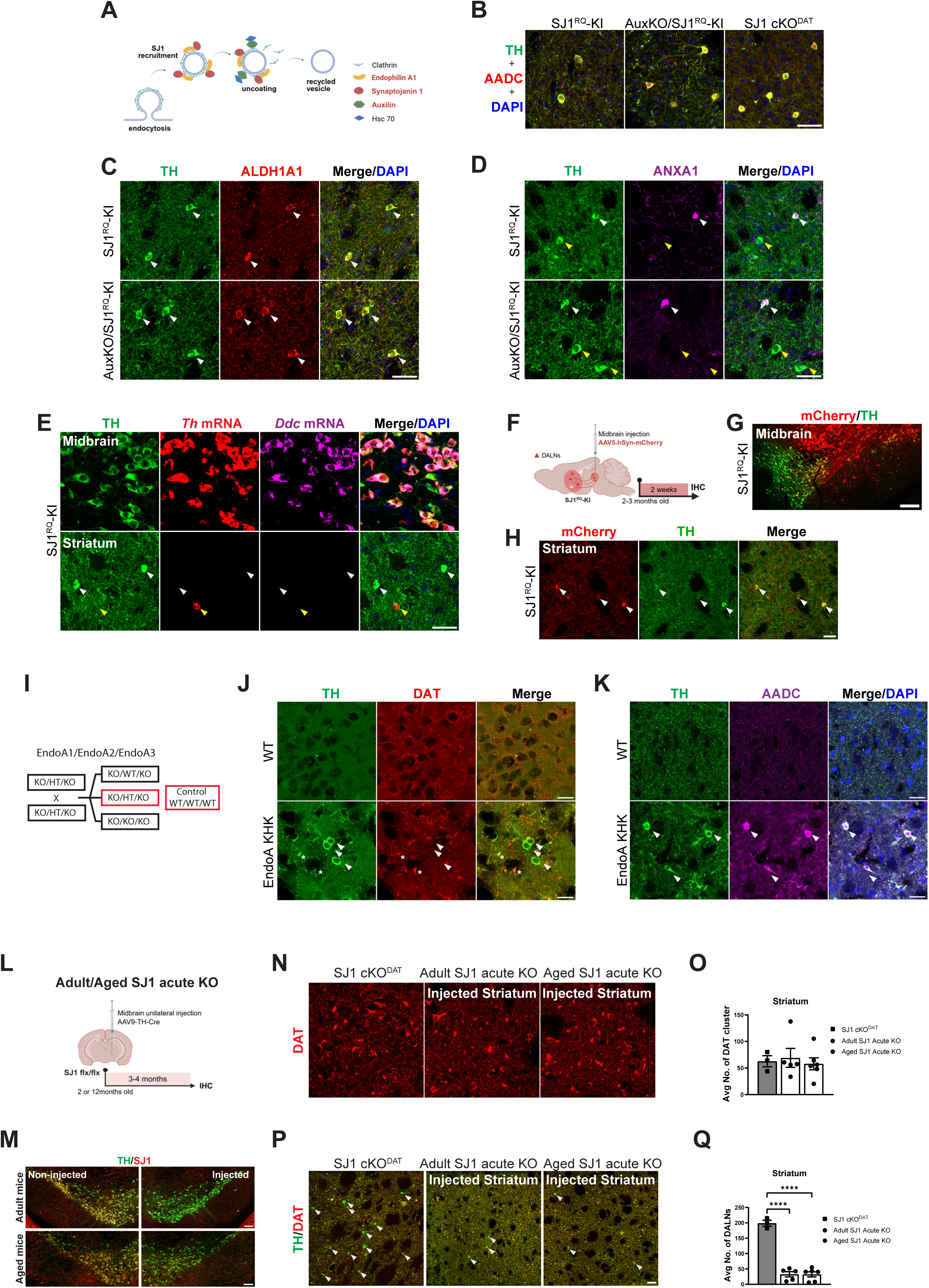
DALN induction is conserved across PD-associated endocytic mutant mice and depends on developmental plasticity (A) Schematic illustration of clathrin-mediated endocytosis. PD-associated endocytic proteins, including, SJ1, Auxilin and Endophilin A1, are involved in clathrin uncoating and synaptic vesicle recycling. (B) Representative TH and AADC immunostaining images of the striatum from SJ1^RQ^-KI, Aux-KO/SJ1^RQ^-KI and SJ1 cKO^DAT^ mice. TH/AADC double-positive DALNs were detected in all three models. Scale bar: 20 µm. (C), (D) Representative TH co-immunostaining with ALDH1A1 (C) or ANXA1 (D) in the striatum of SJ1^RQ^-KI and AuxKO;SJ1^RQ^-KI mice, showing distinct DALN subtypes. Scale bars: 20 µm. (E) Representative RNAscope-IF images of TH protein together with *Th* and *Ddc* mRNAs in the midbrain and striatum of SJ1^RQ^-KI mice. Midbrain DANs expressed TH protein together with *Th* and *Ddc* mRNAs, whereas striatal TH-positive DALNs lacked detectable *Th* and *Ddc* mRNAs (white arrows). WT THINs expressed Th mRNA but lacked detectable TH protein and *Ddc* mRNA (yellow arrows). Scale bar: 20 µm. (F) Schematic showing the experimental procedure and timeline. AAV5-hSyn-mCherry was injected into the ventral midbrain of 2–3-month-old SJ1^RQ^-KI mice. Brains were collected 2 weeks after injection for immunohistochemical analysis. (G) Representative images showing TH immunostaining and mCherry expression in midbrain DANs of SJ1^RQ^-KI mice. Scale bar: 100 µm. (H) Representative images showing TH immunostaining and mCherry expression in the striatum of SJ1^RQ^-KI mice. In addition to dopaminergic fibres, mCherry signals were detected in TH-positive DALNs (white arrows). Scale bar: 20 µm. (I) Breeding strategy for generation of Endophilin A1/2/3 KO/Ht/KO mice. Male and female Endophilin A1/2/3 KO/Ht/KO mice were crossed, and KO/Ht/KO offspring were selected for analysis, with WT mice as the control. (J), (K) Representative striatal immunostaining of Endophilin A1/2/3 KO/Ht/KO mice showing TH/DAT-positive dopaminergic terminal abnormalities (L) and TH/AADC double-positive DALNs (M). White arrows indicate DALNs and white asterisks indicate DAT-positive clusters. Scale bars: 20 µm. (L) Schematic showing unilateral stereotaxic injection of AAV9-TH-Cre virus into the ventral midbrain of 2 months (adult) or 12 months (aged) SJ1 flx/flx mice to obtain SJ1 acute KO in DANs. Animal brains were collected 3-4 months post injection for IHC. (M) TH and SJ1 immunostaining of the adult and aged SJ1 acute KO mice showing SJ1 specific KO in the injected midbrains. Scale bar: 100 µm. (N) DAT immunostaining of the adult and aged SJ1 acute KO mice showing similar clustering phenotype in dorsal (DS) and ventral striatum (VS) as observed in SJ1 cKO^DAT^ mice. Scale bar: 20 µm. (O) No significant difference of the number of DAT clusters between SJ1 cKO^DAT^, adult and aged SJ1 acute KO. The average number of clusters is counted in five randomly selected, 300 × 300-µm regions of interest (ROIs). n=3-4 mice per genotype. (P) Representative images of striatum show comparable TH/DAT-positive clusters but much lesser DALNs in the injected hemisphere of adult and aged SJ1 acute KO mice when compared to the striatum of SJ1 cKO^DAT^ mice. White arrows: DALNs. Scale bar: 50 µm. (Q) Number of DALNs in the adult and aged SJ1 acute KO is significantly lower than SJ1 cKO^DAT^ mice. Number of DALNs was quantified in stitched coronal section of the injected striatum. n=3-4 mice per genotype. Data are represented as mean ± SEM. Statistical analysis: ordinary one-way ANOVA with post hoc Tukey’s test (C, N, P); *p < 0.05, **p < 0.01, ***p < 0.001 and ****p < 0.0001.

Endophilin A is another PD-linked clathrin uncoating protein that forms a functional complex with SJ1^5–7,42^. We therefore examined 9-month-old Endophilin A1/2/3 KO/Ht/KO triple-mutant mice and observed similar and prominent dopaminergic terminal dystrophy in the striatum, accompanied by TH/AADC-positive DALNs (Fig. 8I-K, S14B, C). A subset of these dorsal DALNs also expressed ALDH1A1 and ANXA1, extending the A9-like molecular phenotype to an independent endocytic mutant (Fig. S14D, E). Across models, DALN induction increased with endocytic dysfunction severity, from little or no response in Aux-KO mice, to restricted dorsal induction in SJ1^RQ^-KI, broader induction in Aux-KO/SJ1^RQ^-KI mice, and the most widespread response in Endophilin A mutants and DAN-specific SJ1 cKO mice (Fig. S14F, G). This cross-model pattern supports DALN emergence as a shared consequence of chronic presynaptic endocytic disruption.

To test whether developmental timing influences DALN adaptation, we acutely deleted SJ1 in adult or aged DANs by unilateral AAV9-TH-Cre injection into the ventral midbrain of SJ1 floxed mice (Fig. 8L). Control injections in WT mice showed no nonspecific toxicity to DAN somata or fibers (Fig. S14H, I). Three to four months later, SJ1 was efficiently depleted in ipsilateral DANs (Fig. 8M), and DAT-positive dystrophic clusters appeared throughout the striatum, resembling genetic SJ1 cKO pathology (Fig. 8N, O). However, despite comparable terminal pathology, DALN induction was markedly reduced, reaching only about 10% of that observed in genetic SJ1 cKO mice (Fig. 8P, Q). Robust DALN induction is therefore strongly dependent on developmental plasticity.

Finally, we tested whether DALNs represent a general response to dopaminergic denervation. Acute DAN loss induced by either 6-OHDA or AAV-caspase-3 produced local TH-positive striatal neurons, consistent with previous reports after DA depletion. However, these cells differed from DALNs in SJ1 cKO mice: they showed much weaker TH immunoreactivity, lacked broader dopaminergic protein markers such as AADC and ALDH1A1, and were not labeled in the Ai9;DAT-Cre background (Fig. S15). Thus, acute lesion-induced TH-positive neurons and DALNs represent molecularly distinct responses to dopaminergic dysfunction.

Together, these findings indicate that DALN induction depends on chronic endocytic dysfunction and developmental timing, rather than acute dopamine depletion alone.

## Discussion

This study identifies a previously unrecognized mechanism of striatal compensation to chronic dopaminergic synaptic failure: selective long-range protein transfer from midbrain DANs to striatal DALNs. In SJ1 DA-cKO mice, severe dopaminergic terminal pathology and DA deficiency did not cause immediate circuit collapse, but instead induced DALNs that acquired a dopamine-like protein state without activating a dopaminergic transcriptional program (Fig. 7X). Functionally, DALN adaptation was associated with local DA production, partial motor preservation and increased resistance to acute DA lesions, suggesting a circuit-level buffering response during early PD progression. Thus, SJ1 loss reveals both a primary vulnerability of dopaminergic terminals and a secondary adaptive mechanism that may delay functional decline in PD.

SJ1 has a clear cell-autonomous role in dopaminergic synaptic transmission, but the underlying defect may not simply reflect canonical clathrin-mediated endocytosis failure. Unlike conventional fast synapses, SJ1 cKO dopaminergic terminals showed whorl-like membrane structures rather than obvious clathrin-coated vesicle accumulation, suggesting that neuromodulatory dopaminergic terminals, which signal through slow volume transmission, may rely on additional or alternative membrane-retrieval mechanisms^43,44^. The genotype–phenotype correlation and graded pathology across SJ1, Auxilin and Endophilin A mutant models support chronic clathrin-uncoating disruption as a shared driver of dopaminergic vulnerability, with nigrostriatal projections to the dorsal striatum being particularly sensitive. Complete SJ1 loss also affected ventral striatal projections, indicating that severe endocytic failure can compromise broader DA circuits involved in motivation and reward.

Despite marked presynaptic dysfunction, SJ1 DA-cKO mice retained motor coordination. This dissociation is consistent with DA-specific cKO mouse models of the Ca²⁺ sensor Syt1 and the release site organizer RIM1/2, in which phasic DA release is strongly impaired but overt motor defects are limited^45,46^. Together, these models support the idea that motor coordination can be maintained during early dopaminergic synaptic dysfunction through residual basal DA tone, postsynaptic receptor adaptation and local striatal compensatory plasticity. They may therefore capture a prodromal or early compensatory stage of PD, before extensive DA denervation produces irreversible motor decline.

A striking finding of this study is the robust emergence of DALNs as a distinct adaptive striatal state. DALNs differ from WT THINs by showing strong TH accumulation, AADC co-expression, circuit-patterned acquisition of nigrostriatal markers such as ALDH1A1 and ANXA1, and reduced intrinsic excitability. These features suggest two complementary modes of compensation: local DA production through acquired dopaminergic enzymes and reduced inhibitory interneuron output that may disinhibit downstream striatal targets and enhance residual DA signaling. DALN induction was robust in developmental SJ1 cKO mice and shared across PD-linked endocytic mutant models, but was markedly reduced after adult SJ1 deletion and not reproduced by acute DA lesions. Thus, DALNs are not simply a response to DA depletion, but a conserved, developmentally constrained plasticity program induced by chronic presynaptic endocytic stress.

Our data strongly support that DALNs acquire a DA-like state predominantly via long-range protein transfer rather than transcriptional reprogramming. DALNs contained dopaminergic proteins but lacked the corresponding mRNAs; DAN ablation caused progressive loss of these proteins without eliminating DALNs; and viral tracing showed that DAN-derived proteins appear in striatal DALNs. The transfer pathway is selective, favoring soluble proteins while restricting membrane, nuclear and organelle-associated cargoes. Together with the absence of detectable microglial or astrocytic activation (Fig. S3A–C), this cargo selectivity argues against nonspecific leakage or glia-mediated clearance of cellular debris, and instead supports a regulated inter-neuronal transfer process.

Importantly, DA was detected in DALNs and was further increased by L-DOPA treatment, indicating that transferred TH and AADC can support local DA production. This concept is consistent with PD gene therapy strategies designed to restore striatal DA synthesis^47,48^. Although vMAT2 was not detected, non-vesicular DA release has been proposed in DANs^49^. Directly testing DA release from DALNs remains challenging because they are sparse and lack a selective genetic handle. Future DALN-specific tools will be needed to determine their contribution to striatal DA tone. This DAN–DALN transfer pathway may also contribute to early neuroprotection while carrying potential pathological risk. SJ1 cKO mice were more resistant to acute DA lesions, and detection of transferred caspase-3 in DALNs raises the possibility that transfer redistributes stress-related cargo away from vulnerable DANs. However, the same route also conveyed disease-associated proteins, including α-synuclein and tau, suggesting that an initially adaptive protein-sharing mechanism may later enable pathological propagation^50–52^.

The present data define the phenomenon, its selectivity and its functional consequences, but do not yet identify the cellular route of transfer. Possible mechanisms include endocytic uptake, nanotube-like contacts^51,53–55^, extracellular vesicles^50,56,57^ or exchange sites created by dystrophic terminals^22,58^. Finally, TH-positive neurons have been reported in human striatum and are altered in PD, but whether they correspond to DALNs remains unclear^29–31^. Human spatial transcriptomics and protein-level mapping will be important for assessing clinical relevance.

## Methods

### Animals

All experiments involving animals were conducted according to the protocol approved by SingHealth Institutional Animal Care and Use Committee (IACUC). Mice were housed in SPF rooms with a 12-hour light/dark cycle. Aux-KO, SJ1^RQ^-KI and Aux-KO/SJ1^RQ^-KI mouse line was generated as described previously^21,22^ and SJ1-KI^RQ^ mouse line was a gift from Prof Pietro De Camili (Yale University). SJ1 flx/flx (C57BL/6J-Synj1^em1cyagen^; S-CKO-00357, Cyagen) was generated by inserting loxP sites flanking exon 4 of Synj1 using CRISPR/Cas9. These mice were crossed with DAT-IRES-Cre (C001024, Cyagen) and TH-IRES-Cre knock in line (EM:00254, European Mouse Mutant Archive; kindly provided by Dr Sarah Luo, IMCB-A*STAR) to obtain double heterozygous mice, respectively. To create DAN-specific SJ1 conditional KO mice, double heterozygous mice were crossed with homozygous SJ1 flx/flx, obtaining DAT-Cre^+/-^;SJ1 flx/flx and TH-Cre^+/-^; SJ1 flx/flx mice, referred to as SJ1 cKO^DAT^ and SJ1 cKO^TH^, respectively. All mice used in the experiments with Cre-dependant promoter were heterozygous (+/-), simply referred as DAT-Cre and TH-Cre in the manuscript. Appropriate littermate or age-matched controls (SJ1 flx/flx or DAT-Cre;SJ1 +/flx or TH-Cre;SJ1 +/flx) were used in all experiments. Specifically, DAT-Cre;SJ1 +/flx mice were used as control for western blot, acute slice DA release assay and drug-induced open field test.

Ai9 reporter mice (JAX #007909, RRID: IMSR_JAX:007909) were kindly provided by Prof Enrico Petretto (Duke-NUS) for specific labelling following Cre-mediated recombination. Homozygous Ai9 mice were first crossed with homozygous SJ1 flx/flx to obtain double heterozygous, Ai9^+/-^;SJ1 +/flx mice. Subsequent models involving Ai9 were all heterozygous Ai9^+/-^, hence simply referred as Ai9 in the manuscript. Ai9;SJ1 +/flx mice were then crossed with SJ1 cKO^DAT^ and SJ1 cKO^TH^ to obtain Ai9;SJ1 cKO^DAT^ and Ai9;SJ1 cKO^TH^, respectively. Appropriate littermate or age-matched controls were used in all experiments.

Endophilin-A1 knockout/A2 heterozygous/A3 knockout mice (EndoA1^−/−^; EndoA2^+/−^; EndoA3^−/−^, hereafter EndoA1KO/A2HT/A3KO) on a C57BL/6J background were generated as previously described^5^. Experimental EndoA1KO/A2HT/A3KO mice were generated by crossing EndoA1^−/−^; EndoA2^+/−^; EndoA3^−/−^ breeders. 9 months old male and female mice were included in the study. Animals were group-housed in under a 12:12-h light/dark cycle, with controlled temperature and humidity, and were provided standard laboratory chow and water ad libitum. Nesting material, shelters and environmental enrichment were provided. All experimental procedures were reviewed and approved by the CNC-UC Animal Welfare Body (ORBEA 320). Experiments were conducted in accordance with European Directive 2010/63/EU and Portuguese Decree-Law no. 113/2013 concerning the protection of animals used for scientific purposes. Animals were maintained at the licensed CNC-UC animal facility, and all procedures were performed by trained investigators accredited by the Portuguese authorities.

### Stereotaxic injections

Mice were anesthetized with isoflurane (5% induction, 2% maintenance at 0.5 to 0.8 L/min oxygen or air flow) and received unilateral stereotaxic injections into the right ventral midbrain (anteroposterior, AP −3.3 mm, mediolateral, ML 1.2 mm, and dorsoventral, DV −4.3 mm relative to bregma), unless stated otherwise.

For SJ1 acute KO mouse model, AAV9-TH-Cre (107788-AAV9, Addgene; 1 μL) was injected to SJ1 flx/flx mice, and brain tissues were collected 3 to 4 months later.

For genotype-dependent lesion resistance studies, SJ1 cKO^DAT^ and DAT-Cre;SJ1 +/flx control mice received unilateral right ventral midbrain lesions. Casp3-mediated ablation was induced by injection of AAV5-FLEX-Caspase3 (45580-AAV5, Addgene; 3.5 × 10¹² vg/mL, 1 μL; low-dose Casp3), whereas 6-hydroxydopamine (6-OHDA, #H4381, Sigma-Aldrich) lesions were induced by injection of 6-OHDA (3 μg in 1 μL of 0.2% ascorbate/saline; H4381, Sigma-Aldrich). DAN survival, postoperative survival, and rotational behaviour were subsequently assessed.

For examination of the dependence of dopaminergic protein content in DALNs on intact midbrain dopaminergic innervation, SJ1 cKO^DAT^ mice received high-dose AAV5-FLEX-Caspase3 (7 × 10¹² vg/mL, 1 μL). Brain tissues were collected at baseline and 2, 4, and 6 weeks after injection, dopamine immunostaining was performed at 2 weeks, and spontaneous rotational behaviour was assessed at baseline and 2 and 4 weeks.

For acute dopaminergic lesion models, C57BL/6J mice were used for 6-OHDA lesions, whereas DAT-Cre or TH-Cre heterozygous mice were used for Casp3-mediated ablation. Mice receiving 6-OHDA were pretreated with desipramine (25 mg/kg, i.p.; D3900, Sigma-Aldrich) 30 minutes before surgery, followed by unilateral injection of 6-OHDA (2.88 μg in 0.2% ascorbate/saline). Casp3-mediated ablation was induced by unilateral injection of AAV5-FLEX-Caspase3 (3.5 × 10¹² vg/mL, 1 μL) injected into the right ventral midbrain of Cre-heterozygous mice. Brain tissues were collected 2 weeks post injection.

For AAV-mediated protein-transfer experiments, Cre-independent or Cre-dependent viral vectors were unilaterally injected into the right ventral midbrain or the right striatum (AP 0.2 mm, ML –2.0 mm, DV −2.5 and −4.0 mm). Unless otherwise stated, 1μL of virus was delivered per animal, and brain tissues were harvested at 2 or 4 weeks post injection. Cre-independent vectors included AAV5-hSyn-mCherry (114472-AAV5, Addgene), AAV2-mTH-EGFP (BC-0319, BrainCase), and AAV9-CMV-hSNCA-EGFP (BC-3422/9, BrainCase). Cre-dependent vectors included AAV5-FLEX-tdTomato (28306-AAV5, Addgene), AAV5-DIO-ChR2-EYFP (35509-AAV5, Addgene), AAV2/9-DIO-Caspase3-P2A-EGFP (PT-1230, BrainVTA), AAV2/9-DIO-hM3Dq-EGFP (PT-0891, BrainVTA), AAV2/9-DIO-H2B-EGFP (PT-0258, BrainVTA), AAV2/9-DIO-tau-mCherry (PT-0527, BrainVTA), AAV2/9-DIO-tau(P301L)-mCherry (PT-1601, BrainVTA), AAV2/9-DIO-mito-EGFP (PT-8144, BrainVTA), and AAV2/9-DIO-synaptophysin-mCherry (PT-2755, BrainVTA). Reporter expression in midbrain DANs and the appearance of the encoded proteins in striatal DALNs were assessed by immunohistology.

For in vivo induction of α-synuclein pathology, active mouse recombinant α-synuclein preformed fibrils (Type 1; SPR-324B, StressMarq Biosciences; 2 mg/mL) were thawed at room temperature and fragmented before using a QSonica Q700 sonicator (700 W, 20 kHz) at 80% amplitude with alternating 30-s ON and 30-s OFF cycles for 30 cycles, while maintaining the bath temperature at approximately 10 °C. Sonicated fibrils (1 μL; 2 μg) were stereotaxically injected into the right ventral midbrain. Brain samples were collected one month after injection and analysed for α-synuclein pathology.

### Motor Behavioral Tests

Mice between 2-8 months old were used for balance beam and rotarod tests. The mice were divided into two age groups: 2-3 months old (n= 13 for Ctrl, n=15 for SJ1 cKO^DAT^) and 6-8 months old (n=9 for Ctrl and n=10 for SJ1 cKO^DAT^). Both males and females were used for the behavioral assays. Both tests were conducted during the light period. The mice were allowed to be acclimatized to the test for at least 30 min before each test.

#### Balance Beam Test

A narrow beam (10 mm width) was suspended 15 cm above a soft padding. Mice were placed on end of the beam and during their trip towards the other end, the number of missteps (paw slips) was recorded. The test was conducted thrice for each mouse, and the average number of missteps was calculated.

#### Rotarod Test

Mice were placed on a rod which is rotating at 4 rpm with an acceleration to 40 rpm within 6 min. The duration of the mice staying on the rod is measured three times at 10 min intervals and the average ‘latency to fall’ was determined for each mouse.

#### Open Field Test (OFT)

Mice were placed in a chamber measured 40 cm (length) x 40 cm (width) x 30 cm (height) and recorded for 15 min. Recorded videos were analyzed with TopScan video tracking software. The mice path tracking and time spent in the defined area were analyzed. The mice used for two age groups: 2-3 months old (n= 7) and 6-8 months old (n= 6), unless otherwise stated.

For drug treatment open field test, after 15 min baseline recording, intraperitoneal injections of amphetamine at 3.0 mg/kg (D961, Age D’or) or SKF81297 at 2.5mg/kg (S143, Sigma-Aldrich) or quinpirole at 0.2mg/kg (sc-253339, Santa Cruz) were administered, dosage was optimized with reference to literature^46^. Total traveled distance of the mice after 60 min injection was recorded by the tracking system and the results were normalized to individual baseline. Traveled distance was expressed by percent of baseline. Percent of baseline was calculated per animal as (post_mean / baseline_mean) × 100, where baseline_mean is the mean distance over the 15-min pre-drug period and post_mean is the mean distance over the 60 min post-drug test.

#### Spontaneous rotation test

Spontaneous rotational behaviour was assessed before lesion and at 2 and 4 weeks after unilateral Casp3 or 6-OHDA injection. During the dark phase, mice were individually placed in a glass cylinder and recorded for 10 min under dim illumination. Full-body ipsilateral and contralateral 360° rotations were counted manually. Data were expressed as the percentage of contralateral rotations relative to total rotations.

### Antibodies

The antibodies used in this study were obtained from commercial sources as stated: mouse anti-α-synuclein (610786, RRID: AB_398107), mouse anti-AP2 (611350, RRID: AB_398872) and mouse anti-Dynamin (610245, RRID: AB_397640) from BD Biosciences; goat anti-ChAT (AB144P, RRID: AB_2079751), mouse anti-clathrin light chain (AB9884, RRID: AB_992745), rat anti-DAT (MAB369, RRID: AB_2190413), mouse anti-SV2C (MABN367, RRID: AB_2905667) and rabbit anti-TH (AB152, RRID: AB_390204) from Merck Millipore; mouse anti-β-actin (sc-47778, RRID: AB_2714189) and mouse anti-BIN1 99D (sc-13575, RRID:AB_626753) from Santa Cruz Biotechnology; mouse anti-NeuN (MAB377, RRID: AB_2298772), rabbit anti-ALDH1A1 (HPA002123, RRID: AB_1844722), rabbit anti-GFAP (ZRB2383, RRID: AB_2905668) and rabbit anti-SJ1 (HPA011916, RRID: AB_1857692) from Sigma-Aldrich; rabbit anti-Endophilin 1 (159002, RID: AB_887757), mouse-anti Calretinin (214111, RRID: AB_2619904), mouse anti-TH (213211, RRID: AB_2636901), rabbit anti-Amphiphysin 1 (120002, RRID: AB_887690), rabbit anti-Synaptogyrin 3 (103302, RRID:AB_2619752), rabbit-anti Synaptophysin 1 (101002, RRID:AB_887905), rabbit anti-AADC (369003, RRID: AB_2620131), guinea pig-anti Calbindin D28k (214005, RRID: AB_2619902), mouse anti-Parvalbumin (195011, RRID: AB_2619883) and rabbit anti-D2R (376 203, RRID: AB_2636918) from Synaptic Systems; rabbit anti-vMAT2 (MSFR106400, RRID: AB_2571857) from Nittobo Medical; rabbit anti-DARPP-32 (2306, RRID: AB_823479) and rabbit anti-Neuropeptide Y (11976, RRID: AB_2716286) from Cell Signaling Technology; rabbit anti-Iba1 (019-19741, RRID: AB_839504) from FUJIFILM Wako Pure Chemical Corporation; goat anti-ALDH1A1 (AF5869_RRID: AB_2044597) from R&D Systems; rabbit anti-alpha synuclein phospho S129 (EP1536Y, RRID:AB_765074) from abcam; rabbit anti-DRD1 (17934-1-AP, RRID:AB_10598308), mouse anti-GAPDH (60004-1, RRID: AB_2107436) and rabbit anti-GAPDH (10494-1-AP, RRID: AB_2263076) from Proteintech; rabbit anti-Annexin A1 (71-3400, RRID:AB_88080) from Thermofisher; rabbit anti-dopamine (IS1005) from ImmuSmol; and rabbit anti-Tyrosine Hydroxylase (Ser40) (p1580-40, RRID:AB_2492279) from phosphosolutions. The rabbit anti-Synapsin was obtained from Prof Pietro De Camilli’s lab at Yale University.

Secondary antibodies used were all purchased from commercial sources as stated: donkey anti-mouse IgG (H + L) Alexa Fluor Plus 488 (A32766, RRID: AB_2762823), donkey anti-mouse IgG (H + L) Alexa Fluor 594 (A21203, RRID: AB_2535789), donkey anti-mouse IgG (H + L) Alexa Fluor Plus 647 (A32787, RRID: AB_2762830), donkey anti-rabbit IgG (H + L) Alexa Fluor Plus 488 (A32790, RRID: AB_2762833), donkey anti-rabbit IgG (H + L) Alexa Fluor Plus 594 (A32754, RRID: AB_2762827), donkey anti-rabbit IgG (H + L) Alexa Fluor 647 (A32795, RRID: AB_2762835), donkey anti-rat IgG (H + L) Alexa Fluor Plus 488 (A48269, RRID: AB_2893137), donkey anti-rat IgG (H + L) Alexa Fluor 594 (A21209, RRID: AB_2535795), donkey anti-goat IgG (H + L) Alexa Fluor 488 (A11055, RRID: AB_2534102), donkey anti-goat IgG (H + L) Alexa Fluor 594 (A32758, RRID: AB_2762828), goat anti-guinea pig IgG (H + L) Alexa Fluor 488 (A11073, RRID: AB_2534117) and goat anti-guinea pig IgG (H + L) Alexa Fluor 647 (A21450, RRID: AB_2535867) from Life Technologies; goat anti-mouse IgG light chain HRP (AP200P, RRID:AB_805324) and mouse anti-rabbit IgG light chain HRP (MAB201P, RRID:AB_827270) from Merck; IRDye 800CW donkey anti-rabbit IgG (926-32213, RRID: AB_621848), IRDye 800CW donkey anti-mouse IgG (926-32212, RRID: AB_621847), IRDye 680RD donkey anti-mouse IgG (926-68072, RRID: AB_10953628) and IRDye 800CW goat anti-rat (926-32219, RRID: AB_1850025) from LI-COR Biosciences.

### Brain Histology and Immunofluorescence

Mice were anesthetized with a Ketamin/Xylazine anesthetic cocktail injection, perfused transcardially with ice-cold 4% paraformaldehyde (PFA) in 1 X phosphate-buffered saline (PBS) and the brains were kept in the same fixative overnight at 4°C. On the next day, brains were transferred to 30% sucrose in 1 X PBS and kept overnight at 4°C on top of a roller. Brains were then embedded in OCT (Tissue-Tek) and freeze in liquid nitrogen-cooled 2-methylbutane (Isopentane). Coronal or sagittal sections of 20 µm thickness were cut with a cryostat. The sections were either mounted on adhesive slide, SuperFrost® Plus (VWR) or collected as floating sections in 1 X PBS in a 24-well plate. Sections were then blocked in 5% (bovine serum albumin) BSA and 0.1% Triton X-100 in 1 X PBS for 1 hour at room temperature. After blocking, sections were incubated with primary antibodies and kept overnight at 4°C. Subsequently, sections were washed 3 times for 10 min with 0.1% Triton X-100 in 1 X PBS, then incubated with Alexa-conjugated secondary antibodies for 1 hour at room temperature. Finally, the sections were mounted with Fluoromount-G (Invitrogen) and sealed with nail polish. The floating sections were mounted onto slides after all the staining procedures. Images were acquired with Andor BC43 (Oxford Instruments) with 10x, 20x and 40x oil objective or a spinning disk system (Gataca Systems) based on an inverted microscope (Nikon Ti2-E; Nikon) equipped with a confocal spinning head (CSU-W, Yokogawa) and a Plan-Apo 20x and 40x oil objective.

For stereological analysis of TH-positive DANs in the midbrain, 30 µm coronal section were collected incubated with 0.1% H2O2 for 5 min to quench endogenous peroxidase activity and subsequently incubated with buffer containing 5% BSA, and 0.1% Triton X-100 in 1X PBS for 1 hour at room temperature. Next, sections were incubated overnight at 4°C with anti-TH primary antibody.The sections were then washed with 1X PBS three times for 10 min and incubated with a biotinylated anti-rabbit secondary antibody (Vector Laboratories, USA) for 1 hour at room temperature, later followed by incubation with Avidin/Biotin complex (ABC) reagent (Vector Laboratories, USA) at room temperature. Finally, immunoreactivity was revealed by incubation with diaminobenzidine (DAB). Stereological analysis was performed using the optical fractionator probe in the Stereo Investigator (MBF Bioscience) software. A total of 7-9 coronal sections of every fourth sections were collected across the midbrain and used for counting. The SNc and VTA regions were outlined based on the Allen mouse brain atlas and counting was performed using 63x oil lens. The parameters used were guard zone of 2 μm, dissector height of 13 μm and counting frame of 200 μm x 200 μm.

For dopamine immunostaining, brain tissues were processed using the STAINperfect Immunostaining Kit A (SP-A-1000; ImmuSmol) together with a rabbit anti-dopamine antibody (1:500) (IS1005; ImmuSmol), according to the manufacturer’s instructions. Sections were subsequently co-stained with the indicated protein markers and imaged by confocal microscopy. Where indicated, mice received benserazide (15 mg/kg, i.p.) 30 min before L-DOPA administration (30 mg/kg, i.p.) to inhibit peripheral L-DOPA metabolism, and brain tissues were collected 60 min after L-DOPA injection to assess dopamine production and AADC activity in striatal DALNs.

### RNAscope combined with Immunofluorescence

RNAscope fluorescence in situ hybridization combined with immunofluorescence was performed using the RNAscope Multiplex Fluorescent Reagent Kit v2 and Integrated Co-Detection workflow (ACD) according to the manufacturer’s instructions with minor modifications. Fixed-frozen mouse brains were cryosectioned at 10 μm, mounted on Superfrost Plus slides, and stored at −80 °C. Sections were baked, post-fixed in 4% PFA, dehydrated through graded ethanol, treated with hydrogen peroxide, and subjected to target retrieval. Primary antibodies were applied overnight at 4 °C, followed by post-fixation and Protease Plus treatment. RNAscope probes were then hybridized and amplified according to the ACD protocol, after which probe signals were developed using TSA fluorophores. Sections were incubated with fluorescent secondary antibodies, mounted with Fluoromount-G with DAPI (Invitrogen) and sealed with nail polish, and imaged by Andor BC43 (Oxford Instruments) with ×10 air and ×40 oil-immersion objectives. All probes were obtained from ACD.

The following mouse probes were used: Th-C1 (317621), Aldh1a1-C2 (491321-C2), Ddc-C2 (318681-C2), Slc6a3-C2 (315441-C2), tdTomato-C2 (317041-C2), and Anxa1-C3 (509291-C3). Signals from Polr2a and Ppib in the ACD positive-control probe set were used to assess RNA integrity, while a negative-control probe targeting bacterial dapB was used to evaluate nonspecific background.

### Quantitative Image Analysis

All quantitative analyses were performed using Fiji software (Version ImageJ 2.160/1.54p/Java 1.8.0_322 (64-bit), RRID: SCR_002285). Image acquisition parameters were kept constant for experimental groups.

#### Quantification of SJ1 Expression in DANs

TH-positive DANs were identified by manual intensity thresholding of the TH channel followed by size filtering to exclude particles smaller than 60 µm², generating neuronal region of interests (ROIs). These ROIs were applied to the SJ1 channel after background subtraction. Mean fluorescence intensity of SJ1 within TH-positive ROIs was quantified and averaged per animal to yield a single mean intensity for the SNc and VTA. For each experiment, control values were normalized to 100%, and SJ1 cKO^DAT^ values were expressed relative to control mean. At least three mice from each genotype were collected for this purpose.

#### Quantification of DAT Immunoreactivity Clusters

For each mouse, 5 random sampling sites were selected respectively for both dorsal and ventral striatum. The threshold setting for the clusters was set to quantify puncta bigger than 5 μm^2^ to ensure normal positive axonal terminals were not counted in. The average size of clusters and average number of clusters was counted automatically with Fiji plugin “Analyse Particles” based on all five random sampling sites for each mouse. Density of clusters was calculated as the number of clusters per ROI area and normalized to a standard 300 × 300 µm² area. At least three mice from each genotype and each age group were collected for this purpose.

#### Colocalization Analysis

Colocalization between different markers was assessed using Pearson’s correlation coefficient (PCC). PCC values were calculated using the JaCoP plugin in Fiji. Mean PCC values were obtained from 3–7 sections per animal and averaged for analysis.

#### Glia Quantification

Iba1-positive microglial and GFAP-positive astrocytic area fractions were quantified in striatal sections using custom macros implemented in Fiji and executed in batch mode. Images were first converted to 8-bit grayscale and median-filtered (radius = 1 pixel) to reduce background noise while preserving cellular morphology. Binary masks were generated by automated thresholding using the *Triangle* method for Iba1-stained images and the *Yen* method for GFAP-stained images. Following thresholding, despeckling was applied to remove isolated noise artifacts. The percentage area fraction of positive staining was then calculated from the resulting binary masks. At least three mice from each genotype were collected for this purpose.

#### Quantification of RNAscope Signals

RNAscope signals were quantified in Fiji within manually defined neuronal ROIs based on the corresponding immunofluorescence signals and cellular morphology. Discrete transcript puncta were counted directly within each cellular ROI. For densely clustered signals in which individual puncta could not be reliably resolved, the integrated fluorescence intensity of the cluster was divided by the mean intensity of isolated single puncta to estimate the corresponding puncta number. The number of transcript puncta per cell was quantified, and measurements from multiple cells were averaged for each animal to generate a single animal-level value for statistical analysis.

#### Quantification of Marker Co-expression

Marker co-expression was assessed within manually identified cellular ROIs. For immunofluorescence and viral reporter signals, cells were classified as positive when the mean fluorescence intensity within the cellular ROI exceeded the mean background intensity by at least three standard deviations. For RNAscope signals, cells containing at least two discrete transcript puncta within the corresponding cellular ROI were classified as transcript-positive. Cells meeting the positivity criteria for both indicated markers were classified as double-positive, whereas cells meeting the criterion for only one marker were classified as single-positive. For each section, the percentage of reference marker-positive cells that were also positive for the second marker was calculated. Percentages from multiple sections were averaged for each animal to generate a single animal-level value for statistical analysis.

#### Quantification of Protein and Dopamine Signals in Striatal DALNs

For quantification of protein and dopamine signals in striatal DALNs, individual TH-positive somata were manually delineated in Fiji to generate cellular ROIs. Mean fluorescence intensities were measured within these ROIs after subtraction of local background fluorescence. For viral reporter-transfer experiments, transfer efficiency was calculated as the mean reporter fluorescence intensity in DALNs divided by the mean reporter fluorescence intensity in midbrain DANs and expressed as a percentage. Measurements from all analysed cells were averaged for each animal to generate a single animal-level value for statistical analysis.

#### Quantification of Midbrain DAN Survival and the intensity of DAT-positive DA fibers

TH-positive midbrain DANs were quantified in matched coronal sections from the lesioned and unlesioned hemispheres. Neuronal survival was expressed as the number of TH-positive neurons on the lesioned side relative to the contralateral side. Striatal DAT-positive innervation was quantified by measuring mean DAT fluorescence intensity within matched striatal ROIs after local background subtraction. DAT positive terminal integrity was expressed as the fluorescence intensity on the lesioned side relative to the contralateral side. Values from multiple sections were averaged for each animal to generate a single animal-level value for statistical analysis.

### Quantification of Striatal neurons

Quantification of TH-positive and tdTomato-expressing cells were conducted using Fiji software (Version ImageJ 2.160/1.54p/Java 1.8.0_322 (64-bit), RRID: SCR_002285) with full stitched half coronal section, unless stated otherwise. Image acquisition involved tiling of the entire section with either 10x or 20x objective lens on Andor BC43. Cell Counter plugin in Fiji software was used to count visible cell bodies across the striatum (outlined based on the Allen Mouse Brain Atlas). Cell count from individual channels was used for percentage ratio quantification. For analyses comparing the dorsal and ventral striatum, regional boundaries were defined according to the Allen Mouse Brain Atlas on the full stitched coronal section prior to cell quantification. Quantification of TH-positive cells overlapping with other marker of interest were counted in three to four randomly selected striatum ROIs.

Quantification for TH-positive cells intensity in lesioned and genetic mouse models was done using Fiji software. A consistent threshold was applied across all images to isolate TH-positive signals. The 8-bit figure type was used to measure the mean density of the TH signal. Background fluorescence was determined from adjacent areas without specific staining. All measurements were performed under identical settings to ensure consistency across experimental groups. For each mouse, six random TH-positive cells were selected from both dorsal and ventral striatum. At least three mice from each condition were collected for this purpose.

Quantification for tdTomato expression in Ai9;SJ1 cKO^TH^ mouse model for WT THINs and DALNs was quantified from stitched coronal striatum images. Cells were classified as either tdTomato-positive only or tdTomato/TH double-positive based on TH immunoreactivity. For each animal, five random ROIs containing tdTomato signal were selected from the stitched image for analysis. Cells were identified by manual intensity thresholding of the tdTomato channel followed by size filtering to exclude particles smaller than 60 µm², generating neuronal ROIs. Mean fluorescence intensity of tdTomato within ROIs was quantified and averaged per animal to yield a single mean intensity. Fluorescence intensity was measured in Fiji using corrected total cell fluorescence (CTCF), calculated as integrated density minus the product of cell area and mean background fluorescence. Background fluorescence was measured from a cell-free area within the same stitched image, and the same background value was applied to all cells quantified from that stitched image. A total of four mice were used for quantification.

Long soma diameter for WT THINs and DALNs was quantified in the same brain sections collected from Ai9;SJ1 cKO^TH^ mouse model. Using the same ROIs above, the diameter of each cell was measured in Fiji by tracing the longest axis of the soma. For statistical analysis, individual cell measurements were averaged per mouse, and the mean long soma diameter for each subpopulation was calculated. At least three mice were used for this quantification.

### Electron Microscopy

2 months old mouse was anesthetized and fixed by transcardial perfusion with 4% PFA and 0.125% glutaraldehyde (EMS, Hatfield, PA) in 0.1 M PB buffer. Brain was removed and dissected in small pieces (1□×□1□×□1□mm^3^) and further incubated in 2.5% glutaraldehyde in 0.1 M PB buffer at 4□°C overnight. The samples were then washed for 5 min in 1 X PBS for three times. The samples were post-fixed with 1% osmium tetroxide for 1 hour and washed with distilled water. They are then dehydrated through an ethanol series (25%, 50%, 75%, 95% and 100%) and absolute acetone, before infiltrating with araldite resin/acetone mixtures. On the next day, samples were further infiltrated in pure resin at 40°C to 50°C, then embedded in resin and polymerised at 60°C for 24 hours. The embedded sample was sectioned into 100nm slices with a Leica UC6 ultramicrotome and stained with lead citrate. Images were captured with JEOL JEM-1400Flash TEM, operating at 100kV.

### Immunoblotting

Striatum tissues were homogenized in buffer containing 20 mM Tris-HCl (pH 7.4), 150 mM NaCl, 2 mM EDTA and cOmpleteTM EDTA-free Protease Inhibitor Cocktail (Roche, USA). The homogenized samples were centrifuged at 700 g for 10 min to obtain the post-nuclear supernatant (PNS). Protein concentration was determined by the Pierce BCA Protein Assay Kit. SDS-PAGE and western blotting were performed by standard procedures. The membranes were blotted with primary antibodies and then IRDye® Secondary Antibodies (LI-COR, USA) or HRP light chain secondary antibodies. The protein bands were detected using Odyssey ® CLx imaging system or Bio-Rad ChemiDoc Touch Imaging System, and its intensity was quantified via Image Studio. A minimum of three independent sets of samples were used for quantification.

### HPLC

Striatum tissues dissected from mice brain were homogenized in 0.5 N perchloric acid with 100 µM of deferoxamine mesylate and 100 µM of glutathione. The homogenized samples were further sonicated, centrifuged and the supernatants were filtered using 0.1 µm PVDF centrifugal filters before collecting the filtrates for HPLC analysis. A reversed-phase UltiMate 3000 HPLC system (Thermo Fisher Scientific) with an electrochemical detector and a reversed-phase column (Vydac Denali, C18, 4.6 x 250 mm, 5 µm particle size, 120 Å pore size) were used to run the samples. The HPLC run was performed at a flow rate of 1 ml per minute with a mobile phase containing 1.3% sodium acetate, 0.01% EDTA (pH8.0), 0.5% sodium 1-heptanesulfonate, 7% acetonitrile (v/v) and 2% methanol (v/v), adjusted to pH 4.0 with acetic acid. All buffers used for HPLC analysis were double filtered through 0.2 µM nylon membranes. Dopamine, DOPAC and HVA levels in the samples was identified by retention time of their respective standard and quantified by measuring the area under the peak using the software ChromeleonTM 7.2 Chromatography Data System (Thermo Fisher Scientific). The areas under the peaks were normalized to their respective tissue weight.

### Acute Brain Slice Preparation

Mouse brains were rapidly removed after decapitation and placed in high-sucrose ice-cold oxygenated artificial cerebrospinal fluid (ACSF) containing the following (in mM): 125 NaCl, 25 NaHCO_3_, 10 glucose, 2.5 KCl, 2.5 MgCl_2_, 0.5 CaCl_2_, 1.24 NaH_2_PO_4_, 1.68 HEPES, pH 7.4 and 95% O_2_/5% CO_2_. Coronal brain slices were cut at a thickness of 250 μm using a vibratome (VT1200S, Leica Biosystems) and immediately transferred to an incubation chamber filled with ACSF containing the following (in mM): 125 NaCl, 25 NaHCO_3_, 10 glucose, 2.5 KCl, 2 MgCl_2_, 1 CaCl_2_, 1.24 NaH_2_PO_4_, 1.68 HEPES, pH 7.4 (300–310 mOsm), equilibrated with 95% O_2_ and 5% CO_2_. Slices were allowed to recover at room temperature at least for 60 min.

For KCl-evoked and AMPH-induced release, striatum brain slices were incubated with 300 μL pre-warmed ACSF containing the following (in mM): 125 NaCl, 25 NaHCO_3_, 1.25 NaH_2_PO_4_.2H_2_O, 2.5 KCl, 1.8 CaCl_2_, 1 MgCl_2_, 0.4 ascorbic acid, pH 7.4 (300–310 mOsm) at 37 °C, 5 min. The incubation medium in this step served as the baseline control for subsequent measurements. Another 300μL ACSF containing 40 mM KCl or 10 μM amphetamine (AMPH; d-(+)-Amphetamine sulfate, D961, Age D’or) were added to incubated with the same striatum brain slices at 37 °C, 5 min. The incubation medium in this step served as the KCl-evoked or AMPH-induced DA released samples from striatum of control (DAT-Cre;SJ1 +/flx) and SJ1 cKO^DAT^ mice for subsequent HPLC detecting. After adjusting the final concentration of the ACSF buffer to 0.1 N perchloric acid, the DA released samples was filtered through 0.1 µm PVDF centrifugal filters and subjected to HPLC analysis using the same parameters as those used for striatum tissues detection. The areas under the peaks were normalized to their respective protein concentration.

### Electrophysiology

#### Brain slice preparation

Mouse brains were rapidly removed after decapitation and placed in high-sucrose ice-cold oxygenated artificial cerebrospinal fluid (ACSF) containing the following (in mM): 250 sucrose, 26 NaHCO_3_, 10 glucose, 2.5 KCl, 4 MgCl_2_, 0.1 CaCl_2_, 1.25 NaH_2_PO_4_.2H_2_O, 3 Myo-Inositol, 2 Na-pyruvate, 0.5 ascorbic acid,1 kyneurenic acid, pH 7.4 and 95% O2/5% CO_2_. Coronal brain slices were cut at a thickness of 250 μm using a vibratome (VT1200S, Leica Biosystems) and immediately transferred to an incubation chamber filled with ACSF containing the following (in mM): 125 NaCl, 25 NaHCO_3_, 1.25 NaH_2_PO_4_.2H_2_O, 2.5 KCl, 1.8 CaCl_2_, 1 MgCl_2_, pH 7.4 (300–310 mOsm), equilibrated with 95% O2 and 5% CO_2_. Slices were allowed to recover at 32 °C for 30 min and then maintained at room temperature.

#### Whole-cell patch-clamp recordings and data analysis

The neurons were recorded with the internal solution (pipette solution) containing (in mM) 130 K-gluconate, 10 KCl, 5 EGTA, 10 HEPES, 1 MgCl_2_, 0.5 Na_3_GTP, 4 Mg-ATP, 10 Na-phoshocreatine pH 7.4 (adjusted with KOH) and external solution containing (in mM): 10 glucose, 125 NaCl, 25 NaHCO_3_, 1.25 NaH_2_PO4.2H_2_O, 2.5 KCl, 1.8 CaCl_2_, 1 MgCl_2_, pH 7.4 (300–310 mOsm), equilibrated with 95% O2 and 5% CO_2_.Whole cell recording are performed with multi-clamp 700b amplifier (Molecular Device), low-pass filtered at 1 kHz and the series resistance was typically <20 MΩ after >50% compensation. Cell-attached recordings for SNc and VTA neurons were performed before break-in with 0 mV holding. Current clamp recordings for spontaneous firing were performed with 0 pA current injection. To evoke action potential firing, SNc neurons were first held with 0 pA current holding followed by injection of increasingly positive current injection from –80 pA to 110 pA with 10 pA increment. VTA neurons were first held with 0 pA current holding followed by injection of increasingly positive current injection from –50 pA to 140 pA with 10 pA increment. Striatal neurons were first held with 0 pA current holding followed by injection of increasingly positive current injection from –100 pA to 280 pA with 20 pA increment. Whole-cell recordings were obtained from 3 control and 3 SJ1 cKO^DAT^ mice. A total of 79 WT THINs, 29 SNc and 21 VTA DANs were recorded from the control mice, 63 DALNs, 13 SNc and 21 VTA DANs were recorded from SJ1 cKO^DAT^ mice. Data was analyzed with Clampfit 11.4.2 and Prism.

Intrinsic electrophysiological properties were extracted as previously described^24,34,41,59^. A total of 14 parameters were quantified for striatal neurons, whereas 12 parameters were extracted for midbrain DANs, reflecting cell type–specific electrophysiological features.

For striatal neurons (WT THINs and DALNs), the extracted parameters included: maximal firing frequency, input resistance, resting membrane potential (RMP), action potential (AP) threshold, AP width at 50% relative to baseline and threshold, AP amplitude relative to baseline and threshold, after-hyperpolarization (AHP) amplitude, spontaneous activity, plateau potential duration, low-threshold spike (LTS) component, hyperpolarization-activated current (Ih ratio), and membrane capacitance.

For midbrain DANs, parameters included: maximal firing frequency, input resistance, AP threshold, AP width at 50% relative to baseline and threshold, AP amplitude relative to baseline and threshold, AHP amplitude, cell-attached firing frequency, break-in firing frequency, Ih ratio, and membrane capacitance.

For population-level analysis of striatal neurons, extracted intrinsic electrophysiological parameters were compiled into feature matrices and subjected to dimensionality reduction and clustering. Analyses were first performed on wild-type (WT) THINs to define baseline electrophysiological subtypes, followed by analysis of WT THINs–DALNs combined datasets using the same analytical framework to enable direct comparative mapping between conditions. This framework enabled unbiased identification of electrophysiologically distinct neuronal subtypes and quantitative assessment of genotype-dependent shifts in intrinsic neuronal properties.

Principal component analysis (PCA) was performed using singular value decomposition implemented in the pcaMethods package in R, with mean centering and unit variance scaling. Principal component scores were extracted, and components capturing the majority of variance were used for downstream clustering. Unsupervised hierarchical clustering was performed using Euclidean distance and Ward’s minimum variance method (ward.D2). The optimal number of clusters was determined by silhouette analysis using the cluster package.

For visualization, three electrophysiological parameters with the greatest discriminatory power between clusters were selected to generate three-dimensional feature-space representations. These were constructed in three configurations: WT clusters alone, DALNs projected onto WT-defined clusters, and combined clustering of WT THINs and DALNs. Radar plots were generated following feature-wise normalization to a 0–1 range to compare averaged electrophysiological profiles across clusters and conditions.

All statistical analyses and data visualization were performed in R using standard packages, including pheatmap, FactoMineR, ggplot2, and related libraries.

### Laser capture microdissection (LCM)

Fresh mouse brains were collected by decapitation and post-fixed in 4% PFA for 2 h at 4°C, followed by overnight cryoprotection in 30% sucrose. Brains were then embedded as described above and cryosectioned at 10 μm thickness onto PEN membrane slides (11600288, Leica). Prior to LCM, tissue sections were dehydrated through a graded ethanol series (50%, 70%, 75%, 90%, 95%, and 100%) and air-dried completely. Neurons were identified microscopically and isolated by LCM, with captured tissue collected directly into 8-strip collection caps. Total RNA was extracted from the microdissected tissue using the Arcturus PicoPure FFPE RNA Isolation Kit (KIT0312I, Thermo Fisher Scientific) according to the manufacturer’s instructions.

### Reverse Transcription PCR (RT-PCR)

Complementary DNA (cDNA) was synthesized from the extracted RNA using PureNA cDNA Synthesis Kit (KR01-100) according to the manufacturer’s instructions. PCR amplification was performed using 2 × Rapid Taq Master Mix (P222, Vazyme). Following are the gene-specific primers: NeuN, forward CACCACTCTCTTGTCCGTTTGC, reverse GGCTGAGCATATCTGTAAGCTGC; TH, forward TCAGAGGAGCCCGAGGTC, reverse GGGCGCTGGATACGAGAG; AADC, forward TACACATCTGATCAGGCGCA, reverse AAGGGCAGAAGCTCTCATGG; DAT, forward GCAGCTGGCACATCTATCCT, reverse ACCACGACCACATACAGAAGG; vMAT2, forward ATTGGCTTTCCTTGGCTCAT, reverse GGTACGGCTGGACATTATTCTG; ANXA1, forward CCTCATCTTCGCAGAGTGTTTC, reverse TCGGCAAAGAAAGCTGGAGT; ALDH1A1, forward GGAATACCGTGGTTGTCAAGCC, reverse CCAGGGACAATGTTTACCACGC; Gad2, forward CTGATTAAATGTGACGAGAGAGGG, reverse TCAGCTACAGCCAAGAGAGGA. The PCR products were separated by agarose gel electrophoresis and visualized using a nucleic acid staining dye under UV illumination. The presence of amplicons of the expected size was used to confirm successful amplification of the target genes.

## Statistical Analysis

All statistical analysis was performed using GraphPad Prism (Ver 10.6.1, RRID: SCR_002798) otherwise stated. All graphs were plotted on GraphPad Prism otherwise stated. Data are presented as mean ± SEM unless otherwise stated. Comparisons between two independent groups were conducted using two-tailed Welch’s unpaired t-tests. When multiple comparisons were performed, *p*-values were adjusted using the Holm-Šídák or Šídák’s correction. For the western blot analysis, a paired t-test was used because each mutant sample was matched to a corresponding control sample based on experimental batch and/or age. For analyses involving more than two groups, one-way or two-way ANOVA with Tukey’s honestly significant difference (HSD) post hoc test was applied. For repeated measures data from the 60 min open field test, two-way ANOVA with Tukey’s HSD post hoc test was performed. For normally distributed datasets from electrophysiology experiments, significance analysis was performed using unpaired Student’s t-tests (two-tailed) or ANOVA, followed by Tukey’s post-hoc test. For non-normally distributed datasets, Mann-Whitney U tests or Kruskal-Wallis tests were used, with Dunn correction for multiple comparisons. Data with *p*-values <0.05, <0.01, <0.001, and <0.0001 are represented by asterisks *, **, ***, and ****.

## Resource Availability

The datasets generated and/or analyzed in this study are available from the corresponding author Mian Cao on request. All other data needed to evaluate the conclusions in the paper are present in the paper and/or the Supplementary Materials.

## Author Contributions

Z.L., Y.L., B.M., M.L., I.M. and M.C. conceptualized and designed the study. Z.L. and Y.L. contributed equally. A.C.R., B.M. and M.L. contributed equally. Z.L., Y.L., M.L. and C.L. performed sample preparation, histology experiments and microscopy imaging. Y.L. performed HPLC and western blotting with assistance from X.C.. M.L. performed stereological quantification. Z.L., Y.L., M.L., X.C. and Z.W. performed behavioural experiment and analysis. Z.L. B.M., M.L. and C.L. performed stereotaxic injections. H.H. and Z.D. performed electrophysiology experiments. A.C.R and I.M. performed histology experiments, data collection and analysis for Endophilin A mouse model. M.L. and X.C. performed mouse genotyping and maintained the mouse breeding colony. Z.L., and Y.L. prepared the final figures. Z.L., Y.L., I.M. and M.C. wrote the paper, other authors contributed to its revision and final version.

## Acknowledgements

We thank Prof Zhang Suchun (Duke-NUS) and Asst Prof Jun Nishiyama (Duke-NUS) for providing some reagents for the experiments. We thank Prof Pietro De Camili (Yale University), Prof Shirish Shenolikar (Duke University) and Prof Wang Hongyan (Duke-NUS) for reading the manuscript and providing comments. We thank Prof Pietro De Caimili, Dr Sarah Luo Xinwei (IMCB-A*STAR) and Prof Enrico Petretto (Duke-NUS) for kindly providing SJ1^RQ^-KI, TH-IRES-Cre and Ai9 mouse line, respectively. We thank Asst Prof Wong Peiyan (Duke-NUS) from animal behaviour core for training. We thank Electron Microscopy Unit (NUS) for EM preparation and imaging. Some illustrations were created using BioRender. Select portions of this manuscript were edited for clarity and grammar using the ChatGPT language model (OpenAI). Scientific content and interpretations were solely formed by the authors.

## Supplemental information

Document S1. Figures S1–S15 and Table S1

