## Supplementary Figure Legends and Table S1 for "Synaptic endocytic failure drives dopamine deficiency and protein transfer-mediated striatal dopamine-like neuron compensation"

**Supplementary Figure Legend**

**Supplementary Figure 1. Further characterization of DA system in SJ1 cKO^DAT^ mice**

(A) Generation of SJ1 conditional KO mice by crossing SJ1 flx/flx mice with DAT-IRES-Cre line to create DA-specific KO, referred to as SJ1 cKO^DAT^. Appropriate littermate control (SJ1 flx/flx or DAT-Cre;SJ1 flx/+) were used in all experiments.

(B) Double staining of TH and SJ1 to show SJ1 expression in hippocampus, indicating that other regions were not affected in SJ1 cKO^DAT^ mice. Scale bar: 20 µm.

(C) Representative images of DAT immunostaining in the striatum of SJ1 cKO^DAT^ mice at 3 weeks, 2 months, 6 months and 12 months age group. Scale bar: 50 µm.

(D), (E) Tiling of a single coronal striatum section labelled for DAT in 2 months old SJ1 cKO^DAT^ (D) and SJ1^RQ^-KI mice (E). Unlike SJ1^RQ^-KI that showed clusters in dorsal striatum only, DAT clusters were observed in most of the striatum region of SJ1 cKO^DAT^ mice, including dorsal striatum (DS), ventral striatum (VS) and olfactory tubercle (OT). Scale bar: 500 µm. Inset scale bar: 50 µm.

(F) Immunostaining in nucleus accumbens (NAc) core showing TH and DAT positive clusters in the striatum of SJ1 cKO^DAT^ mice Scale bar: 50 µm.

(G) Immunostaining in NAc medial shell reveals lack of DAT signal and intact TH fibers in both control and SJ1 cKO^DAT^ mice Scale bar: 50 µm.

(H), (I) TH and DAT immunostaining of the prefrontal cortex (PFC) (H) and basolateral amygdala (BLA) (I), both receiving DA projections from the VTA shows intact TH fibers and lack of DAT expression. No obvious differences in TH-positive fibers were observed between control and mutant mice. Scale bar: 50 µm.

**Supplementary Figure 2. Other DA and synaptic markers examined in the SJ1 cKO^DAT^** **striatum**

(A) Immunostaining of vMAT2 with SV2C showed colocalization for clusters in the striatum of 2 months SJ1 cKO^DAT^ mice. Scale bar: 20 µm.

(B) Immunostaining of DAT with Syt1 showed partial colocalization with DAT-positive clusters in the striatum of 2 months SJ1 cKO^DAT^ mice. Scale bar: 20 µm.

(C), (D) Immunostaining of DAT with postsynaptic markers, D2R (C) and D1R (D), showed no colocalization with DAT-positive clusters in the striatum of 2 months SJ1 cKO^DAT^ mice, suggesting there is no effect on postsynaptic neurons. Scale bar: 20 µm.

(E) Immunostaining of DAT with Darpp32 showed no colocalization with DAT-positive clusters in the striatum of 2 months SJ1 cKO^DAT^ mice. Scale bar: 20 µm.

(F) Quantification of colocalization using Pearson’s correlation coefficient reflects the visual relationships shown in the striatum of SJ1 cKO^DAT^ mice (A–E). Higher r value for SV2C/vMAT2 indicate strong linear correlation, while other markers with lower r values show moderate to weak relationship. Data are represented as mean ± SEM. Data was computed using 4–6 randomly selected ROIs in the striatum.

**Supplementary Figure 3. Glia cell markers and alpha-Synculein examined in the SJ1 cKO^DAT^** **striatum**

(A), (B) Immunostaining of DAT with GFAP (A) and Iba1 (B) showed similar glia cell pattern in both Ctrl and SJ1 cKO^DAT^ mice at 2 months old, suggesting there is no gliosis and inflammation in the striatum. Scale bar: 20 µm.

(C) GFAP and Iba1 immunoreactive area fraction in the striatum of Ctrl and SJ1 cKO^DAT^ mice at 2 months old reveals no significant differences. n=3 per genotype.

(D) Immunostaining of alpha-Synuclein showed partial colocalization with DAT-positive clusters in the striatum of 2 months SJ1 cKO^DAT^ mice. Scale bar: 20 µm.

(E), (F) Immunostaining of pSer129-alpha-Synuclein with DAT (E) and TH (F) showed no detection of misfolded alpha-Synuclein in DAT-positive clusters and DANs, respectively, at 2 months old. Scale bar: 20 µm.

(G) Pearson’s correlation coefficient reflects the visual relationships shown in (D) and (E), showing mild and weak relationship, respectively. Data was computed using 4–5 randomly selected ROIs in the striatum.

Data are represented as mean ± SEM. Statistical analysis: Welch’s unpaired t-test (C).

**Supplementary Figure 4. Endocytic markers and whorl-like membrane accumulation examined in the SJ1 cKO^DAT^** **striatum**

(A)-(D) Double staining of DAT with SJ1 (A), clathrin-light chain (CLC) (B), amphiphysin 1 (C) and endophilin 1 (D) showed no colocalization of these clathrin-related endocytic proteins with DAT-positive clusters in the striatum of 2 months old SJ1 cKO^DAT^ mice. Scale bar: 20 µm.

(E) Pearson’s correlation coefficient reflects the visual relationships shown in (A)-(D), reflecting low colocalization between the endocytic proteins and the clusters. Data are represented as mean ± SEM. Data was computed using 3 randomly selected ROIs in the striatum.

(F) Additional representative EM images to show the abnormal membrane accumulation which is correlated with DAT/TH-positive protein clusters in the striatum of 2 months old SJ1 cKO^DAT^. Abnormal multilayered whorl membrane (red arrows) surrounding the synaptic vesicles (SVs) and endosomes (endo). Mitochondria (mt) also labelled in the terminals. Scale bar: 200 nm and 500nm.

**Supplementary Figure 5. Distribution and cell type identity of DALNs in SJ1 cKO^DAT^ striatum**

(A) Tiling of a single coronal striatum section with TH immunostaining in 2 months old SJ1 cKO^DAT^ mice, Scale bar: 500 µm. Insets: TH-positive clusters and DALNs were both observed throughout the striatum, including dorsal striatum (DS), ventral striatum (VS) and olfactory tubercle (OT), Scale bar: 20 µm. (B) Representative images of TH immunostaining in the striatum of SJ1 cKO^DAT^ at 3 weeks, 2 months, 6 months and 12 months age group. TH-positive DALNs (white arrows) were labelled. Scale bar: 50 µm.

(C) Quantification of DALNs in the striatum of SJ1 cKO^DAT^ mice from 3 weeks to 12 months old. Number of DALNs were quantified on single coronal striatum sections of one hemisphere. Data are represented as mean ± SEM. n= 1-3.

(D)-(F) Immunostaining of TH with NPY (D), PV (E) and ChAT (F) in SJ1 cKO^DAT^ striatum showed that DALNs is a distinct group of neurons from other striatal interneurons. Scale bar: 20 µm.

(G), (H) Very few DALNs in the striatum of SJ1 cKO^DAT^ occasionally co-express Calretinin (G) and Darpp32 (H), suggesting heterogeneity within DALNs subpopulation. Scale bar: 20 µm.

**Supplementary Figure 6. A subpopulation of DALNs in SJ1 cKO^DAT^** **dorsal striatum expresses ALDH1A1 and ANXA1**

(A) Immunostaining of TH and ALDH1A1 in the midbrain of 2 months old mice shows enriched ALDH1A1 in the ventral tier of SNc DANs. Scale bar: 200 µm.

(B) Tiling of a single coronal striatum section immunostained for TH and ALDH1A1 of 2 months old Ctrl and SJ1 cKO^DAT^ mice. TH/ALDH1A1-positive DALNs were mainly distributed in DS, while VS DALNs are ALDH1A1 negative. Scale bar: 500 µm

(C) Immunostaining of TH and ANXA1 in the midbrain of 2 months old mice shows Anxa1 is enriched in the ventral tier of SNc DA neurons in both genotypes. Scale bar: 200 µm.

(D) Tiling of a single coronal striatum section immunostained for TH and ANXA1 of 2 months old Ctrl and SJ1 cKO^DAT^ mice. TH/ANXA1 double positive DALNs were mainly distributed in dorsolateral striatum. Scale bar: 500 µm

(E) Immunostaining shows ANXA1-positive DANs within a subpopulation of ALDH1A1-positive DANs in the midbrain of 2 months old mice. Scale bar: 200 µm.

(F) A subset of ALDH1A1-positive neurons is positive for ANXA1 in the dorsolateral striatum of SJ1 cKO^DAT^ mice (white arrows). Scale bar: 50 µm.

(G) Simplified illustrations of Calbindin distribution in the midbrain and striatum.

(H) Immunostaining of TH and Calbindin in the midbrain of 2 months old mice show some VTA DANs expressing Calbindin. Scale bar: 20 µm.

(I) Immunostaining of TH and Calbindin shows that Calbindin was not detected in VS DALNs of 2 months old SJ1 cKO^DAT^ mice. Scale bar: 50 µm.

**Supplementary Figure 7. SJ1 cKO^TH^ mice phenocopies SJ1 cKO^DAT^ pathological characteristics**

(A) Double staining of TH and SJ1 showed specific loss of SJ1 in SJ1 cKO^TH^ DANs. SJ1 expression in hippocampus shows that other regions were not affected. Scale bar: 20 µm.

(B), (C) Double staining of ALDH1A1 and TH shows ALDH1A1 is detected in both dystrophic DA terminals (asterik) and DALNs (white arrows) in SJ1 cKO^TH^ dorsal (B) but not ventral (C) striatum. Scale bar: 50 µm.

(D) Representative images of tdTomato expression shown in the midbrain TH-positive DANs of 2 months old Ai9;TH-Cre (Ctrl) and Ai9;SJ1 cKO^TH^ mice. Scale bar: 100 µm.

(E) Representative images of tdTomato expression shown in the midbrain TH-positive DANs in the midbrain of 2 months old Ai9;DAT-Cre (Ctrl) and Ai9;SJ1cKO^DAT^ mice. Scale bar: 100 µm.

**Supplementary Figure 8. Electrophysiological properties of DALNs and DANs**

(A) Bar charts showing the proportion of WT THINs and DALNs within each of the four clusters identified by PCA-based clustering of the combined dataset. Data from WT THINs and DALNs were pooled prior to dimensionality reduction and clustering.

(B) Bar charts comparing ten representative electrophysiological parameters among 3 WT THIN clusters and DALNs, including action potential (AP) width at 50% relative to baseline and threshold, AP amplitude relative to baseline and threshold, AP threshold, maximal firing frequency, after-hyperpolarization (AHP) amplitude, hyperpolarization-activated current (Ih ratio), membrane capacitance and low-threshold spike (LTS).

(C) Representative images of a patch-clamped tdTomato-positive DANs in the SNc. Upper: differential interference contrast (DIC) image showing the patched neuron. Lower: epifluorescence image showing tdTomato (RFP) expression in the same cell. Scale bar: 20 µm.

(D) Spontaneous firing of Ctrl and SJ1 cKO^DAT^ SNc DANs recorded in cell-attached (on-cell) configuration.

(E) Intrinsic firing of Ctrl and SJ1 cKO^DAT^ SNc DANs recorded in whole-cell current-clamp mode (I = 0 pA).

(F) Representative evoked firing traces of Ctrl and SJ1 cKO^DAT^ SNc DANs.

(G) Representative traces of the first action potential elicited in Ctrl and SJ1 cKO^DAT^ SNc DANs.

(H) Bar charts comparing Ctrl and SJ1 cKO^DAT^ SNc DANs. Compared to Ctrl, cKO neurons exhibited significantly reduced AP width at 50% relative to threshold and baseline, as well as a lower AP threshold. Conversely, AP amplitude relative to both threshold and baseline was significantly increased. Each data point represents one cell, n = 3 mice per group.

(I) Representative images of a patch-clamped tdTomato-positive DAN in the VTA. Upper: differential interference contrast (DIC) image showing the patched neuron. Lower: epifluorescence image showing tdTomato (RFP) expression in the same cell. Scale bar: 20 µm.

(J) Spontaneous firing of Ctrl and SJ1 cKO^DAT^ VTA DANs recorded in cell-attached (on-cell) configuration.

(K) Intrinsic firing of Ctrl and SJ1 cKO^DAT^ VTA DANs recorded in whole-cell current-clamp mode (I = 0 pA).

(L) Representative evoked firing traces in Ctrl and SJ1 cKO^DAT^ VTA DANs.

(M) Representative traces of the first action potential elicited in Ctrl and SJ1 cKO^DAT^ VTA DANs.

(N) Bar charts comparing Ctrl and SJ1cKO^DAT^ VTA DA neurons. Compared to Ctrl, SJ1 cKO^DAT^ VTA DANs showed a significant increase in input resistance, with no other intrinsic properties significantly altered. Each data point represents one cell, n = 3 mice per group.

Data are presented as mean ± SEM. Statistical analysis: Kruskal–Wallis test followed by Dunn’s multiple comparison test (C), two-tailed Mann–Whitney U test (H, N); **p < 0.01, ***p < 0.001, ****p < 0.001.

**Supplementary Figure 9. RNAscope-IF validation of TH protein and dopaminergic transcripts across brain regions and genotypes**

(A) Representative RNAscope-IF images of TH protein and *Th* mRNA in the locus coeruleus (LC), olfactory bulb (OB), cortex and cerebellum of SJ1 cKO^DAT^ mice. TH-positive neurons in these regions also expressed *Th* mRNA. Scale bar: 20 µm.

(B) Representative RNAscope-IF images of TH protein and *Th* mRNA in the midbrain and striatum of SJ1 cKO^TH^ mice. Midbrain DANs were positive for both TH protein and *Th* mRNA. In the striatum, DALNs were TH-positive but *Th* mRNA-negative (white arrows), whereas WT THINs were TH-negative but *Th* mRNA-positive (yellow arrows). Scale bar: 20 µm.

(C, D) Representative RNAscope-IF images of TH protein and *Th* mRNA in the midbrain and striatum of TH-Cre (C) and DAT-Cre (D) control mice. Midbrain DANs expressed both TH protein and *Th* mRNA, whereas striatal WT THINs were TH-negative but *Th* mRNA-positive (yellow arrows). Scale bars: 20 µm.

(E) Representative RNAscope-IF images of TH protein together with *Th* and *Ddc* mRNAs in the midbrain and striatum of WT mice. Midbrain DANs expressed TH protein, *Th* mRNA and *Ddc* mRNA. Striatal WT THINs expressed *Th* mRNA but lacked detectable TH protein and *Ddc* mRNA (yellow arrows). Scale bar: 20 µm.

**Supplementary Figure 10. RNAscope-IF analysis of additional dopaminergic markers and assay controls in SJ1 cKO^DAT^ mice**

(A) Representative RNAscope-IF images of TH and ALDH1A1 proteins together with *Aldh1a1* mRNA in the midbrain and striatum. ALDH1A1-positive midbrain DANs expressed the corresponding *Aldh1a1* mRNA, whereas ALDH1A1-positive DALNs lacked detectable *Aldh1a1* mRNA (white arrows). Scale bar: 20 µm.

(B) Representative RNAscope-IF images of TH and ANXA1 proteins together with *Anxa1* mRNA. ANXA1-positive midbrain DANs expressed *Anxa1* mRNA. In the striatum, both ANXA1-negative (white arrows) and ANXA1-positive (yellow arrows) DALNs lacked detectable *Anxa1* mRNA. Scale bar: 20 µm.

(C) Representative RNAscope-IF images of TH and DAT proteins together with *Slc6a3* mRNA. Midbrain DANs expressed both DAT protein and *Slc6a3* mRNA, whereas TH-positive DALNs lacked detectable DAT protein and *Slc6a3* mRNA (white arrows). Yellow asterisks indicate DAT-positive clusters in the striatum. Scale bar: 20 µm.

(D, E) Representative RNAscope-IF images of TH protein together with the housekeeping transcripts *Polr2a* (D) and *Ppib* (E). Both transcripts were detected in midbrain DANs, striatal DALNs (white arrows) and surrounding cells. Scale bars: 20 µm.

(F) Representative images of the negative-control probe *DapB-C1* in the midbrain and striatum. No detectable signal was observed in either region. Scale bar: 20 µm.

**Supplementary Figure 11. RNAscope-IF analysis of Ai9-tdTomato reporter expression in DALNs and WT THINs**

(A) Representative RNAscope-IF images of TH and tdTomato proteins together with *tdT* mRNA in the midbrain and striatum of Ai9;SJ1 cKO^TH^ mice. Midbrain DANs were positive for TH protein, tdTomato protein and *tdT* mRNA. In the striatum, DALNs were positive for TH and tdTomato proteins but lacked detectable *tdT* mRNA (white arrows), whereas WT THINs were positive for tdTomato protein and *tdT* mRNA but lacked detectable TH protein (pink arrows). Scale bar: 20 µm.

(B) Representative RNAscope-IF images of TH protein together with *Th* and *tdT* mRNAs in Ai9;SJ1 cKO^TH^ mice. Midbrain DANs were positive for TH protein and both transcripts. Striatal DALNs were TH-positive but lacked detectable *Th* and *tdT* mRNAs (white arrows), whereas WT THINs expressed both transcripts but lacked detectable TH protein (pink arrows). Scale bar: 20 µm.

(C) Representative RNAscope-IF images of TH and tdTomato proteins together with *tdT* mRNA in Ai9;TH-Cre control mice. In the striatum, WT THINs were positive for tdTomato protein and *tdT* mRNA but lacked detectable TH protein (pink arrows). Scale bar: 20 µm.

(D) Representative RNAscope images of *Th* and *tdT* mRNAs in the striatum of Ai9;TH-Cre control mice. WT THINs expressed both transcripts (pink arrows). Scale bar: 20 µm.

**Supplementary Figure 12. Evaluation of cell-type specificity and cargo-transfer selectivity using additional viral constructs**

(A) Schematic showing experiment procedure and timeline. AAV2-mTH-EGFP was injected into the ventral midbrain of B6, SJ1 cKO^TH^ and SJ1 cKO^DAT^ mice at 2-3 months old. The animal brains were collected 2 weeks post injection for IHC.

(B) Representative images of TH immunostaining with EGFP expression in the injected midbrain DANs of both SJ1 cKO^TH^ and SJ1 cKO^DAT^ mice. Scale bar: 100 µm.

(C) Representative images of TH immunostaining and EGFP expression in the striatum. No EGFP expressing neurons observed in B6 striatum. Striatum of injected SJ1 cKO^TH^ and SJ1 cKO^DAT^ mice show EGFP signal specifically in DALNs. White arrows: DALNs double positive for EGFP and anti-TH staining. Scale bar: 20 µm.

(D,E) Percentage of EGFP expressing neurons positive for anti-TH staining in both SJ1 cKO^TH^ and SJ1 cKO^DAT^. The number of neurons was quantified on a single stitched coronal striatal section for both EGFP neurons and TH/EGFP-positive neurons. Data are represented as mean ± SEM. n=3 mice per genotype.

(F) Schematic showing experiment procedure and timeline. AAV5-hSyn-mCherry was injected unilaterally in the midbrain of B6, SJ1 cKO^TH^ and SJ1 cKO^DAT^ mice at 2-3 months old. The animal brains were collected 2 weeks post injection for IHC.

(G) Representative images of TH immunostaining and mCherry expression in the injected midbrain DANs of both SJ1 cKO^TH^ and SJ1 cKO^DAT^ mice. Scale bar: 100 µm.

(H) Representative images of TH immunostaining and mCherry expression in the striatum of injected mice. No mCherry labeled neurons in B6 striatum. Striatum of injected SJ1 cKO^TH^ and SJ1 cKO^DAT^ show mCherry specifically expressed in DALNs. White arrows: DALNs double positive for mCherry and anti-TH staining. Scale bar: 20 µm.

(I,J) Percentage of mCherry expressing neurons positive for anti-TH staining in both SJ1 cKO^TH^ and SJ1 cKO^DAT^. The number of neurons was quantified on a single stitched coronal striatal section for both mCherry and TH/mCherry-positive neurons. Data are represented as mean ± SEM. n=3 mice per genotype.

(K) Schematic showing the experimental procedure and timeline. AAV2-mTH-EGFP was injected into the ventral midbrain of B6, Ai9;TH-Cre and Ai9;SJ1 cKO^TH^ mice at 2–3 months of age. Brains were collected 2 weeks after injection for IHC.

(L) Representative images of TH, tdTomato and EGFP signals in the midbrain and striatum. WT THINs were tdTomato-positive but TH-negative and lacked detectable EGFP signal (pink arrows). In contrast, DALNs were positive for TH, tdTomato and EGFP (yellow arrows). Scale bars: 20 µm.

(M) Schematic showing the experimental procedure and timeline. AAV5-hSyn-mCherry was injected into the striatum of B6 mice, and brains were collected 2 weeks later for IHC.

(N) Representative striatal images showing pan-neuronal expression of mCherry following AAV5-hSyn-mCherry injection. A subset of mCherry-positive neurons also expressed DARPP32. Scale bar: 20 µm.

(O) Experimental design for assessing the transfer of membrane-associated and nuclear-localized cargo. AAV5-DIO-ChR2-EYFP or AAV2/9-DIO-H2B-EGFP was injected into the ventral midbrain, and brains were collected 2 weeks later.

(P, Q) Representative midbrain and striatal images following expression of ChR2-EYFP (P) or H2B-EGFP (Q). Both constructs selectively labelled midbrain DANs, whereas no detectable GFP signal was observed in striatal DALNs. Scale bars: 20 µm.

**Supplementary Figure 13. SJ1 cKO^DAT^ mice show resistance to 6-OHDA-induced dopaminergic lesions**

(A) Experimental design for the 6-OHDA-induced dopaminergic lesion. 6-OHDA (3 µg in 1 µL) was unilaterally injected into the ventral midbrain of control and SJ1 cKO^DAT^ mice. Rotational behaviour was assessed at 0, 2 and 4 weeks, and brains were collected at 2 weeks for immunohistochemical analysis.

(B, C) Representative TH immunostaining of midbrain DANs soma (B) and DAT immunostaining of striatal DA terminals (C) 2 weeks after lesion. Scale bars: 200 µm (B) and 20 µm (C).

(D, E) Quantification of surviving midbrain TH-positive DAN soma (D) and striatal DAT-positive terminals (E). SJ1 cKO^DAT^ mice showed significantly greater survival than controls. Each point represents one mouse. n = 5 mice per genotype.

(F) Postoperative survival curves showing higher survival in SJ1 cKO^DAT^ mice than in controls following 6-OHDA injection. Postoperative survival was analysed in 51 mice, including 11 SJ1 cKO^DAT^ mice and 40 controls. Survival curves were compared using the log-rank (Mant–Cox) test.

(G) Ipsilateral rotation following 6-OHDA injection. Both genotypes showed balanced behaviour at baseline. Control mice developed sustained ipsilateral rotation at 2 and 4 weeks, whereas SJ1 cKO^DAT^ mice maintained behavioural balance. Each point represents one mouse. n = 4 mice per genotype.

Data are presented as mean ± SEM. Statistical analysis: Mann–Whitney test (D,E), Mann–Whitney test, followed by Holm–Šídák correction for multiple comparisons (G); **p < 0.01, ***p < 0.001, ****p < 0.001.

**Supplementary Figure 14. RNAscope analysis of Ai9;DAT-Cre;SJ1^RQ^-KI mice, other dopaminergic markers in Endophilin A1/2/3 KO/Ht/KO striatum and controls for SJ1 Acute KO model**

(A) Representative RNAscope-IF images of TH and tdTomato proteins together with *tdT* mRNA in the midbrain and striatum of Ai9;DAT-Cre;SJ1^RQ^-KI mice. Midbrain DANs expressed TH protein, tdTomato protein and *tdT* mRNA, whereas striatal DALNs were positive for TH and tdTomato proteins but lacked detectable *tdT* mRNA (white arrows). Scale bar: 20 µm.

(B) Tiling of a single coronal striatum section with TH immunostaining in Endophilin A1/2/3 KO/Ht/KO mice, Scale bar: 500 µm. Insets: TH-positive clusters and DALNs were both observed throughout the striatum, including dorsal striatum (DS) and ventral striatum (VS), Scale bar: 20 µm.

(C), (D), (E) Representative striatal immunostaining from Endophilin A1/2/3 KO/Ht/KO mice for SV2C (C), ALDH1A1 (D) and ANXA1 (E) together with TH. White arrows indicate marker-positive DALNs where applicable. Scale bars: 20 µm.

(F) Quantification of DALN numbers across genotypes. There is no DALNs detected in WT and AuxKO striatum. AuxKO/SJ1^RQ^-KI mice exhibited significantly more DALNs than SJ1^RQ^-KI mice, whereas SJ1 cKO^DAT^ and Endophilin A1/2/3 KO/Ht/KO mice showed a further increase compared with Aux-KO/SJ1^RQ^-KI mice. Each point represents one mouse. n = 4 mice per genotype.

(G) Schematic summary of the abundance and striatal distribution of DALNs and their ALDH1A1- and ANXA1-positive subtypes in SJ1^RQ^-KI, AuxKO/SJ1^RQ^-KI, Endophilin A1/2/3 KO/Ht/KO and SJ1 cKO^DAT^ mice.

(H), (I) Representative images of adult B6 mice (2 months old) (H) and aged B6 mice (12 months old) (I) unilaterally injected with AAV9-TH-Cre virus show no loss of DANs in the midbrain or DA fibers in the striatum, validating the non-toxicity of AAV9-TH-Cre used. Scale bar: 200 µm.

Data are presented as mean ± SEM. Statistical analysis: one-way ANOVA followed by Tukey’s test (F). *p < 0.05, **p < 0.01, ***p < 0.001 and ****p < 0.0001.

**Supplementary Figure 15. Acute DANs lesions induce striatal TH-positive neurons distinct from DALNs**

(A) Schematic showing the midbrain of 2-3 months old B6 mice unilaterally injected with 6-OHDA and collected 2 weeks post injection for IHC.

(B) Representative TH staining on coronal sections showing the extent of 6-OHDA lesion in the striatum and midbrain DANs of B6 mice. Scale bar: 500 µm.

(C) Schematic showing the midbrain of 2-3 months old DAT-Cre mice unilaterally injected with AAV5-FLEX-Caspase3 (Cre-dependent) and collected 2 weeks post injection for IHC.

(D) Representative TH staining on coronal sections showing the extent of Casp3-mediated lesion in the striatum and midbrain DANs of DAT-Cre mice. Scale bar: 500 µm.

(E) Representative images of TH-positive neurons (white arrows) observed in the striatum of SJ1 cKO^DAT^ and lesioned hemisphere of 6-OHDA and Casp3 injected mice, compared to control/unlesioned striatum. Exposure time and threshold were set the same for anti-TH imaging in all three conditions. Scale bar: 20 µm.

(F) The TH protein expression level in WT THINs observed in 6-OHDA and Casp3 lesioned models is significantly lower than DALNs observed in SJ1 cKO^DAT^ striatum. n=3 mice per genotype.

(G) Unlike DALNs in SJ1 cKO^DAT^ mice, AADC is undetectable in either 6-OHDA or Casp3 lesioned striatum. Scale bar: 20 µm.

(H) Unlike DALNs in SJ1 cKO^DAT^ mice, ALDH1A1 is undetectable in either 6-OHDA or Casp3 lesioned striatum. Scale bar: 20 µm.

(I) Representative striatum images of Ai9;DAT-Cre mice unilaterally injected with Casp3 (Cre-mediated) in the midbrain. TH-positive neurons appearing in the striatum of lesioned model (white arrows) did not show DAT-cre dependent tdTomato expression. Scale bar: 200 µm.

(J) Table for comparison of striatal TH-positive neurons observed in WT, genetic SJ1 DA cKO and acute DA lesion mouse models.

Data are presented as mean ± SEM. Statistical analysis: one-way ANOVA followed by Tukey’s test (F). ****p < 0.0001.

**Supplementary Table 1. Electrophysiological properties of WT THINs and DALNs**

| Parameter | WT Cluster 1 | WT Cluster 2 | WT Cluster 3 | DALNs |
| --- | --- | --- | --- | --- |
| AHP amplitude | 10.5 ± 0.5 (2.2 to 17.0)  n=39 | 8.2 ± 0.5 (1.6 to 12.7)  n=28 | 7.4 ± 1.2 (2.2 to 14.1)  n=12 | 12.2±0.4 (5.7 to 17.0)  n=63 |
| AP amplitude relative to baseline | 78.8 ± 1.5 (58.2 to 99.1)  n=39 | 89.4 ± 1.7 (69.8 to 103.4)  n=28 | 59 ± 4 (41 to 81)  n=12 | 93.8±1.7 (54.9 to 112.6)  n=63 |
| AP amplitude relative to threshold | 68.2 ± 1.4 (50.7 to 86.0)  n=39 | 81.1 ± 1.5 (63.6 to 94.6)  n=28 | 52 ± 3 (36 to 69)  n=12 | 81.5±1.6 (48.5 to 101.5)  n=63 |
| AP threshold | -39.8 ± 0.9 (-51.8 to -22.8)  n=39 | -43.5 ± 1.0 (-54.8 to -33.5)  n=28 | -36.2 ± 0.9 (-41.5 to -31.1)  n=12 | -41.4±0.6 (-50.7 to -26.5)  n=63 |
| AP width 50% relative to baseline | 1.36 ± 0.11 (0.69 to 3.18)  n=39 | 2.72 ± 0.11 (0.97 to 4.00)  n=28 | 3.2 ± 0.5 (1.2 to 5.0)  n=12 | 2.07 ± 0.05 (1.51 to 4.19)  n=63 |
| AP width 50% relative to threshold | 1.23 ± 0.10 (0.59 to 2.82)  n=39 | 2.51 ± 0.11 (0.90 to 3.77)  n=28 | 3.5 ± 0.7 (1.1 to 9.8)  n=12 | 1.88 ± 0.05 (1.34 to 3.81)  n=63 |
| Ih ratio | 1.034 ± 0.006 (1.000 to 1.210)  n=39 | 1.046 ± 0.006 (1.000 to 1.100)  n=28 | 1.047 ± 0.018 (1.000 to 1.240)  n=12 | 1.020 ± 0.003 (1.000 to 1.100)  n=63 |
| Input resistance | 187 ± 15 (70 to 440)  n=39 | 360 ± 18 (120 to 590)  n=28 | 620 ± 50 (450 to 1090)  n=12 | 112 ± 7 (30 to 250)  n=63 |
| LTS component | 0 of 39 | 2 of 28 | 10 of 12 | 0 of 63 |
| Max AP frequency | 42 ± 4 (9 to 97)  n=39 | 16.0 ± 1.0 (6.0 to 25.0)  n=28 | 12 ± 3 (1 to 27)  n=12 | 21.0 ± 1.1 (1.0 to 39.0)  n=63 |
| Membrane Capacitance | 17.8 ± 1.1 (7.8 to 42.0)  n=39 | 18.5 ± 1.4 (8.1 to 42.0)  n=28 | 11.5 ± 1.1 (7.4 to 19.2)  n=12 | 19.4 ± 1.0 (7.4 to 53.3)  n=63 |
| Plateau potential duration | 6 of 39 | 18 of 28 | 5 of 12 | 0 of 63 |
| RMP | -65.3 ± 1.4 (-81.7 to -46.0)  n=39 | -62.1 ± 1.5 (-74.8 to -48.8)  n=28 | -48 ± 3 (-61 to -27)  n=12 | -71.5 ± 0.9 (-84.5 to -52.3)  n=63 |
| Spontaneous Activity | 0 of 39 | 4 of 28 | 8 of 12 | 0 of 63 |
