## Supplementary Figure 1-15 for "Synaptic endocytic failure drives dopamine deficiency and protein transfer-mediated striatal dopamine-like neuron compensation"

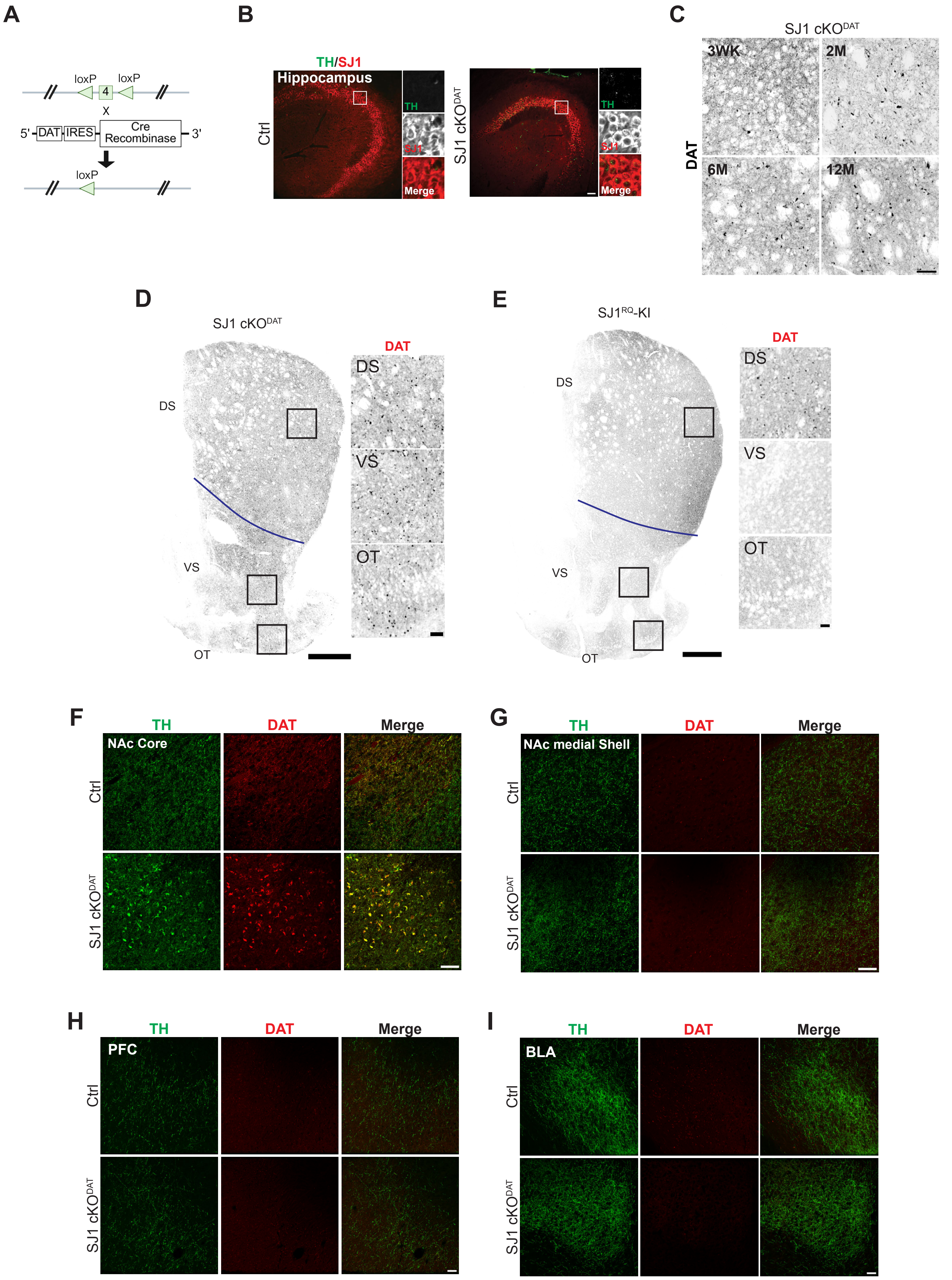


### Supplementary Figure 1

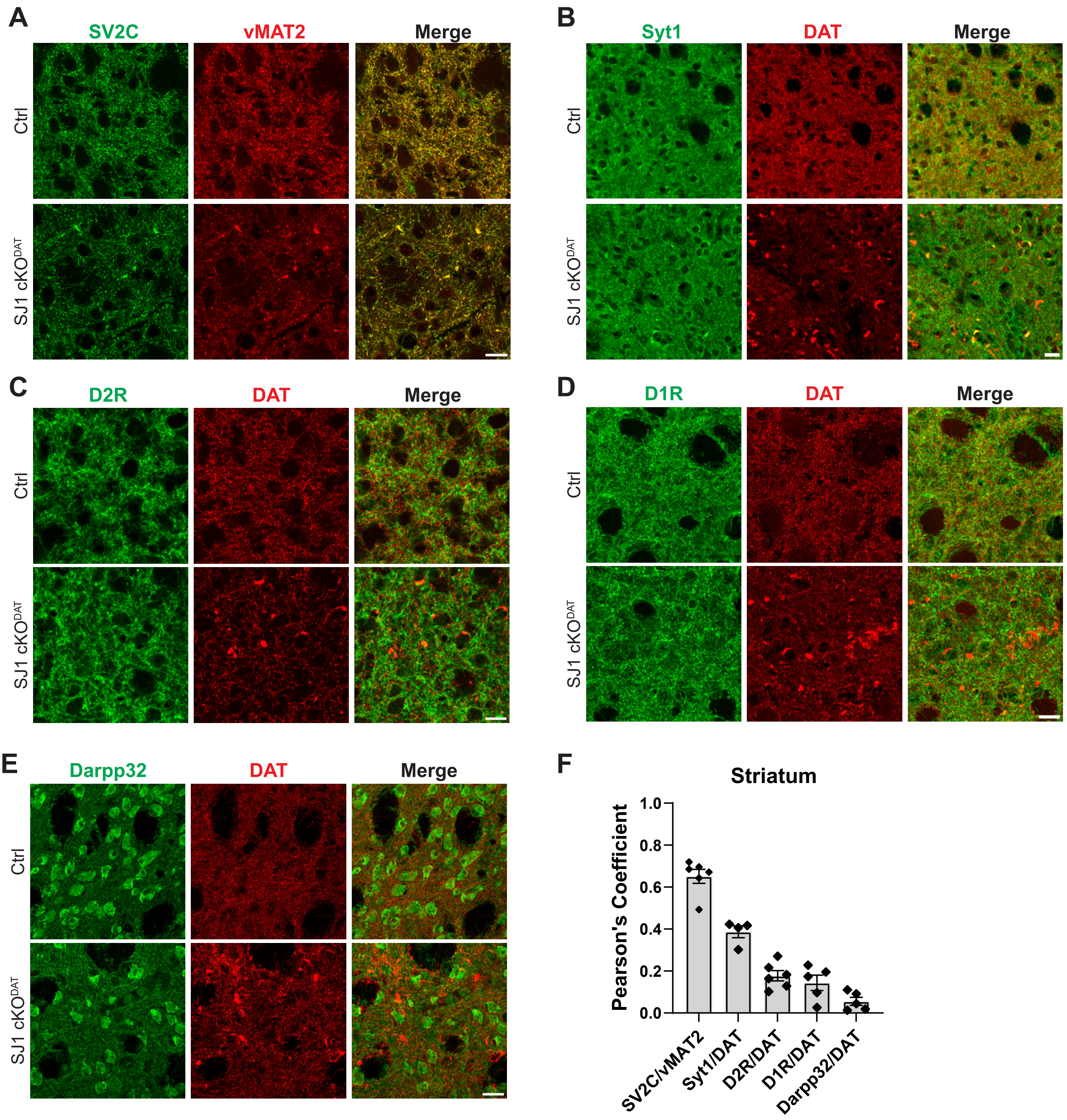


**Supplementary Figure 2**

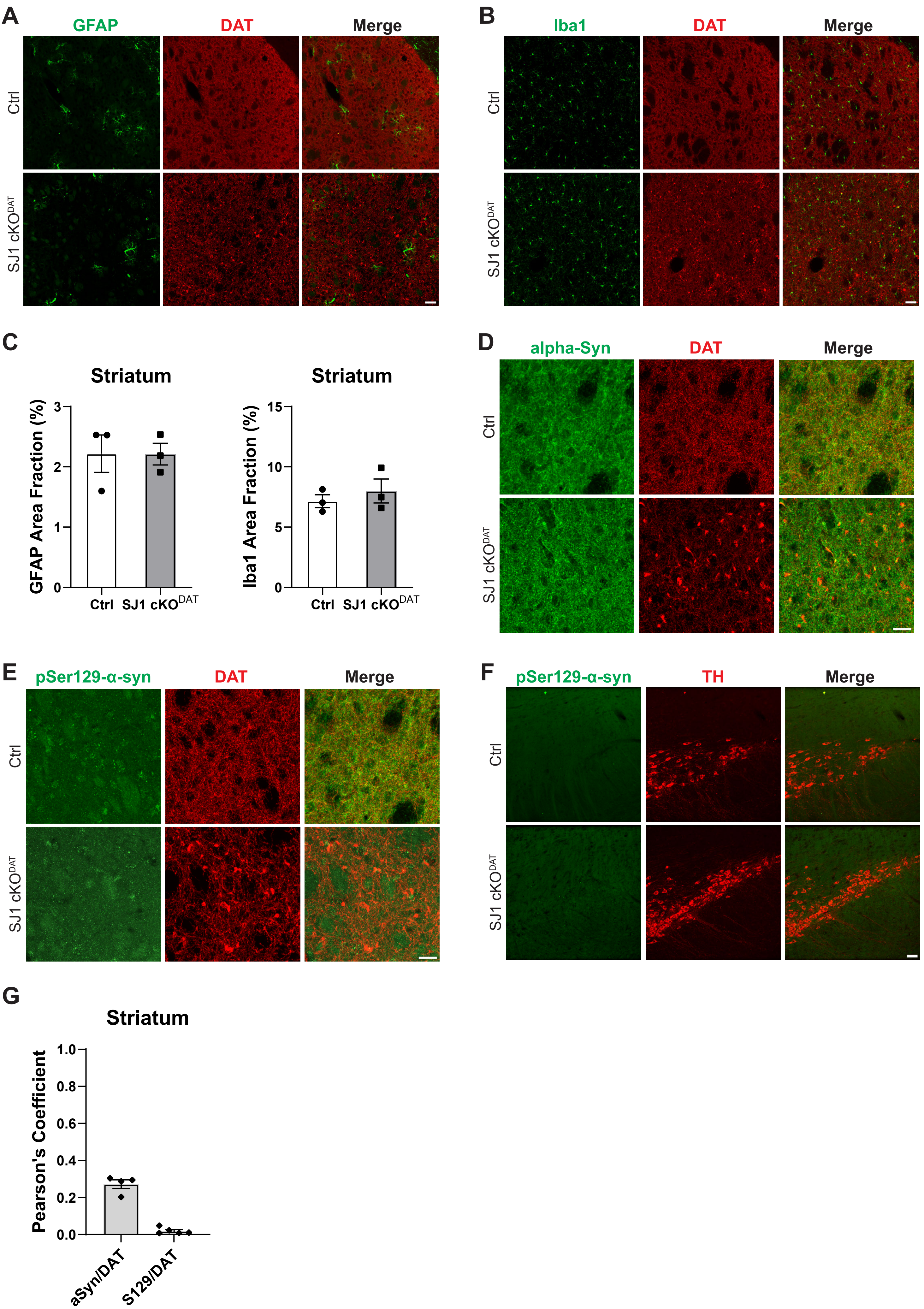


Supplementary Figure 3

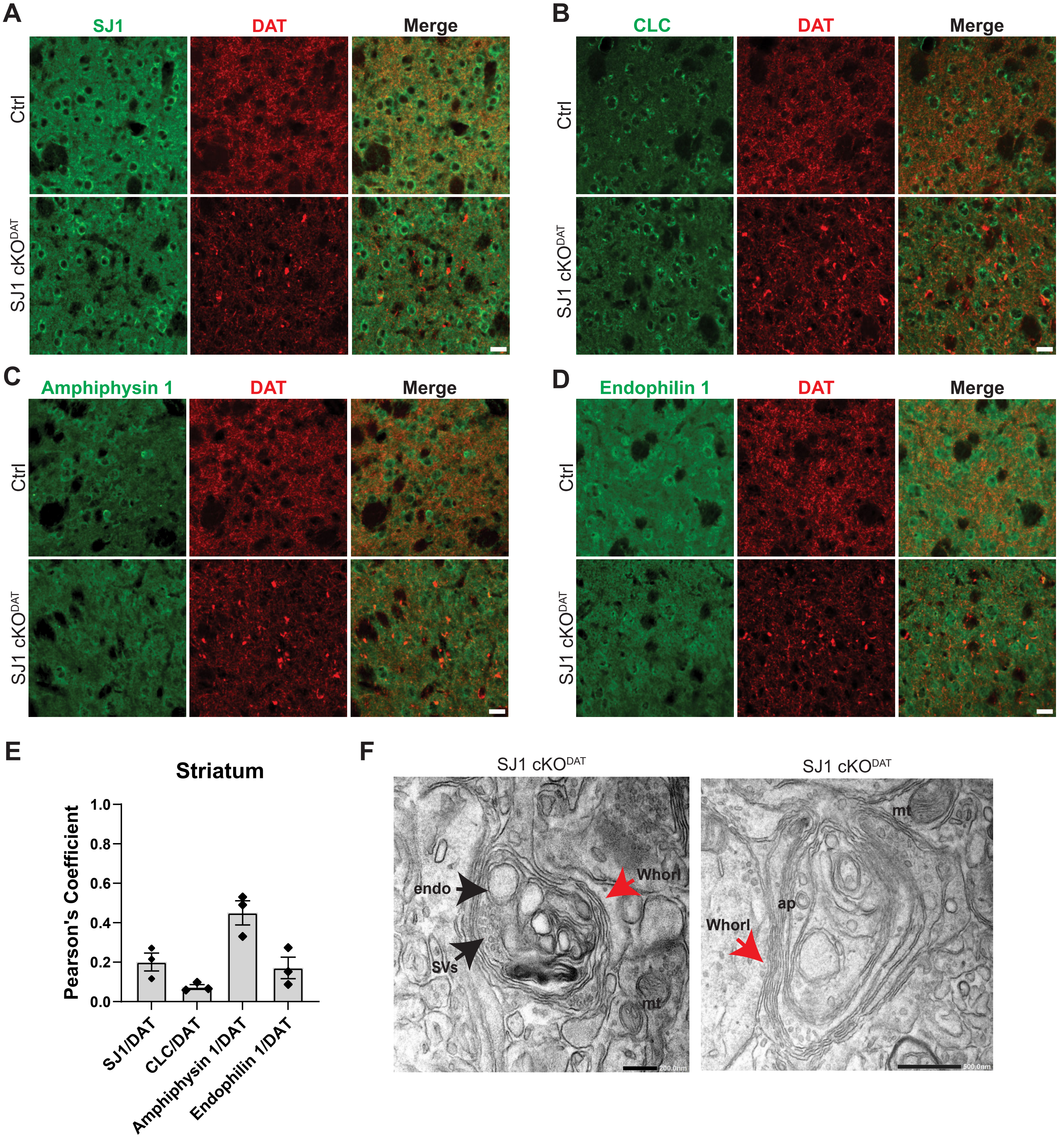


Supplementary Figure 4

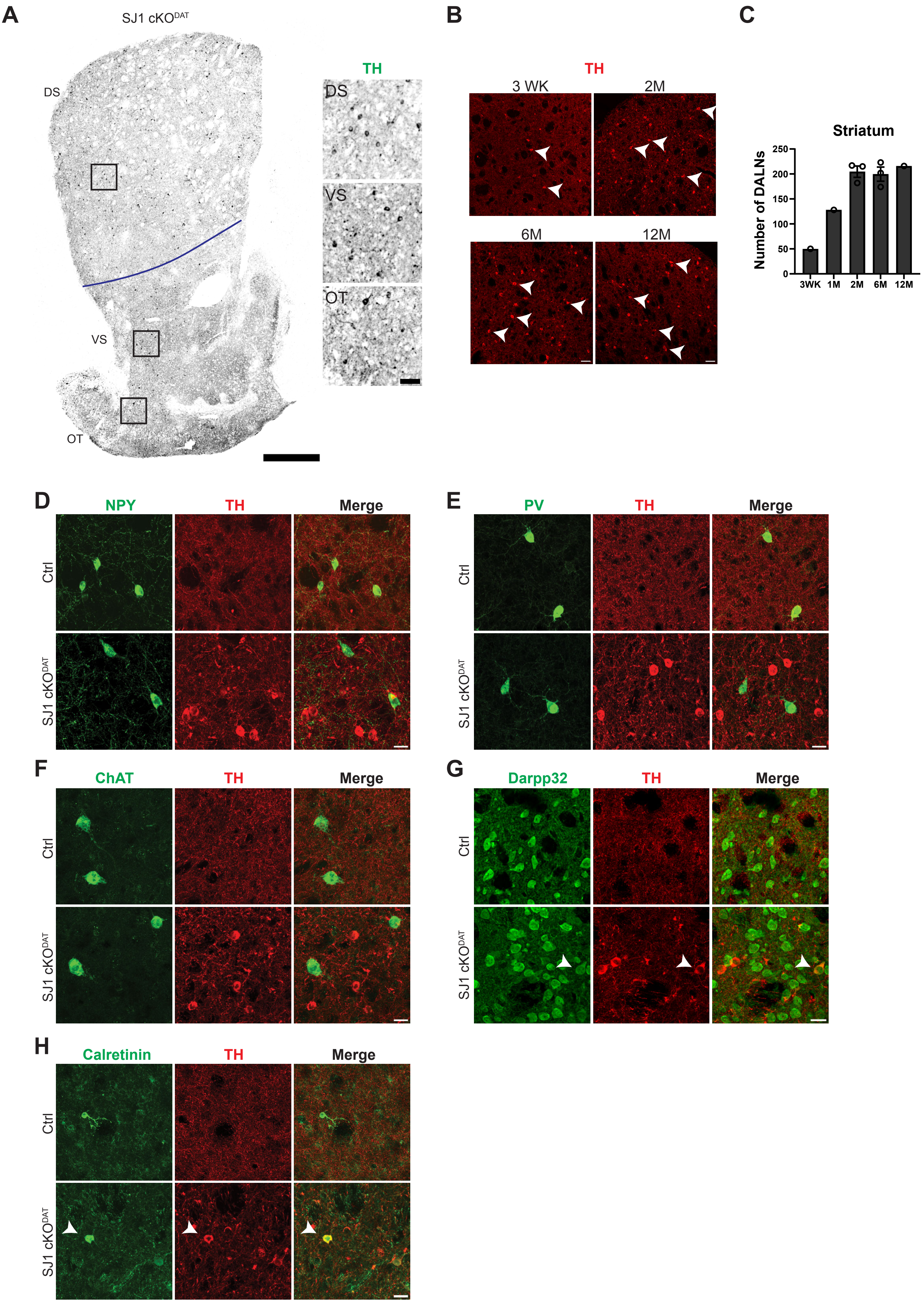


**Supplementary Figure 5**

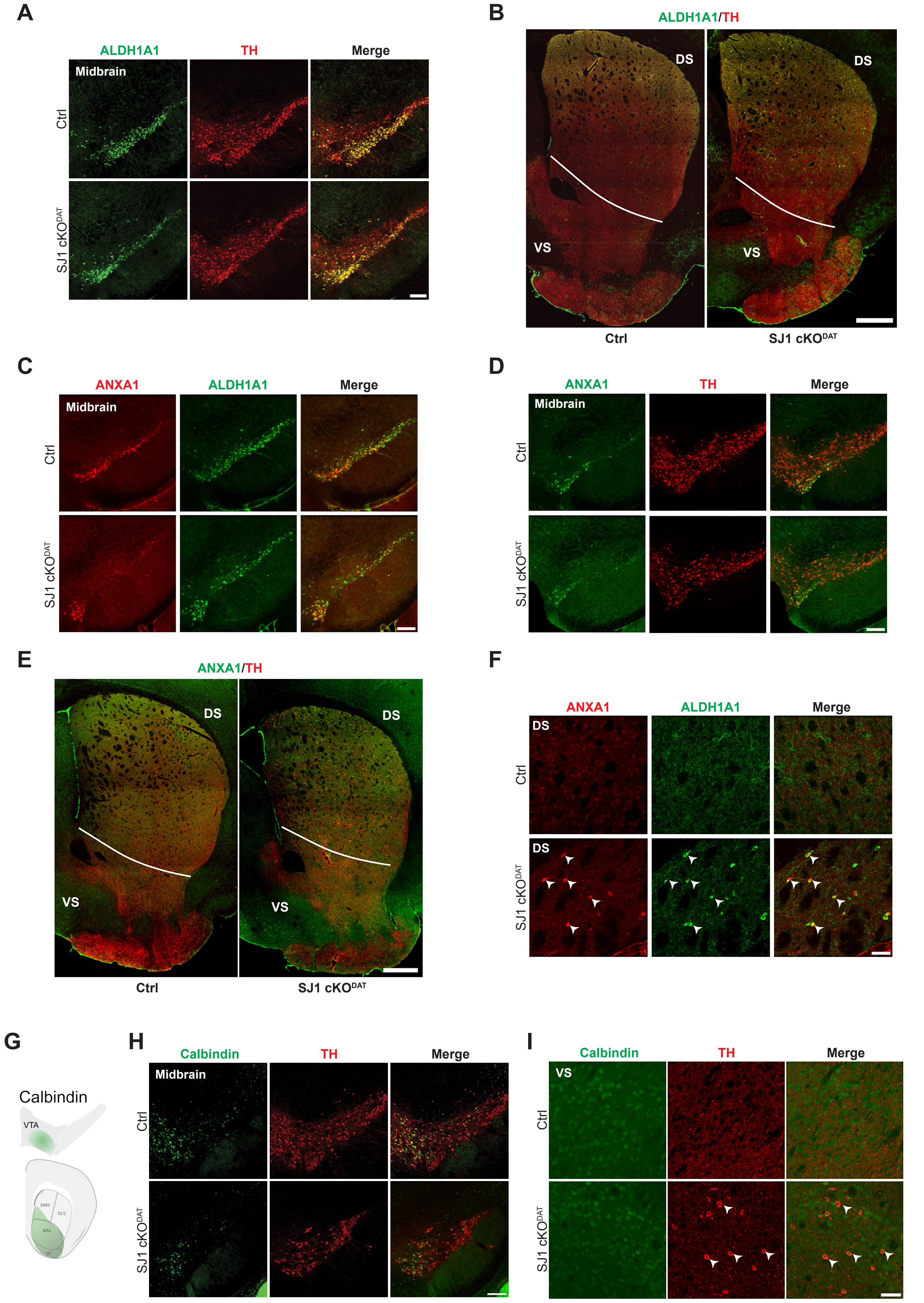


### Supplementary Figure 6

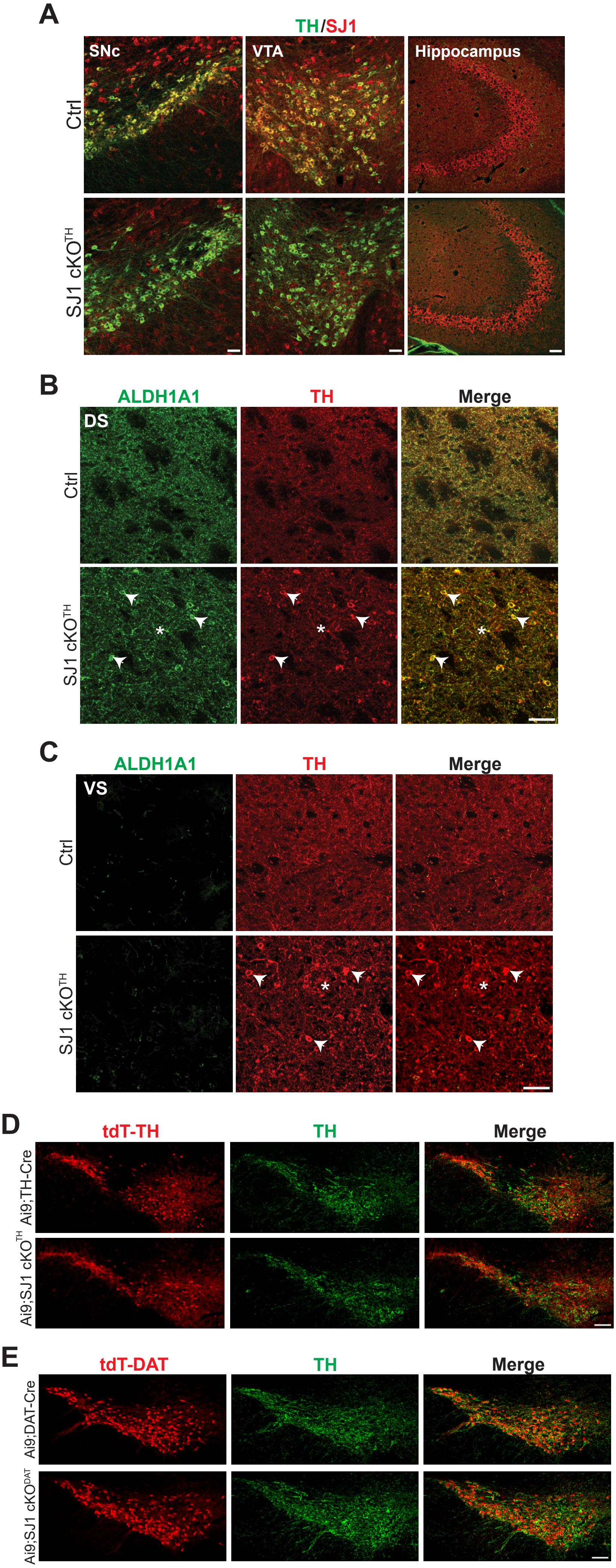


**Supplementary Figure 7**

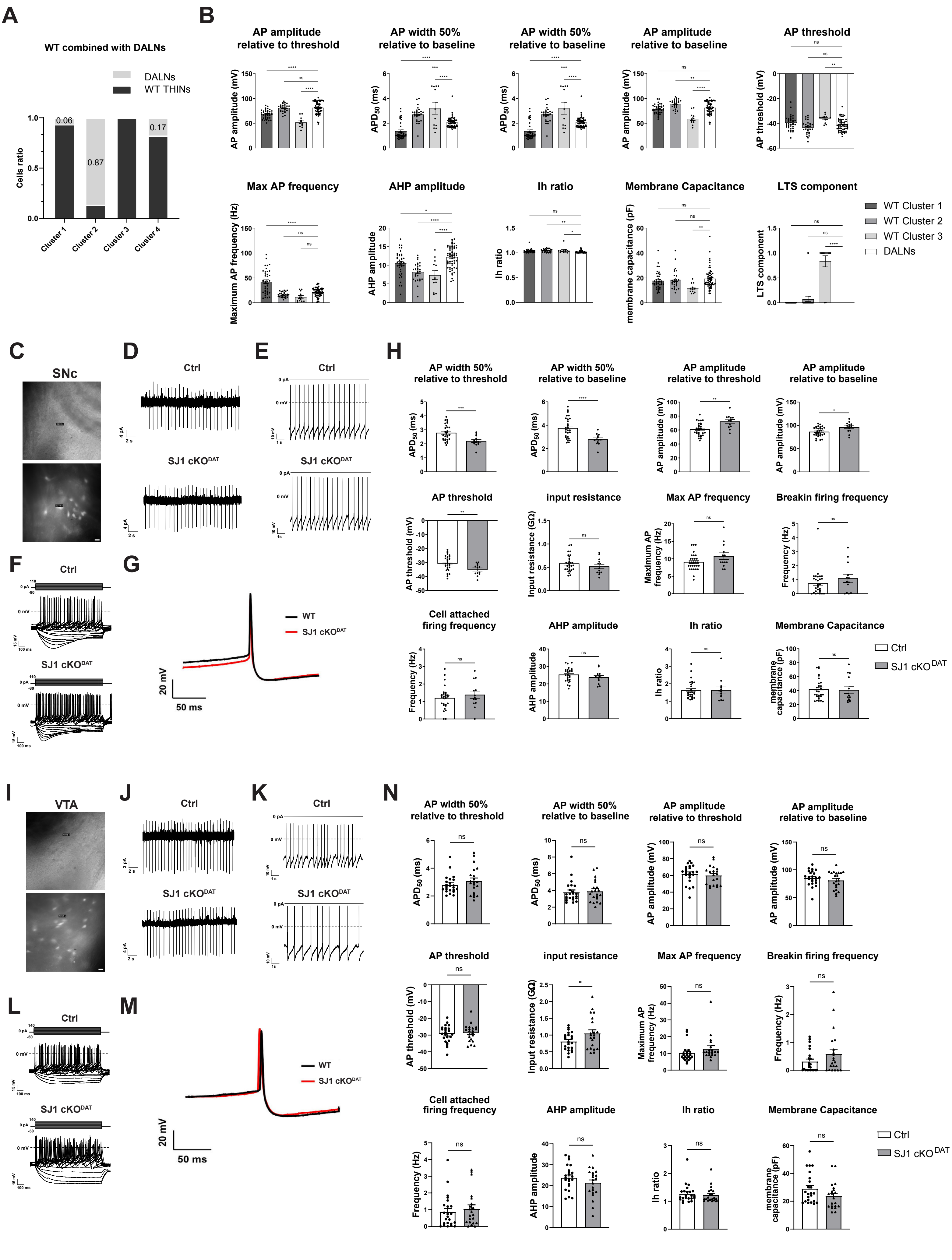


### Supplementary Figure 8

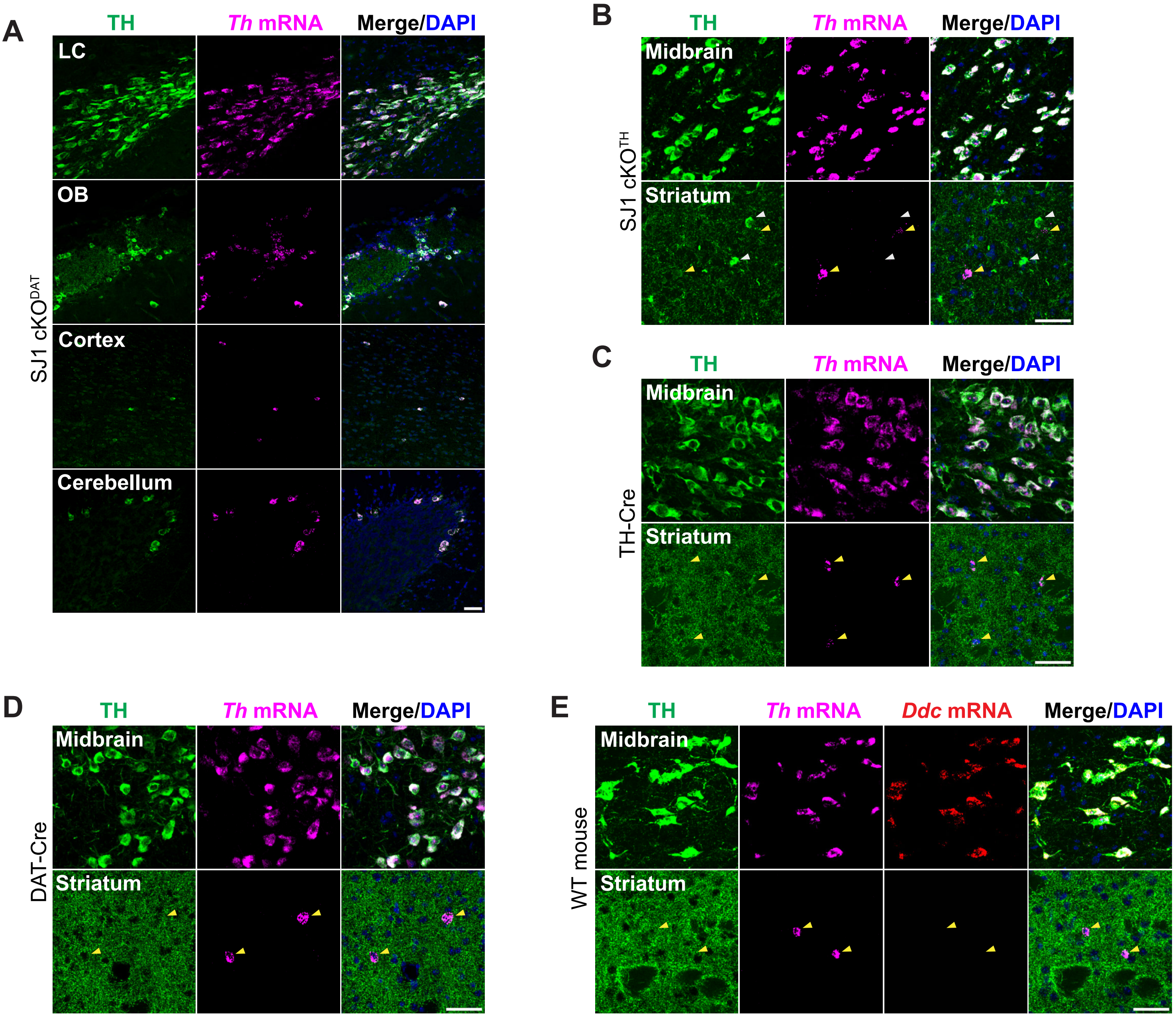


**Supplementary Figure 9**

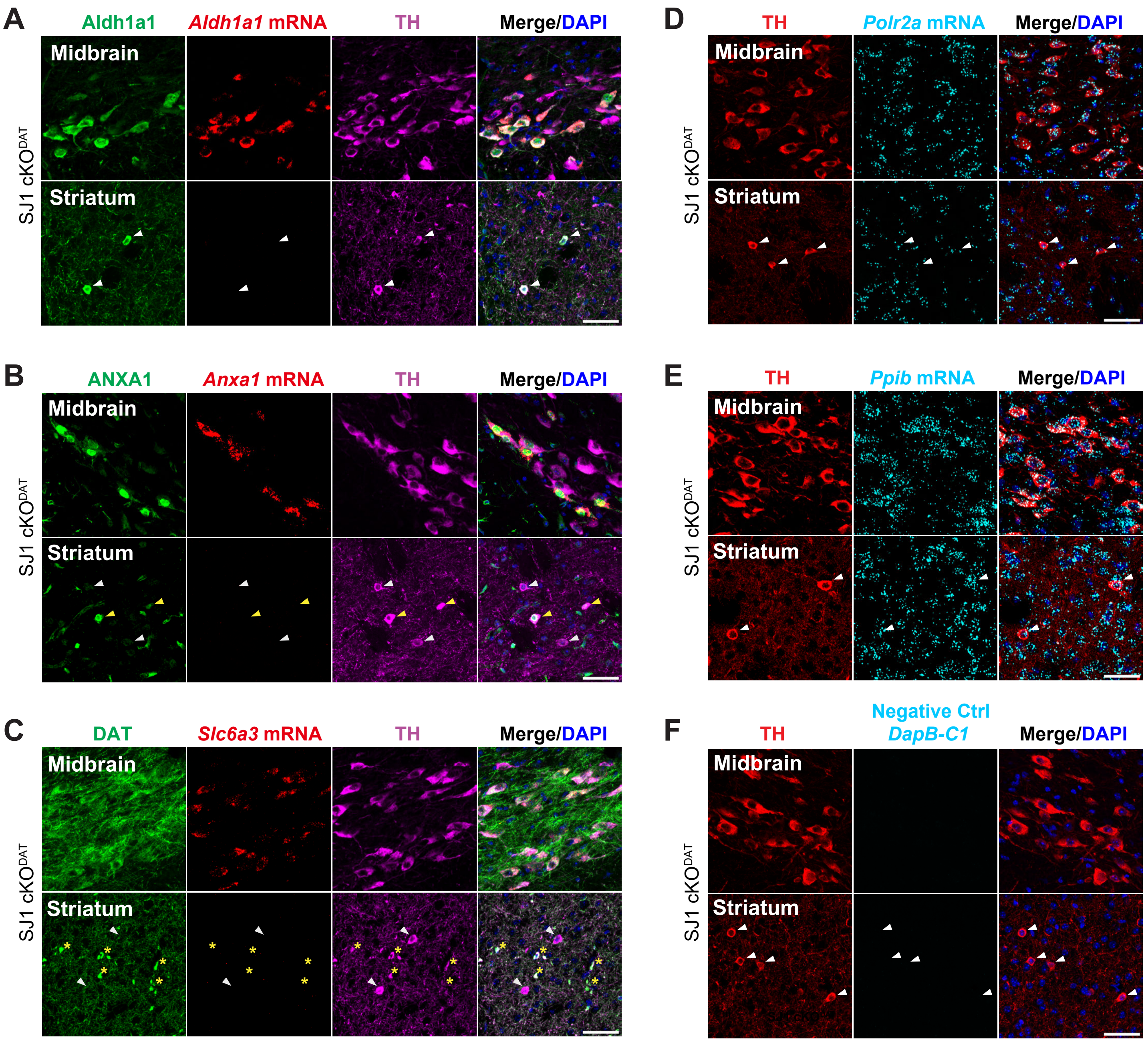


**Supplementary Figure 10**

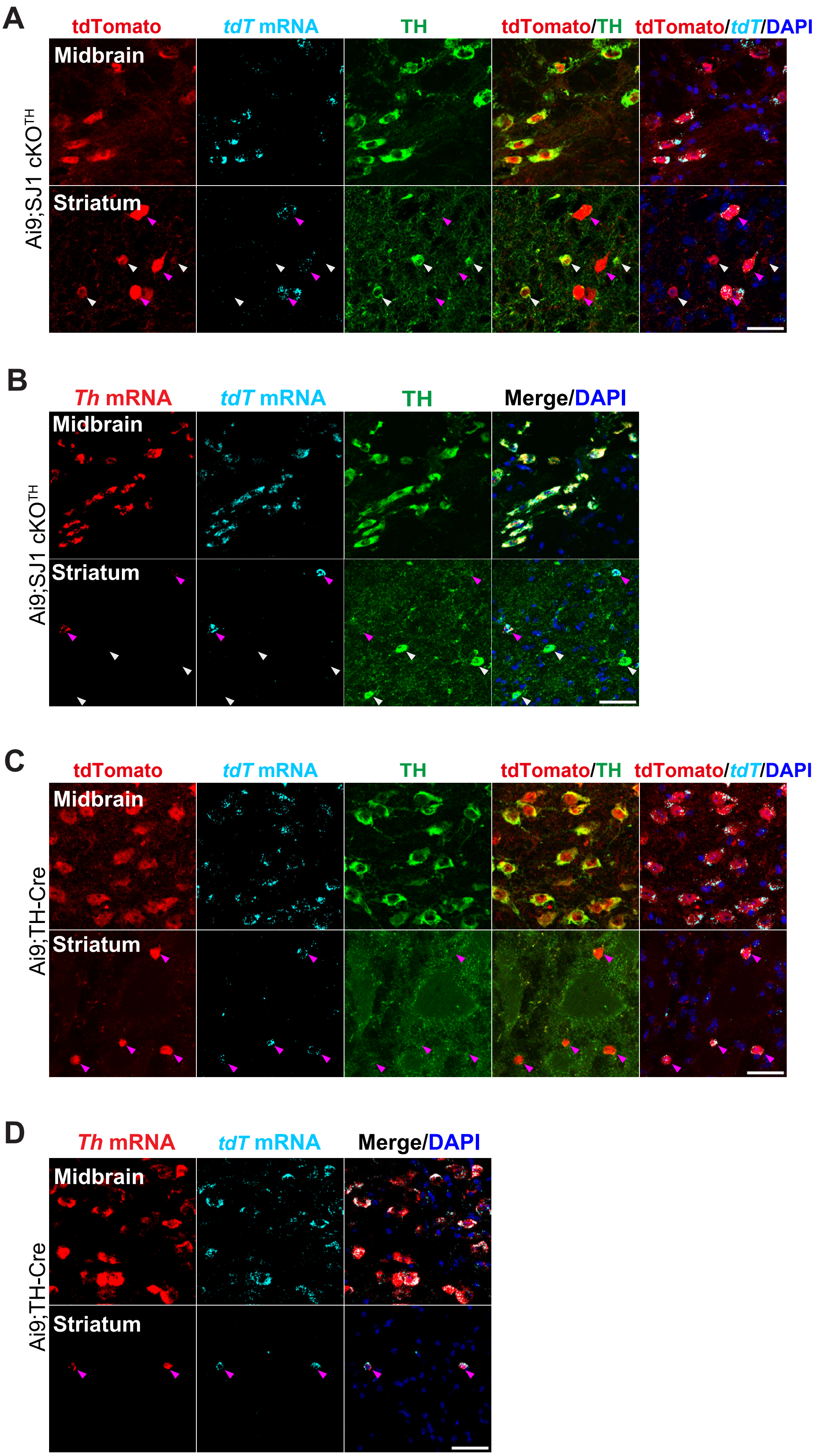


Supplementary Figure 11

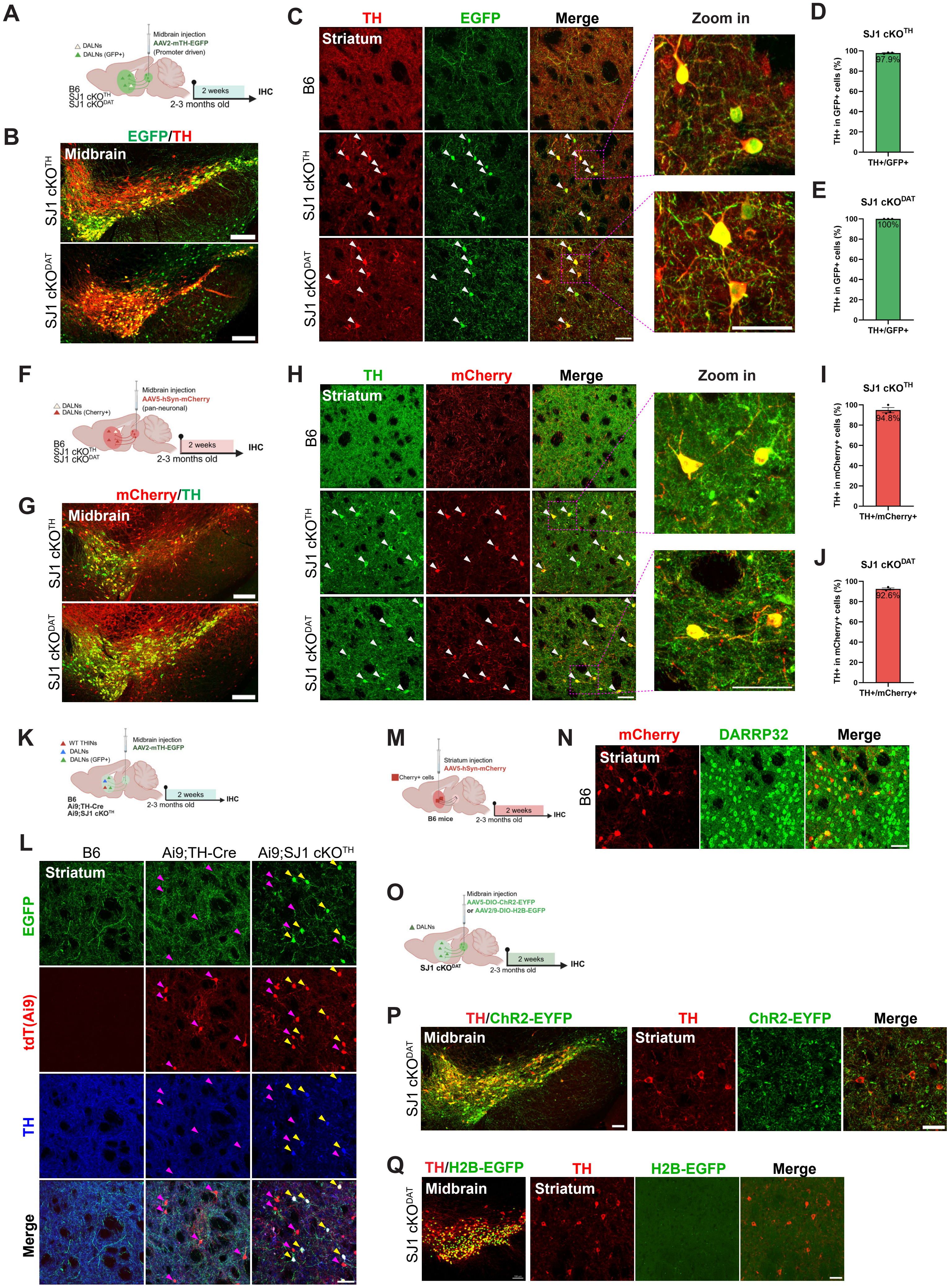


**Supplementary Figure 12**

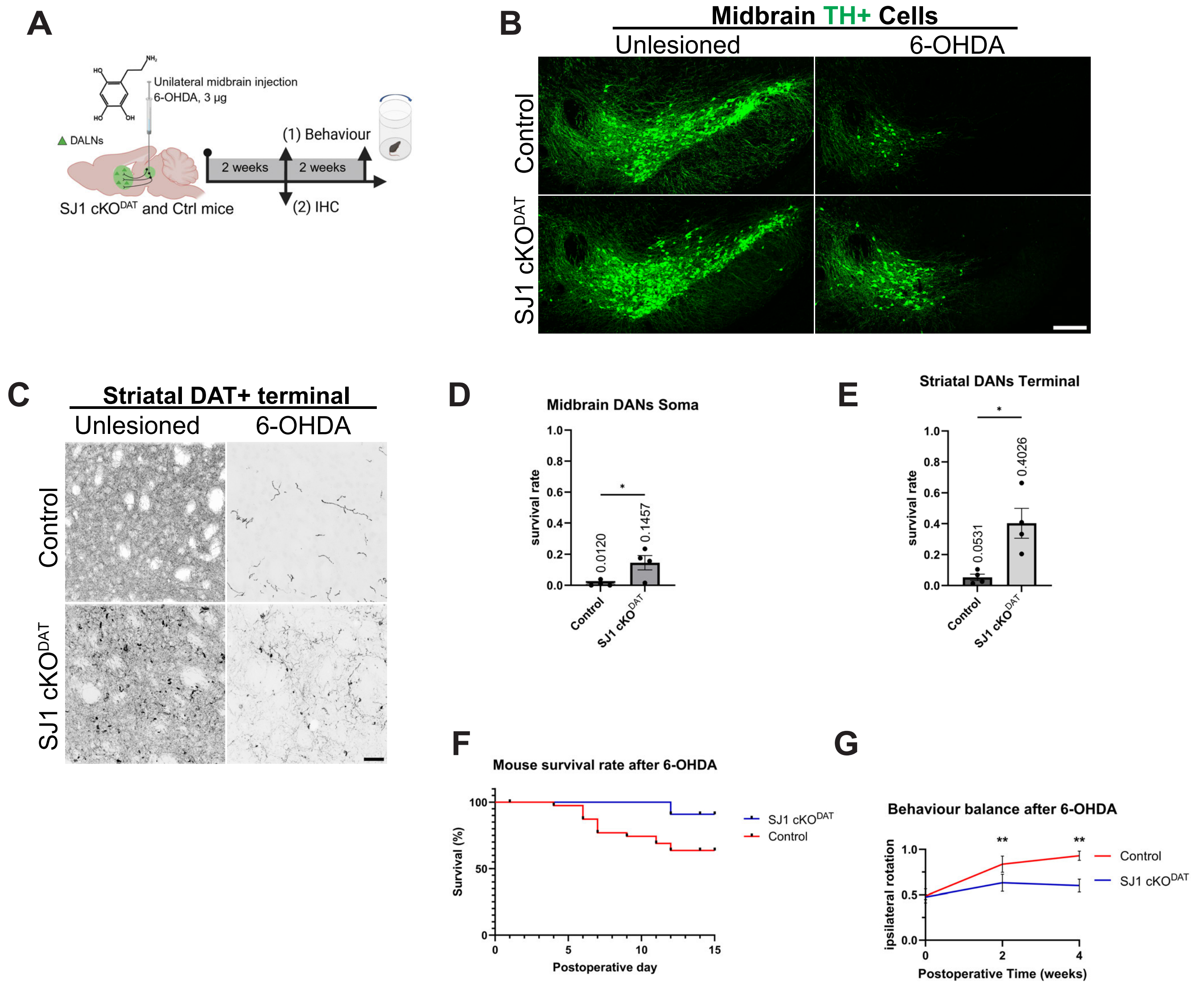


**Supplementary Figure 13**

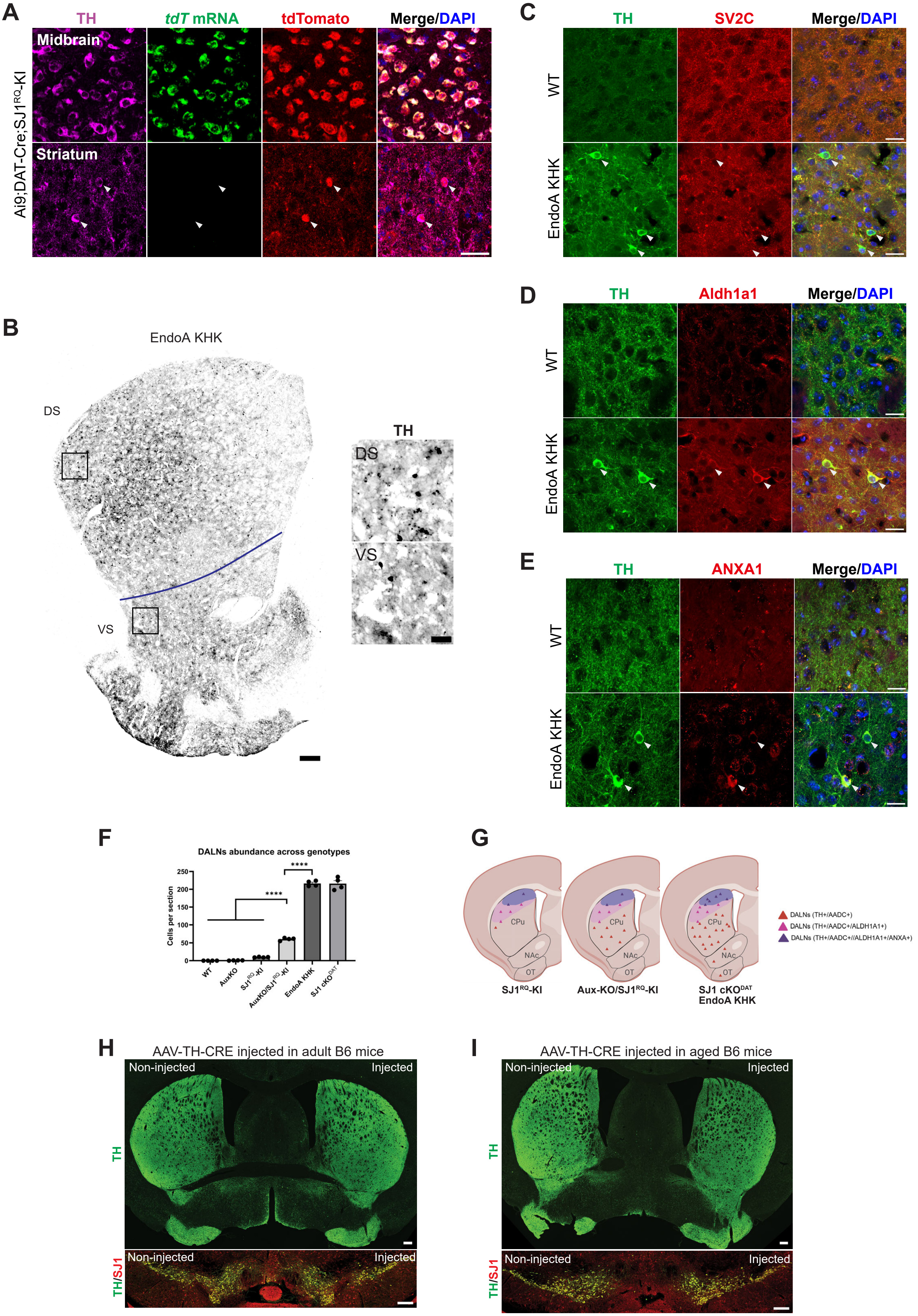


**Supplementary Figure 14**

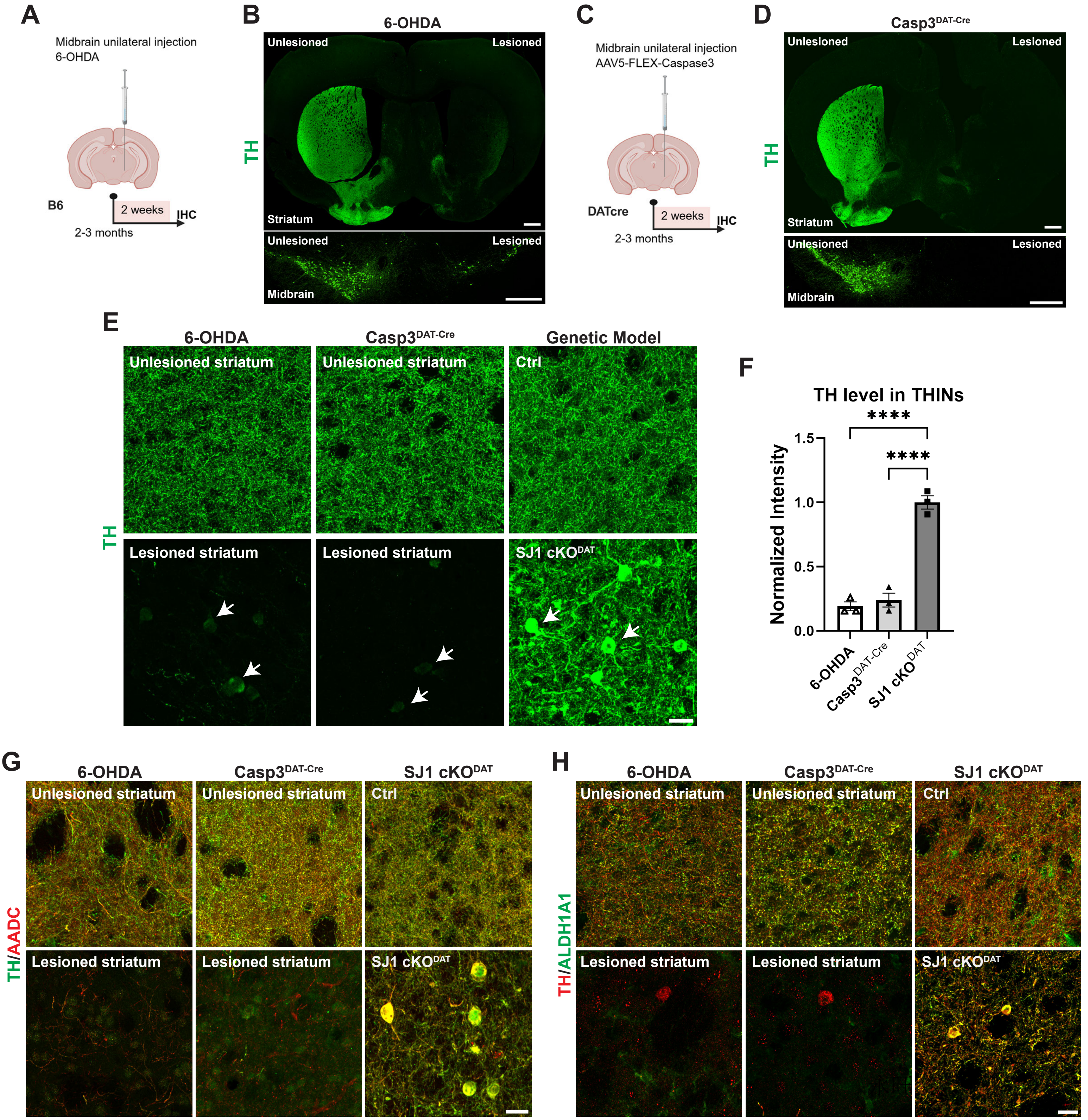


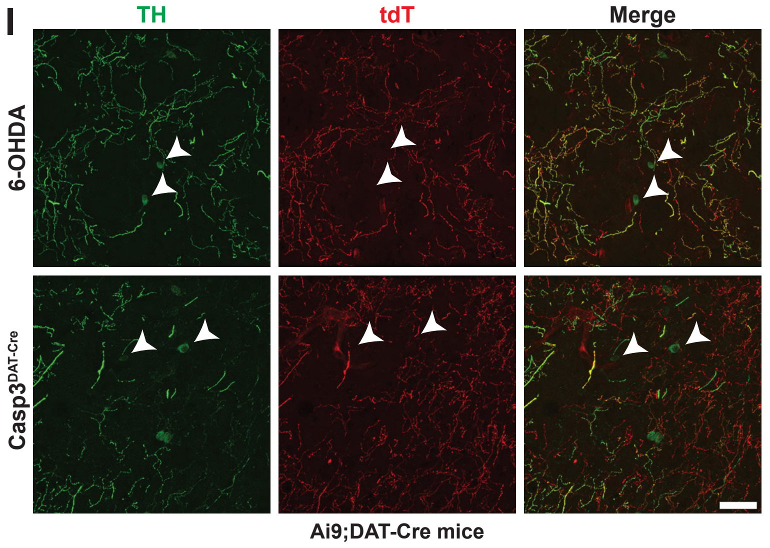


**J**

| Mouse Model | | TH | AADC | ALDH1A1/Anxa1 | DAT | tdT-TH | tdT-DAT |
|---|---|---|---|---|---|---|---|
| **Ai9;TH-Cre** | WT THINs | - | - | - | - | +++ | |
| **SJ1 cKO$^{DAT}$** | DALNs | +++ | +++ | +++ (*dorsal striatum) | - | | +++ |
| **SJ1 cKO$^{TH}$** | DALNs | +++ | +++ | +++ (*dorsal striatum) | - | ++ | |
| **6-OHDA model** | | + | - | - | - | - | |
| **Casp3 model** | | + | - | - | - | - | |

+++ positive and high expression;
++ positive and moderate expression;
+ positive and mild expression;
- negative/cannot be detected with immunostaining;
*remarks: ALDH1A1 largely marks dorsal region of the striatum, Anxa1 marks dorsal-lateral subregion within ALDH1A1

#### Supplementary Figure 15
